# The emergence and neural organisation of facial-expression information during ultra-brief vision

**DOI:** 10.64898/2026.11.17.767142

**Authors:** Renzo C. Lanfranco, Arvid Guterstam, Hugh Rabagliati, Axel Cleeremans, David Carmel, Andrés Canales-Johnson

## Abstract

A glimpse lasting only a few milliseconds can convey meaningful information about a face, but how the brain transforms such minimal input into stable representations of facial expression and conscious experience remains unclear. Across two experiments using ultra-brief face presentations, we combined hierarchical psychophysical modelling, multivariate EEG decoding, and information-theoretic measures to characterise this transformation. Extending previously reported findings, hierarchical modelling quantified distinct exposure-dependent trajectories for intact-face localisation and expression identification, providing a behavioural calibration of weak, intermediate, and stronger perceptual evidence. The critical neural transition occurred between 4.4 and 6.2 ms. At 4.4 ms, expression sensitivity was behaviourally detectable, but neural expression patterns did not reliably generalise to faces excluded from classifier training. At 6.2 ms, previously observed emotion-related ERP differences were accompanied by expression representations that generalised across faces and locally detectable emotion information during the early posterior negativity (EPN) interval. Subjective clarity tracked location-related neural evidence more consistently than expression-specific evidence, indicating that awareness did not simply mirror the emerging expression representation. The degree to which early centroparietal activity statistically predicted later occipital activity differed between emotional and neutral faces as exposure increased, indicating that the relationship between successive neural responses was reorganised. Thus, a few additional milliseconds of visual input did not merely strengthen facial-expression processing; they transformed weak evidence that did not reliably generalise across faces into more stable, generalisable, behaviourally relevant, and distributed neural information.

**SIGNIFICANCE STATEMENT:** A few milliseconds of visual input can support face perception, but it remains unclear whether additional input simply strengthens an emerging facial-expression representation or reorganises expression information across neural processing stages. Across exposures of 0.8–6.2 ms, previous analyses of these experiments established that face localisation and awareness precede reliable expression processing. The present information-theoretic analyses reveal that local emotion information, information shared between successive neural responses, predictive relationships across processing stages, and subjective clarity follow different exposure-dependent trajectories. These dissociations challenge simple accounts in which facial-expression information becomes uniformly available at a single perceptual threshold. Instead, they support a multistage account in which increasing sensory evidence progressively reorganises neural information and its availability to behaviour and awareness.

## INTRODUCTION

Facial expressions convey others’ affective states, intentions, and potential actions. Their perception recruits distributed neural systems that evaluate visual input for social and biological significance [1, 2]. Backward-masking and affective-priming studies show that expression information can influence behaviour, physiology, and neural activity when input or awareness is limited [3–6]. Our complementary minimal-exposure approach varies unmasked face duration directly from 0.8 to 6.2 ms [7–9], tracing expression processing as small increments of visual evidence become available. Previous work has established how little exposure supports expression processing, but not whether additional input strengthens the same representation, makes it more general across faces, or reorganises it across successive neural responses.

Emotional-face processing is a promising domain for studying this because it produces well-characterised time-resolved EEG responses. The posterior P1 around 100 ms post-stimulus onset is associated with early sensory encoding and attention. The EPN is a posterior negativity at 170–300 ms associated with preferential encoding of emotional stimuli, whereas the LPP is a sustained centroparietal positivity from approximately 300 ms associated with motivational evaluation [10–15]. Together, they span early visual processing, emotion-selective encoding, and sustained evaluation, but effects in several components do not reveal how information is organised among them. Signals may carry overlapping information, contribute complementary information jointly, or show directed predictive dependence. This distinction also matters for theories associating awareness with recurrent or globally distributed processing [16, 17], which motivate separating locally detectable information from broader patterns of neural dependence.

A second unresolved question concerns how these changing neural representations relate to subjective experience. Objective discrimination, neural differentiation, and subjective clarity need not develop together. The Perceptual Awareness Scale (PAS) measures graded subjective clarity [18, 19], whereas signal-detection methods quantify how accurately those reports distinguish correct from incorrect perceptual decisions [20, 21]. If awareness reflects access specifically to facial-expression information, PAS should track expression-related neural evidence. Alternatively, it may primarily reflect the location or general visibility of the stimulus. We therefore compared these possibilities trial by trial.

The experiments were previously reported as Experiments 1 and 5 of Lanfranco et al. [8], referred to here as Experiments 1 and 2. Experiment 1 sampled seven exposure durations from 0.8 to 6.2 ms and manipulated face orientation and expression; Experiment 2 recorded EEG at 1.7, 4.4, and 6.2 ms. The original analyses showed that intactface localisation and PAS-based metacognitive sensitivity required less exposure than expression-specific processing. Expression discrimination and multivariate expression decoding became detectable at 4.4 ms, whereas stronger, more sustained decoding and emotion-related EPN and LPP differences emerged at 6.2 ms. These findings established when successive markers became detectable, but not their psychometric emergence, representational stability across faces, organisation of emotion information across successive neural responses, or whether subjective clarity specifically tracked expression-related neural evidence.

Here, hierarchical psychometric and signal-detection models quantified objective and metacognitive sensitivity across exposure durations. Multivariate decoding tested the strength and temporal stability of location and expression representations, including their generalisation across face identities; trial-level analyses related neural evidence to objective accuracy and PAS. Mutual information assessed locally available emotion information, co-information characterised net redundancy- or synergy-dominant relationships with emotional status, and transfer entropy tested directed predictive dependence among P1-, EPN-, and LPP-related sensor patterns [22–24]. We expected objective sensitivity, perceptual clarity, and neural expression information to increase with exposure. Critically, we asked whether these changes occurred in parallel or whether additional visual input reorganised how emotion-related information was represented, distributed across processing stages, and related to perceptual experience.

## METHODS

### Study design

The present study integrated two complementary experiments to characterise how facial-expression information emerges, becomes distributed across neural processing stages, and relates to perceptual awareness during ultra-brief vision. Both experiments were initially reported by Lanfranco et al. [8], but the present study addresses distinct questions using a unified set of hierarchical psychophysical, electrophysiological, multivariate, and information-theoretic analyses. Experiment 1 corresponds to Experiment 1 of the original publication. Experiment 2 corresponds to the original Experiment 5 and is renamed here to reflect its position in the present study.

Experiment 1 provided a densely sampled behavioural characterisation across seven exposure durations from 0.8 to 6.2 ms and included both upright and inverted faces. Experiment 2 concentrated on three theoretically informative durations, 1.7, 4.4, and 6.2 ms, while EEG was recorded. These durations sampled conditions providing weak, intermediate, and stronger evidence for intact-face localisation and expression processing. Experiment 1 was used to estimate the psychometric development of objective and metacognitive sensitivity, whereas Experiment 2 was used to examine the corresponding behavioural and neural information dynamics.

### Participants

Both experiments were approved by the ethics committee of the Faculty of Psychological Science and Education at the Université libre de Bruxelles, Belgium. Participants provided informed consent, had normal or corrected-to-normal vision, and reported no history of neurological or psychiatric disorders. Participants received €15 for Experiment 1 and €30 for Experiment 2.

Excluding pilot participants, data were available from 34 participants who completed Experiment 1. Two participants were excluded because at least 5% of their location and expression reports were missing, and one was excluded because location performance remained at chance across all seven exposure durations. The final Experiment 1 sample therefore comprised 31 participants (mean age = 22.7 years, SD = 2.8, range = 18–40; 16 female; 25 right-handed).

Behavioural and EEG data from Experiment 2 were available for 40 participants. As in Experiment 1, participants were excluded when at least 5% of the relevant location, expression, or PAS reports were missing. One participant exceeded the threshold for both location and expression reports, leaving 39 participants in the behavioural sample (mean age = 23.00 years, SD = 2.61, range = 18– 29; 24 female; all right-handed).

For the Experiment 2 EEG analyses, eight participants were excluded following preprocessing quality-control assessment, leaving 32 participants (mean age = 24.8 years, SD = 5.1, range = 19–29; 20 female; all right-handed). Exclusion criteria were more than 15% bad channels, more than 5% grossly artifactual trials during preliminary review, or more than 10% rejected epochs after independent-component analysis (ICA; see below). All participants retained for the EEG analyses also satisfied the behavioural inclusion criteria. Age, gender, and handedness were not included as predictors because the study had no hypotheses concerning these variables.

Participant inclusion and the analysis-specific samples are summarised in Supplementary Table S1.

### Apparatus and stimuli

Stimuli were presented using a custom LCD tachistoscope based on the design of Sperdin and colleagues [25]. The apparatus comprised vertical and horizontal LCD displays optically superimposed by a semipermeable mirror. A dedicated microcontroller controlled the display back-lights with a reported temporal precision of 0.002 *±* 0.001 ms. One display continuously presented fixation, stimulus placeholders, instructions, and response cues, whereas the second was illuminated only during the brief target presentation (Fig. 1a,b). Technical details are provided in the Supplementary Information accompanying Lanfranco et al. [8].

**Figure 1:**
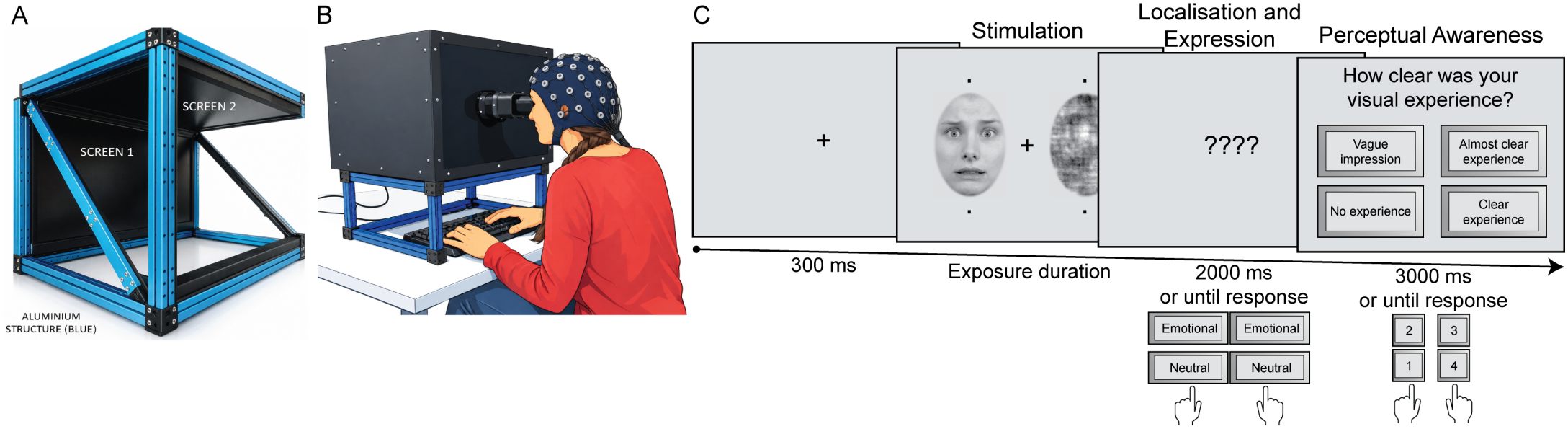
Experimental apparatus and trial sequence. **a**, Custom LCD tachistoscope comprising vertical and horizontal displays optically superimposed by a semipermeable mirror. One display continuously presented fixation, stimulus placeholders, instructions, and response cues, whereas the second was illuminated for the brief stimulus presentation. **b**, Participant performing the task while 64-channel EEG was recorded. The participant viewed the optically superimposed displays through the tachistoscope eyepiece and responded using the keyboard. **c**, Trial sequence. Following fixation and stimulus-location placeholders for 300 ms, an intact face and its scrambled counterpart were presented on opposite sides of fixation for 1.7, 4.4, or 6.2 ms. Participants then had up to 2,000 ms to make a compound response indicating both the location of the intact face and whether its expression was emotional or neutral. They subsequently had up to 3,000 ms to rate the clarity of their visual experience using the four-point PAS, ranging from “no experience” to “clear experience”. All face stimuli, including the intact face shown here, were taken from the Radboud Face Database (RaFD) and are presented as a stimulus example (see: https://rafd.nl/)

Face images were drawn from the Radboud Faces Database [26]. In Experiment 1, ten identities appeared with fearful and neutral expressions in both upright and inverted orientations. In Experiment 2, the stimulus set comprised 20 upright-face identities. Each identity appeared with a neutral expression and with one emotional expression; ten identities belonged to the fearful–neutral context and ten to the happy–neutral context. Identity sets and block order were counterbalanced across Experiment 2 participants. See Lanfranco et al. [8] for details on stimuli selection and standardisation.

In both experiments, an intact face appeared to the left or right of fixation while a scrambled version of the same image appeared at the mirror-symmetric location. The intact and scrambled images therefore contained closely matched low-level visual information, and successful localisation required distinguishing the intact configuration from its scramble.

### Task and procedure

Experiment 1 comprised 16 blocks of 70 trials following a 10-trial practice, producing 1,120 experimental trials. Stimuli were presented for 0.8, 1.7, 2.6, 3.5, 4.4, 5.3, or 6.2 ms. Exposure duration, upright or inverted orientation, fearful or neutral expression, and target side were balanced across the experiment.

Experiment 2 comprised 16 blocks of 105 trials following a 16-trial practice, producing 1,680 experimental trials. Eight blocks contained fearful and neutral faces and eight contained happy and neutral faces. Stimuli were presented for 1.7, 4.4, or 6.2 ms, with 560 trials at each duration and equal numbers of emotional and neutral trials.

The trial sequence was otherwise similar across experiments (Fig. 1c). Each trial began with fixation and stimulus-location placeholders for 300 ms, followed by the brief presentation of the intact face and its scrambled counterpart. Participants then had up to 2 s to make a single compound response indicating both the location of the intact face and whether its expression was emotional or neutral. They subsequently had up to 3 s to rate the clarity of their visual experience using the four-level PAS: 1, “no experience”; 2, “vague impression”; 3, “almost clear experience”; or 4, “clear experience”. The mappings between response keys and compound decisions for emotional-versus-neutral judgements, but not for left-versus-right judgements, and the spatial arrangement of the PAS alternatives were counterbalanced across participants.

### Behavioural analyses

#### Bayesian hierarchical type-1 and type-2 signal detection models

Location and expression decisions were analysed separately using hierarchical signal-detection models, with PAS ratings treated as ordered type-2 responses [20, 21]. Models were fitted separately for the two experiments. In Experiment 1, the location model represented the 28 combinations of seven exposure durations, two face orientations, and two expressions. The expression model represented the 14 combinations of exposure duration and orientation. In Experiment 2, the location model contained the three exposure durations, whereas the expression model contained the six combinations of exposure duration and emotional context, fearful–neutral or happy–neutral.

The models estimated condition-specific type-1 sensitivity (*d*^*′*^), response criterion, meta-*d*^*′*^, and metacognitive efficiency (M-ratio), while partially pooling participant-level variation. Location *d*^*′*^ and meta-*d*^*′*^ were divided by 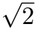 to express sensitivity according to the two-alternative forced-choice (2AFC) convention used in the original study. Because PAS measured the clarity of perceptual experience rather than conventional decision confidence, meta-*d*^*′*^ and M-ratio were interpreted as indices of the correspondence between subjective awareness and objective perceptual accuracy.

The Experiment 1 models included 31 participants, and the Experiment 2 models included 39 participants. Models were fitted using weakly regularising priors. Four Markov chain Monte Carlo (MCMC) chains were run, each with 2,000 warm-up iterations followed by 4,000 sampling iterations (*δ*_adapt_ = .99; maximum tree depth = 15). Estimates and planned contrasts are reported as posterior medians with 95% credible intervals. Convergence was considered satisfactory when the maximum 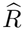 was *≤*1.01, the minimum bulk effective sample size was *≥*400, and no divergent transitions or maximum-tree-depth exceedances occurred. All models met these criteria. Full model specifications and prior distributions are reported in the Supplementary Methods and Supplementary Table S2. Convergence and sampling diagnostics are reported in the Supplementary Results and Supplementary Table S3, with representative Experiment 1 trace plots shown in Supplementary Fig. S1.

### Hierarchical psychometric-function models for Experiment 1

The densely sampled data from Experiment 1 were also analysed using hierarchical cumulative-Gaussian psychometric functions. Location accuracy was modelled separately for upright-neutral, upright-fearful, inverted-neutral, and inverted-fearful faces. Expression accuracy was modelled separately for upright and inverted faces. Chance performance was fixed at .50, and the upper asymptote included a participant-varying lapse rate constrained to be less than .15. Participant-specific intercepts and positive slopes were partially pooled within each condition. For computational efficiency, trials sharing the same participant, condition, and exposure duration were aggregated into binomial counts, which yields the same likelihood as the corresponding trial-level Bernoulli observations.

The primary psychometric outcome was ED_75_, defined as the exposure duration at which expected accuracy reached 75% after allowing for lapse. Planned contrasts tested whether ED_75_ differed between upright and inverted faces and, for location judgements, between fearful and neutral faces. Estimates outside the observed range of 0.8–6.2 ms were retained but interpreted as model-based extrapolations rather than directly observed thresholds.

Psychometric models used the same four-chain sampling schedule and diagnostic criteria as the signal-detection models. Accuracy curves, ED_75_ estimates, and planned contrasts are reported as posterior medians with 95% credible intervals. Prior specifications are reported in the Supplementary Methods and Supplementary Table S4. MCMC diagnostics are reported in the Supplementary Results, Supplementary Table S3, and Supplementary Fig. S2.

### Information shared among behavioural reports in Experiment 2

As a behavioural counterpart to the EEG information analyses, discrete information-theoretic analyses quantified dependence among PAS, location accuracy, and expression accuracy across the three exposure durations in Experiment 2 [23, 24]. These analyses were restricted to Experiment 2 because their purpose was to characterise the relationships among behavioural reports under the same task design and exposure conditions used for the EEG analyses. Primary measures were mutual information between PAS and each accuracy outcome; mutual information between PAS and the joint pair of accuracy outcomes; conditional mutual information between PAS and one accuracy outcome given the other; and co-information among PAS, location accuracy, and expression accuracy.

For Experiment 2, probabilities were estimated separately for each participant and exposure duration using a Jeffreys Dirichlet prior of 0.5 per cell. Information was expressed in bits. We generated 4,000 posterior draws per participant and used a Bayesian bootstrap to aggregate participants with equal expected weight. Targeted within-participant label permutations (2,000 permutations) supplied finite-sample null distributions. Mutual information and conditional mutual information used upper-tail permutation tests. co-information used a two-sided test centred on its targeted permutation-null mean and was reported as observed minus mean null; positive values indicate a redundancy-dominant balance and negative values a synergy-dominant balance under the co-information convention. The three exposure-specific tests for each metric formed a Holm-corrected family, and all three pairwise duration differences were obtained from paired posterior draws.

### Experiment 2 EEG event coding and trial definition

Trials were defined by the presentation of a face stimulus, followed by the location and expression discrimination responses and the PAS rating. Stimulus markers identified the facial expression (neutral, fearful, or happy), exposure duration (1.7, 4.4, or 6.2 ms), and side containing the intact face. Subsequent response markers jointly encoded the accuracy of the location and expression judgements, from which separate trial-wise accuracy measures and PAS ratings were derived.

Stimulus and response events were matched within trials and validated against the behavioural records. Each valid EEG epoch was annotated with its experimental condition, behavioural outcomes, PAS rating, stimulus identity, and trial identifier, allowing trials to be selected separately for each analysis while preserving their original acquisition order.

### Experiment 2 EEG acquisition and preprocessing

EEG was recorded at 512 Hz using a 64-channel BioSemi system and an elastic cap with electrodes positioned according to the international 10–20 system. Electrode offsets were maintained within the limits specified during acquisition.

Preprocessing was performed in MATLAB R2024b using EEGLAB 2026 [27], FieldTrip [28], and custom code.

Continuous data were resampled to 250 Hz and filtered between 0.3 and 40 Hz using a zero-phase finite-impulse-response filter. Filtering was performed before epoching to minimise edge artefacts at epoch boundaries. Electrode positions were defined using the standard BioSemi 64-channel montage.

Data were epoched from -200 to 600 ms relative to stimulus onset. No baseline correction was applied during preprocessing. Baseline correction was subsequently applied to the ERP data, whereas multivariate and information-theoretic analyses were conducted on unbaselined epochs. Channels with median-centred variance greater than 500 *µ*V^2^ or raw variance below 0.5 *µ*V^2^ were provisionally identified as bad and visually reviewed. To prevent global transients from inflating channel-variance estimates, epochs with median-centred variance greater than 5,000 *µ*V^2^ were provisionally omitted during channel assessment. Channels confirmed as bad were excluded from ICA estimation. ICA training additionally excluded epochs containing an absolute amplitude greater than 500 *µ*V in any retained channel or marked during the gross-variance review. These exclusions applied only to ICA estimation, and the affected epochs remained available for final post-ICA artefact assessment. Infomax ICA was estimated using EEGLAB’s algorithm [29] with a deterministic participant-specific random seed. The resulting decomposition was applied to all epochs over the channels included in the ICA fit.

Independent components were classified using the machine-learning-based ICLabel classifier [30]. Components assigned a probability of *≥*.70 to the Muscle, Eye, Heart, Line Noise, or Channel Noise category were removed. Channels identified as noisy during the preceding quality-control assessment were then reconstructed using spherical interpolation. Across the 32 included participants, 89 channel instances were interpolated, corresponding to a mean of 2.78 channels per participant (SD = 2.61, range = 0–9), or 4.35% of the 64 scalp electrodes (SD = 4.08%).

Following component removal and channel interpolation, residual artefacts were identified by an amplitude threshold of *±*100 *µ*V. In total, 1,344 of 53,755 epochs were rejected (2.50%). The participant-level proportion of rejected epochs averaged 2.50% (SD = 2.25%, range = 0.18%–8.63%), leaving 52,411 epochs for analysis. The retained data were re-referenced to the average of the 64 scalp electrodes.

### General EEG inferential framework

Mass-univariate EEG effects were evaluated using the nonparametric cluster-based permutation framework [31], with the family-wise error rate controlled at *α* = .05. Clusters were formed at a sample-level threshold of (*p* < .05), and cluster mass was defined as the sum of the test statistics within each cluster. Adjacency was defined temporally for one-dimensional time courses, across four directly neighbouring cells for time-by-time and time-by-lag maps, and using the FieldTrip [28] spatial-neighbourhood structure together with temporal adjacency for sensor-by-time ERP analyses.

Null distributions were generated from 5,000 within-participant permutations. For paired contrasts, participant-level difference scores were randomly signflipped. Only samples for which at least 12 participants contributed valid observations were tested. Pairwise tests were two-sided, whereas omnibus F-tests were evaluated in the upper tail of the permutation distribution.

When several maps constituted a planned family of tests, maximum-cluster-mass correction first controlled the family-wise error rate across samples within each map. The smallest cluster-corrected *p*-value in each map was then treated as its map-level probability, and these values were adjusted across the planned family using the Holm procedure. Differences between exposure durations were assessed through direct contrasts of the corresponding duration-specific effects.

### Event-related potentials

For ERP analyses, each retained trial was baseline-corrected by subtracting its mean voltage from -200 to 0 ms. Spatial neighbours were defined from the BioSemi sensor geometry using triangulation. ERP tests used 10,000 Monte Carlo randomisations and required at least two neighbouring electrodes to form a spatial cluster. All other cluster-construction and inferential settings followed the general EEG inferential framework described above.

### Exposure-duration ERPs

To characterise general visual-evoked activity, all retained epochs were averaged separately for the 1.7-, 4.4-, and 6.2-ms conditions. An all-electrode repeated-measures cluster *F* -test evaluated the three-level exposure effect. Follow-up paired, two-sided cluster tests compared 1.7 versus 4.4 ms, 4.4 versus 6.2 ms, and 1.7 versus 6.2 ms. The minimum cluster-corrected probability from each pairwise map was Holm-adjusted across the three planned contrasts.

### Emotion-related ERPs

To examine emotion-related ERP activity, fearful and happy trials were pooled into a single emotional condition and compared with neutral trials separately at each exposure duration. All artefact-free trials were included irrespective of location or expression discrimination accuracy. Because neutral stimuli occurred twice as frequently as each individual emotional expression, pooling fearful and happy trials produced closely matched emotional and neutral conditions, and trial subsampling was therefore unnecessary. At 1.7 ms, participants contributed a mean of 272.13 neutral trials (SD = 7.39) and 272.78 emotional trials (SD = 5.97); the corresponding means were 273.56 (SD = 7.20) and 273.25 (SD = 6.41) at 4.4 ms, and 273.41 (SD = 6.62) and 272.72 (SD = 6.67) at 6.2 ms. Across participants and durations, the analysis included 26,211 neutral and 26,200 emotional trials.

Emotional and neutral ERPs were compared at each exposure duration using paired, two-sided sensor-by-time cluster-based permutation tests encompassing all scalp electrodes. The three duration-specific tests constituted a single planned family, and their map-level probabilities were adjusted using the Holm procedure.

### Time-resolved multivariate pattern analysis

Multivariate analyses used the unbaselined, average-referenced single-trial data from all 64 electrodes. Analyses were performed separately at each exposure duration using MVPA-Light [32] and custom MATLAB code. The classifier was regularised linear discriminant analysis (LDA) with analytic shrinkage (*λ* = auto). Classes were balanced by randomly undersampling the training data only, and feature standardisation was estimated on the training fold and applied to the held-out fold.

### Expression and location decoding

The primary expression analysis decoded three classes: neutral, fearful, and happy, at every time sample. A two-class left-versus-right location decoder was run in parallel as a task-relevant control. The primary validation used 10-fold cross-validation repeated five times, with all trials from a face identity assigned to the same fold so that testing was performed on unseen identities. For comparability with the original report, a secondary analysis used stratified 10-fold trial-wise cross-validation. Performance was quantified using macro-averaged receiver-operating-characteristic area under the curve (AUC; chance = .50) and balanced accuracy (chance = 1/3 for expression and 1/2 for location). Macro-averaged AUC was the primary performance measure, whereas balanced accuracy was reported as a complementary measure.

Group inference compared participant performance with chance using sign-flip cluster tests. One-dimensional decoding curves used 5,000 permutations, and duration-specific curves were Holm-adjusted within each analysis family. Identity-generalising expression decoding was additionally compared directly between 6.2 and 4.4 ms using a paired, two-sided temporal cluster-mass sign-flip test with 5,000 permutations. Because this was a single planned duration contrast, no additional Holm adjustment across duration pairs was applied.

### PAS modulation of decoded neural evidence

To determine when whole-scalp task representations covaried with subjective perceptual clarity, PAS ratings were related to time-resolved, cross-validated neural evidence. PAS was not used during classifier training. Instead, the expression and location decoders returned a true-class discriminant margin for each held-out trial, with larger values indicating stronger neural evidence for the expression or target location presented on that trial. These margins were standardised at each time point within participant and exposure duration.

At each time point, standardised neural evidence was modelled as a function of mean-centred PAS, with presented expression, side of the intact face, experimental block, and face identity included as nuisance variables. The resulting PAS coefficient therefore represented the change in neural evidence, in standard-deviation units, associated with a one-level increase in PAS after accounting for these experimental factors. Estimation required at least 30 valid trials and two distinct PAS ratings within each participant and exposure duration. All participant-by-duration cells met these criteria. The mean numbers of eligible trials were 543.63 at 1.7 ms (SD = 14.39, range = 504–560), 545.56 at 4.4 ms (SD = 14.40, range = 502–560), and 544.69 at 6.2 ms (SD = 13.77, range = 508–559).

PAS slopes were estimated separately for expression and location evidence. The expression-minus-location slope provided a specificity test of whether PAS was more strongly associated with expression information than with general spatial or stimulus information. Exposure-duration effects on the expression-evidence slope were evaluated through direct paired contrasts between 6.2 and 4.4 ms, 4.4 and 1.7 ms, and 6.2 and 1.7 ms.

### Sensor-network information analyses

Information analyses used the unbaselined clean epochs and the Gaussian-copula mutual-information estimator (GCMI; 23). The primary signal representation was scalp current density (CSD), calculated using FieldTrip’s spherical-spline method [28] to attenuate broadly distributed, volume-conducted fields. Because CSD reduces but does not eliminate field spread, all connectivity-like quantities were interpreted as sensor-level statistical dependencies rather than anatomical connections or causal information transfer.

### Sensor definitions and trial matching

Sensor definitions were based on the topographies reported by Lanfranco et al. [8], fixed before the network tests, and not selected from statistical peaks in the present analyses. The left-posterior grouping comprised P7, P9, PO7, O1, P5, and PO3, and the right-posterior grouping comprised P8, P10, PO8, O2, P6, and PO4. Their union defined the bilateral posterior EPN node used in the EPN–LPP model. The centroparietal node comprised CPz, Pz, CP1, CP2, P1, and P2; Cz was additionally included in multichannel secondary analyses. A compact occipital node centred on the P1 topography comprised O1, Oz, and O2. For the P1–EPN analyses, O1 and O2 were excluded from the EPN node so that the two nodes did not share sensors, leaving P7, P9, PO7, P5, PO3, P8, P10, PO8, P6, and PO4.

For analyses comparing emotional and neutral expressions, fearful–neutral and happy–neutral trials were matched separately within face identity and target side. Within each stratum, the retained number was the minimum available across emotional status and all three exposure durations. Participants were required to contribute at least 80 matched pairs per emotional expression and exposure duration. When trial matching required random subsampling, a fixed random seed ensured that the same trials were selected each time the analysis was run.

### Information measures

We followed our extensive prior work on information-theoretic analyses of electrophysiological signals [33–38]. In brief, mutual information quantified the information that the signal at an individual node carried about the variable of interest and was expressed in bits:

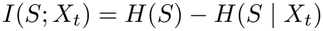

where H denotes entropy, S denotes the variable of interest (emotional status or, in the local PAS analyses, PAS) and X_t_ denotes the signal at a sensor node at time t. Mutual information therefore represents the reduction in uncertainty about S provided by the neural signal.

Co-information quantified the net redundancy–synergy balance in information about emotional status jointly represented by two sensor nodes:

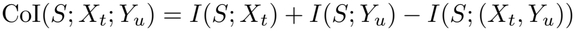

where S denotes emotional versus neutral status, and X_t_ and Y_u_ denote activity at the two nodes at times t and u, respectively. Positive values indicate a net redundancy-dominant balance, whereas negative values indicate a net synergy-dominant balance; they do not constitute separate estimates of redundant and synergistic information.

Directed predictive dependence was quantified using a first-order, single-lag Gaussian-copula transfer-entropy estimator:

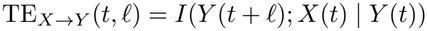

where activity in the source node, X_t_, predicted subsequent activity in the target node, Y_(t+l)_, after conditioning on activity in the target node at the source time, Y_t_. Thus, the estimator quantified conditional predictive dependence across trials.

Transfer entropy was estimated separately for fearful, happy, and their respective matched-neutral conditions. The fearful-minus-neutral and happy-minus-neutral effects were then averaged to obtain the pooled emotional-minus-neutral effect. Both spatial directions were evaluated. These estimates were interpreted as directed predictive dependencies between sensor signals and not necessarily as evidence of causal neural communication.

### Posterior–centroparietal EPN–LPP model

The posterior–centroparietal model examined information dynamics between the bilateral posterior EPN node and the centroparietal LPP node. Each node was represented by the equal-weight mean of its constituent sensors. GCMI quantified information about emotional versus neutral status at each node, with inferential testing restricted to 150–596 ms.

Co-information was evaluated across all posterior-by-centroparietal time pairs within 150–596 ms. Transfer entropy was evaluated in both posterior-to-centroparietal and centroparietal-to-posterior directions using lags from 8 to 120 ms in 4-ms steps. Emotion effects were tested separately at each exposure duration, and direct paired contrasts compared the three exposure durations. Co-information estimates were additionally evaluated relative to a paired-label permutation null.

Because zero-phase filtering can spread post-stimulus signals backwards in time, the interval from -180 to -4 ms was examined as a temporal-leakage diagnostic rather than as a confirmatory analysis interval. Full procedures for these analyses are reported in the Supplementary Methods.

### P1–EPN and P1–LPP models

The P1-network analyses examined whether early occipital activity was informationally related to subsequent activity in the EPN and LPP sensor groups. Theoryguided component intervals were fixed before conducting the information-theoretic tests, drawing on the observed ERP timing [8] and established temporal definitions of the P1, EPN, and LPP. Mutual information about emotional status was evaluated from 105 to 135 ms in the occipital P1 node, from 295 to 325 ms in the EPN node, and from 310 to 430 ms in the LPP node. Because the two network models used the same occipital node and P1 interval, P1 mutual information was estimated once and used in both models. Co-information and transfer entropy were evaluated across all combinations of P1-window and later-component time points, separately at each exposure duration and in direct pairwise exposure-duration contrasts.

Broader-window sensitivity analyses evaluated co-information and transfer entropy between 80–160 ms and 260–360 ms for the P1–EPN model, and between 80–160 ms and 280–500 ms for the P1–LPP model. Both spatial directions were tested. In the occipital-to-later-node direction, early occipital activity predicted subsequent EPN or centroparietal activity after conditioning on early activity at the target node. In the reverse spatial direction, early activity at the EPN or centroparietal sensor group predicted later occipital activity after conditioning on early occipital activity. The node labels therefore identify spatial sensor groupings and do not imply that the corresponding ERP component was present at both source and target times.

The 6.2-versus-4.4-ms P1–LPP transfer-entropy contrast was further examined using surrogate-centred estimates:

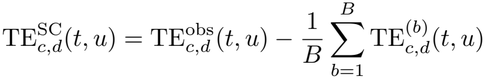

where *c* denotes expression condition, *d* denotes exposure duration, *t* and *u* denote the early source and later target times, *B* is the number of source-trial permutations, and 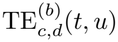 denotes the transfer-entropy estimate obtained from the *b*th permutation. The primary distributed-CSD analysis used B=200.

Surrogate estimates were generated by circularly shifting source trials within face-identity and target-side strata. This procedure preserved condition composition and marginal signal distributions while disrupting trial-specific correspondence between source and target activity. The two spatial directions and their interaction constituted one Holm-corrected inferential family. Sensitivity analyses examined alternative late intervals, sensor definitions, signal representations, expression-specific effects, and participant influence. Additional sensitivity and robustness procedures are described in the Supplementary Methods.

### Local PAS information

Complementing the decoding analysis, this analysis tested nonlinear dependence between subjective clarity and activity within the predefined posterior and centroparietal sensor groups. Trial-wise PAS ratings and node signals were residualised for presented expression, face identity, target side, block, and location accuracy before GCMI estimation. Mutual information between PAS and bilateral-posterior or centroparietal activity was evaluated on a 20-ms grid from 140 to 596 ms. Posterior and centroparietal PAS information were also compared directly for 4.4 versus 1.7 ms, 6.2 versus 4.4 ms, and 6.2 versus 1.7 ms. The six node-by-duration contrasts formed one Holm-corrected family.

### Trial-level neural evidence and behaviour

Complementing the time-resolved decoding analyses, this analysis tested whether trial-to-trial variation in neural evidence was related to objective accuracy and subjective perceptual clarity, and whether these relationships varied with exposure duration. Neural evidence was derived from two regional and temporal feature sets specified a priori, independently of the associations tested here. The posterior EPN feature comprised the 12 bilateral posterior sensors defined above, with activity averaged from 170 to 300 ms within each sensor. The centroparietal LPP feature comprised CPz, Pz, CP1, CP2, P1, P2, and Cz, with activity averaged from 300 to 596 ms. Each trial was therefore represented by a 12- or 7-element spatial pattern.

At each exposure duration, separate three-class expression and two-class location classifiers used the identity-grouped, repeated cross-validation and shrinkage-LDA procedure described above, applied separately to each feature set. The true-class discriminant margin, averaged across cross-validation repetitions, defined the neural evidence for the expression or target location presented on each held-out trial. Expression and location evidence were standardised separately within participant, exposure duration, and feature set. Feature-specific decoding performance was quantified using balanced accuracy and compared with chance using two-sided one-sample t-tests. For each decoding target, the six tests formed by the two feature sets and three exposure durations constituted one Holm-corrected family.

Expression accuracy was analysed using a logistic mixed-effects model containing the interaction between expression evidence and exposure duration, together with location evidence, presented expression, and target side. The corresponding location-accuracy model contained the interaction between location evidence and exposure duration, together with expression evidence, presented expression, and target side. Models initially included random intercepts for participant and face identity and a participant-specific slope for the focal neural-evidence predictor. When the data did not support estimation of the random slope, as indicated by singularity or nonconvergence, the model was reduced to participant- and identity-specific random intercepts. For each outcome and feature set, a joint Wald test evaluated the three exposure-specific evidence slopes. Individual exposure-specific slopes and pairwise differences between slopes were obtained from estimated marginal trends and are reported as odds ratios per standard-deviation increase in neural evidence.

PAS was analysed using separate cumulative-logit models for each participant, preserving its four-level ordinal scale while controlling for face identity and experimental block. Each model included the interactions of exposure duration with expression and location evidence, together with expression accuracy, location accuracy, presented expression, target side, face identity, and block. Exposure-specific participant slopes were evaluated at the group level using two-sided one-sample (t)-tests. For each feature set and evidence type, Hotelling’s (T^2^) test evaluated the joint vector of slopes across the three exposure durations.

Holm correction was applied separately within each outcome and evidence family. Each family comprised the two feature-set omnibus tests, the six feature-by-duration slopes, or the six feature-by-pairwise-duration contrasts, as appropriate.

## RESULTS

### Experiment 1 calibrated the behavioural progression across ultra-brief exposures

Experiment 1 provided a densely sampled behavioural calibration of the perceptual evidence available across seven exposure durations, including the three durations used for EEG in Experiment 2. Location accuracy increased steeply with exposure, reaching 75% at approximately 3–4 ms (Fig. 2a). Marginalising across expression, inversion delayed ED_75_ by 0.606 ms [0.216, 0.998], whereas the fearful-minus-neutral ED_75_ difference was not credible (0.126 ms [-0.260, 0.509]). Face orientation therefore affected the exposure required for intact-face localisation, whereas fearful expressions did not facilitate localisation, in line with our previous results [8].

**Figure 2:**
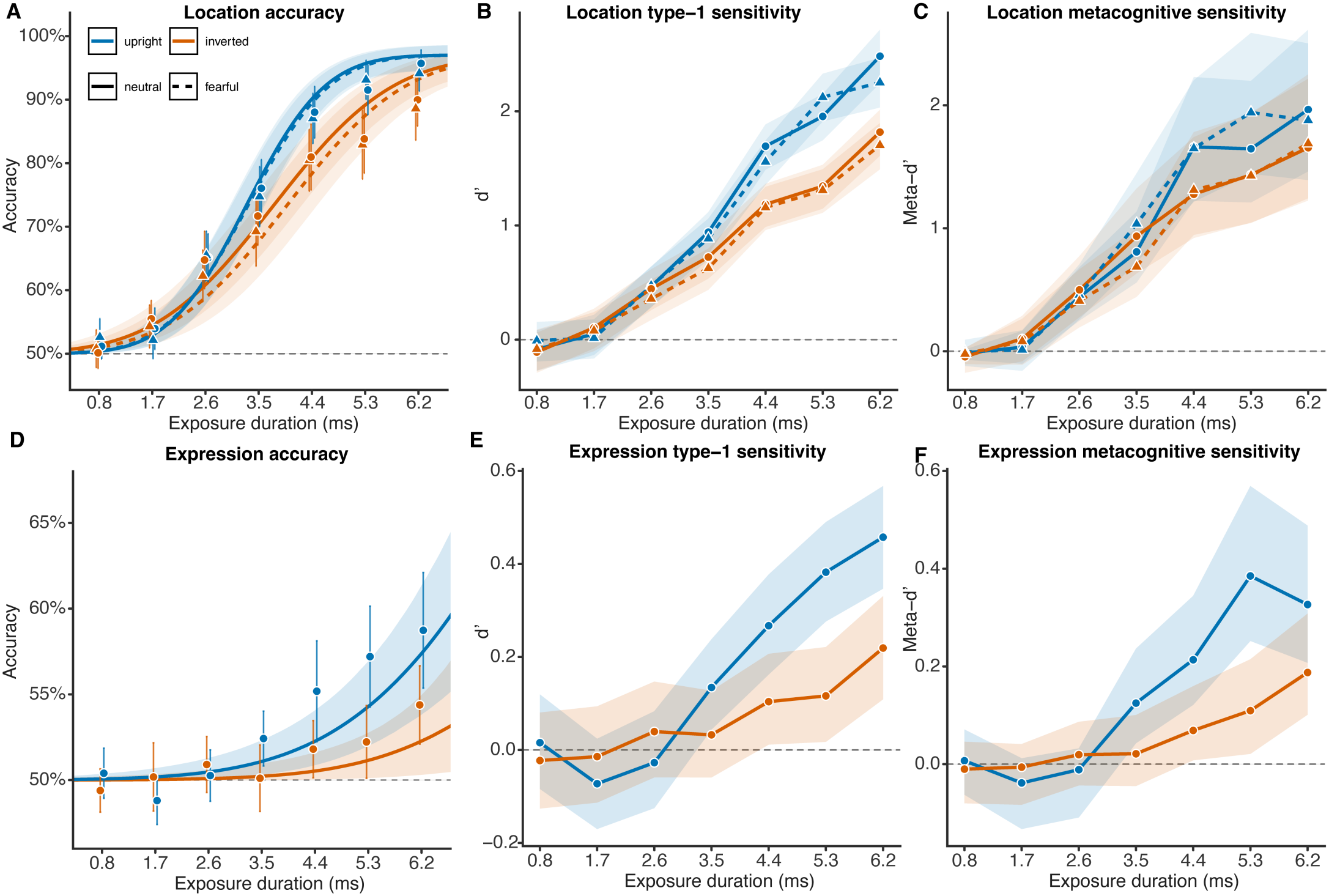
Psychometric and signal-detection characterisation of Experiment 1. **a**, Location accuracy across the seven exposure durations. Points show observed participant means with 95% confidence intervals; lines and shaded regions show posterior medians and 95% credible intervals from the hierarchical cumulative-Gaussian psychometric models. **b**, Bayesian hierarchical estimates of location type-1 sensitivity (*d*^*′*^). **c**, Corresponding estimates of PAS-based location metacognitive sensitivity (meta-*d*^*′*^). **d**, Observed expression accuracy and fitted psychometric functions. **e**, Bayesian hierarchical estimates of expression *d*^*′*^. **f**, Corresponding estimates of expression meta-*d*^*′*^. Blue denotes upright faces and orange denotes inverted faces. In the location panels, solid lines and circles denote neutral faces, whereas dashed lines and triangles denote fearful faces. Grey dashed lines indicate chance accuracy in a and d and zero sensitivity in b, c, e, and f. Lines and shaded regions in the signal-detection panels show posterior medians and 95% credible intervals.

The hierarchical signal-detection estimates supported this pattern (Fig. 2b,c). At 1.7 ms, the credible intervals for location *d*^*′*^ and meta-*d*^*′*^ included zero in all four orientation-by-expression conditions. Both measures subsequently increased steeply and were clearly positive in every condition by 4.4 ms, in line with our previous results [8]. At this duration, the upright-minus-inverted difference in location *d*^*′*^ was 0.509 [0.331, 0.690] for neutral faces and 0.399 [0.225, 0.574] for fearful faces. Corresponding differences in meta-*d*^*′*^ were 0.380 [0.162, 0.649] and 0.336 [0.122, 0.588]. The upright-face advantage persisted at the longer exposure durations, indicating that orientation affected both objective localisation and the correspondence between localisation accuracy and subjective experience. Metacognitive-efficiency estimates are reported in the Supplementary Results, Supplementary Fig. S3, and Supplementary Table S5.

Expression identification developed more gradually (Fig. 2d–f). Expression *d*^*′*^ became credibly positive at 4.4 ms for upright faces (0.267 [0.162, 0.377]) and inverted faces (0.104 [0.012, 0.207]), and increased by 6.2 ms to 0.458 [0.347, 0.567] and 0.219 [0.109, 0.331], respectively.Meta-*d*^*′*^ followed a similar trajectory, increasing from 0.214 [0.121, 0.344] for upright faces and 0.069 [0.008, 0.160] for inverted faces at 4.4 ms to 0.327 [0.207, 0.488] and 0.188 [0.101, 0.308] at 6.2 ms. Upright faces produced greater expression *d*^*′*^ and meta-*d*^*′*^ at 4.4, 5.3, and 6.2 ms, with all corresponding 95% credible intervals excluding zero, in line with our prior conventional non-hierarchical SDT analyses [8]. Neither orientation reached 75% expression accuracy within the sampled exposure range; the extrapolated ED_75_ estimates are reported in Supplementary Table S5.

Together, these findings reveal distinct behavioural trajectories for intact-face localisation and expression identification. Localisation approached high accuracy within the sampled range, whereas expression information continued to develop at the longest exposures. In line with our previous study, these results also establish 1.7, 4.4, and 6.2 ms as conditions providing weak, intermediate, and stronger perceptual evidence for the EEG experiment.

### Behavioural sensitivity and perceptual awareness increased with exposure duration in Experiment 2

In Experiment 2, Bayesian hierarchical type-1 SDT analysis showed that location sensitivity was indistinguishable from chance at 1.7 ms (median *d*^*′*^ = 0.107, 95% credible interval [CrI] = [-0.009, 0.224]) but increased sharply at 4.4 ms (*d*^*′*^ = 1.689 [1.569, 1.809]) and again at 6.2 ms (*d*^*′*^ = 2.617 [2.494, 2.742]; Fig. 3a). The increase was credible both between 1.7 and 4.4 ms (Δ*d*^*′*^ = 1.581 [1.543, 1.620]) and between 4.4 and 6.2 ms (Δ*d*^*′*^ = 0.928 [0.873, 0.983]). Our Bayesian hierarchical type-2 SDT analysis showed that PAS-based metacognitive sensitivity followed the same progression. Location meta-*d*^*′*^ was near zero at 1.7 ms (0.051 [-0.004, 0.124]) but was clearly positive at 4.4 ms (0.891 [0.611, 1.327]) and 6.2 ms (1.143 [0.789, 1.698]). Meta-*d*^*′*^ increased by 0.839 [0.580, 1.243] between 1.7 and 4.4 ms and by a further 0.251 [0.169, 0.383] between 4.4 and 6.2 ms. Metacognitive efficiency nevertheless remained below one: median M-ratios were 0.528 [0.365, 0.783] at 4.4 ms and 0.437 [0.302, 0.650] at 6.2 ms. Thus, once participants could reliably localise the intact face, their subjective reports tracked the accuracy of that perceptual decision, although not with fully efficient access to all decision-relevant information.

**Figure 3:**
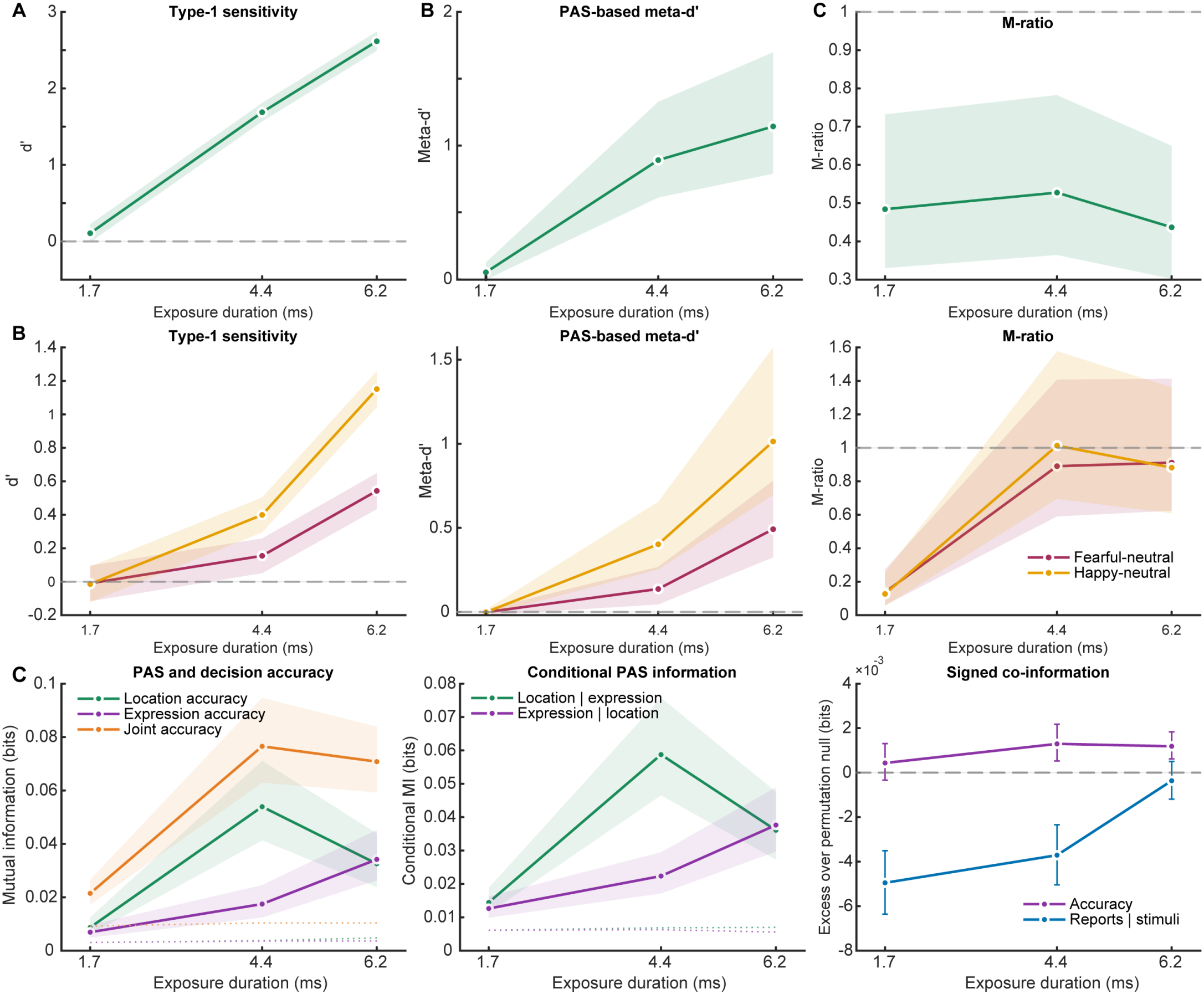
Behavioural sensitivity, metacognitive sensitivity, and information shared with perceptual awareness. **a**, Bayesian hierarchical estimates of location type-1 sensitivity (*d*^*′*^), PAS-based metacognitive sensitivity (meta-*d*^*′*^), and metacognitive efficiency (M-ratio) across exposure durations. **b**, Corresponding estimates for expression discrimination in the fearful–neutral and happy–neutral contexts. Points show posterior medians and error bars show 95% credible intervals. Grey dashed lines denote zero sensitivity or an M-ratio of one. **c**, Information-theoretic relationships among PAS, location accuracy, and expression accuracy. The left plot shows mutual information between PAS and location accuracy, expression accuracy, or their joint outcome. The middle plot shows the information shared by PAS and each accuracy measure after conditioning on the other accuracy measure. The right plot shows co-information among PAS, location accuracy, and expression accuracy, together with co-information among the three reports conditional on the presented stimuli. Positive co-information indicates a redundancy-dominant balance and negative co-information a synergy-dominant balance. Solid lines and shaded regions in the first two plots show group estimates and 95% Bayesian-bootstrap intervals; dotted lines show permutation-null means. Co-information is expressed relative to its permutation-null mean, with error bars showing the corresponding 95% null interval.

Expression sensitivity also increased with exposure duration, but depended on emotional context (Fig. 3b). At 1.7 ms, sensitivity was indistinguishable from zero for both fearful–neutral (*d*^*′*^ = -0.010 [-0.116, 0.096]) and happy– neutral decisions (*d*^*′*^ = -0.014 [-0.121, 0.090]). At 4.4 ms, sensitivity became credibly positive for fearful faces (*d*^*′*^ = 0.155 [0.051, 0.258]) and was larger for happy faces (*d*^*′*^ = 0.399 [0.295, 0.503]). At 6.2 ms, sensitivity increased further in both contexts (fearful: *d*^*′*^ = 0.543 [0.434, 0.649]; happy: *d*^*′*^ = 1.151 [1.043, 1.258]).

The happy–fearful difference was absent at 1.7 ms (Δ*d*^*′*^ = -0.005 [-0.072, 0.062]), emerged at 4.4 ms (Δ*d*^*′*^ = 0.245 [0.184, 0.306]), and increased at 6.2 ms (Δ*d*^*′*^ = 0.609 [0.537, 0.679]). PAS-based expression meta-*d*^*′*^ showed a closely corresponding pattern. It was near zero at 1.7 ms, became positive at 4.4 ms (fearful: 0.137 [0.045, 0.268]; happy: 0.402 [0.254, 0.655]), and increased further at 6.2 ms (fearful: 0.492 [0.325, 0.780]; happy: 1.014 [0.697, 1.573]). The happy–fearful meta-*d*^*′*^ difference was present at 4.4 ms (0.267 [0.175, 0.423]) and increased at 6.2 ms (0.520 [0.353, 0.816]).

Once expression sensitivity was measurable, expression M-ratios were compatible with one. At 4.4 ms, median M-ratios were 0.890 [0.590, 1.408] for fearful faces and 1.013 [0.695, 1.577] for happy faces; at 6.2 ms, they were 0.912 [0.624, 1.413] and 0.882 [0.609, 1.361], respectively. Together, these new analyses confirm and expand our previous findings [8] by indicating that reliable localisation and PAS-based awareness were established at 4.4 ms, whereas expression sensitivity continued to develop substantially between 4.4 and 6.2 ms.

### Perceptual awareness initially tracked location and only later expression accuracy

As a behavioural counterpart to the EEG information analyses, we quantified the information shared by PAS and the two objective discrimination decisions in Experiment 2.

PAS contained a small but reliable amount of information about location accuracy already at 1.7 ms (*p*_Holm_ = .0015), whereas PAS–expression information was not distinguishable from its finite-sample null at this duration (*p*_Holm_ = .223; Fig. 3c). At 4.4 ms, PAS shared information with both location accuracy (median MI = 0.0539 bits, 95% interval = [0.0413, 0.0711], *p*_Holm_ = .0015) and expression accuracy (0.0175 bits [0.0124, 0.0245], *p*_Holm_ = .0015). At 6.2 ms, PAS–location information remained reliable (0.0325 bits [0.0238, 0.0439], *p*_Holm_ = .0015), while PAS– expression information increased to a comparable magnitude (0.0342 bits [0.0263, 0.0452], *p*_Holm_ = .0015).

The two relationships followed different exposure-dependent trajectories. PAS–location information increased from 1.7 to 4.4 ms (ΔMI = 0.0450 bits [0.0322, 0.0626]) and from 1.7 to 6.2 ms (0.0238 bits [0.0141, 0.0357]), but was lower at 6.2 than at 4.4 ms (-0.0212 bits [-0.0383, -0.0060]). By contrast, PAS–expression information increased between every successive duration: 4.4–1.7 ms, ΔMI = 0.0106 bits [0.0046, 0.0181]; 6.2–1.7 ms, 0.0272 bits [0.0188, 0.0384]; and 6.2–4.4 ms, 0.0165 bits [0.0090, 0.0261]. Conditional MI analyses produced the same dissociation: location contributed information about PAS beyond expression accuracy from the shortest duration onward, whereas the independent contribution of expression accuracy became reliable only at 4.4 and 6.2 ms.

Co-information further indicated a small redundancy-dominant component shared among PAS, location accuracy, and expression accuracy at 4.4 ms (observed-minus-null co-information = 0.00129 bits, *p*_Holm_ = .0090) and 6.2 ms (0.00119 bits, *p*_Holm_ = .0015), but not at 1.7 ms. Thus, awareness was not equally related to all perceptual attributes from the outset. Its earliest reliable relationship was with localisation, whereas expression-related information entered subjective reports progressively as exposure increased. Experiment 2 behavioural and report-information estimates are provided in Supplementary Table S6.

### Exposure duration broadly altered visually evoked activity, whereas emotion-related ERPs emerged only at 6.2 ms

Our new cluster-based ERP analysis showed that exposure duration produced widespread changes in the evoked response (Fig. 4a,b). The omnibus cluster test identified a reliable effect extending from approximately 48 to 596 ms across the scalp (cluster *p* < .0001). All three pair-wise duration contrasts were significant after correction (all *p*_Holm_ = .0006). Differences between 1.7 and 4.4 ms began at approximately 48–52 ms and continued through the remainder of the epoch. The 4.4–6.2-ms and 1.7– 6.2-ms contrasts likewise revealed broad, sustained differences involving both positive- and negative-polarity electrode clusters. Ultra-brief changes in physical exposure therefore produced extensive differences in the ensuing visual response rather than merely rescaling a restricted late component.

**Figure 4:**
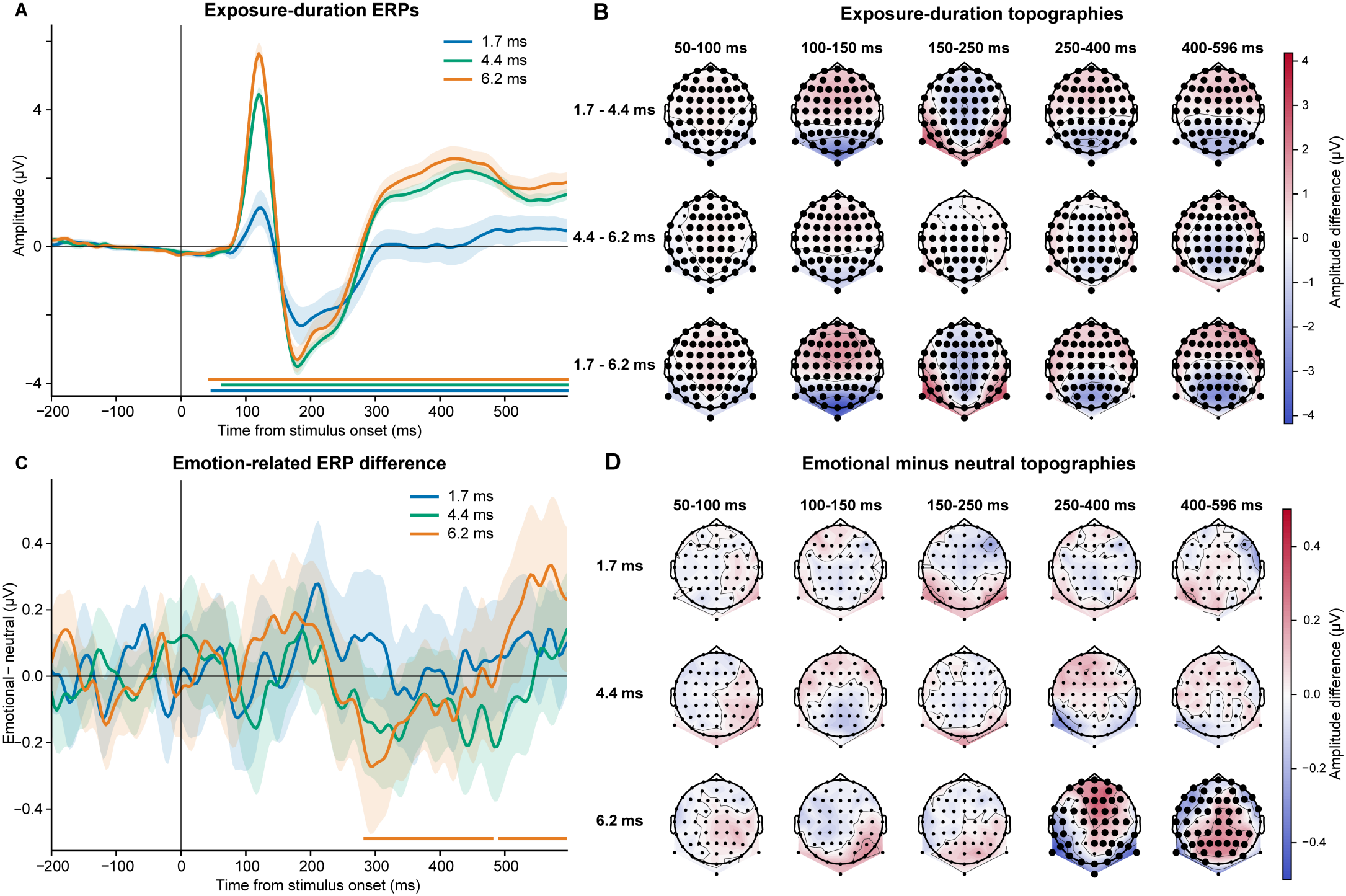
Exposure-duration and emotion-related event-related potentials. **a**, Baseline-corrected ERPs averaged across O1, Oz, O2, PO7, PO3, POz, PO4, and PO8 for the three exposure durations. Lines show participant means and shading shows 95% confidence intervals. Horizontal bars below the waveforms mark significant sensor-by-time clusters for the three planned pairwise duration contrasts indicated beside the plot. **b**, Scalp distributions of the pairwise exposure-duration differences averaged over the indicated time windows. **c**, Emotional-minus-neutral ERP difference at each exposure duration, averaged across the same posterior electrodes. Coloured horizontal bars indicate significant duration-specific emotion–neutral clusters. **d**, Scalp distributions of the emotional-minus-neutral difference across the indicated time windows and exposure durations. In b and d, red denotes a positive voltage difference and blue a negative voltage difference; large bold electrode markers identify electrodes belonging to a significant cluster during the corresponding interval, whereas small markers show the remaining electrodes. Statistical inference used two-sided sensor-by-time cluster-based permutation tests, with Holm correction across the relevant planned contrasts.

Emotion-related activity showed a more selective pattern (Fig. 4c,d). Emotional and neutral faces did not differ reliably at 1.7 ms (minimum *p*_Holm_ = .462) or 4.4 ms (*p*_Holm_ = .462). At 6.2 ms, however, emotional and neutral faces produced distributed differences from approximately 284 to 596 ms (map-level *p*_Holm_ = .003). These included clusters of both scalp polarities, consistent with a spatially distributed difference rather than a uniform voltage shift. The 6.2-ms condition was therefore the first exposure duration at which a reliable emotion–neutral ERP difference emerged, which is in line with our previous analysis [8].

### Expression information generalised across face identities only at the longest exposure

Time-resolved multivariate decoding provided a complementary test of the information expressed across the scalp (Fig. 5a). The primary classifiers were evaluated on face identities that had not appeared in their training folds. Location could be decoded throughout the post-stimulus interval at both 4.4 and 6.2 ms (0–596 ms, both Holm-adjusted *p* = .0008), but not at 1.7 ms. This widespread location signal paralleled the marked increase in behavioural localisation sensitivity.

**Figure 5:**
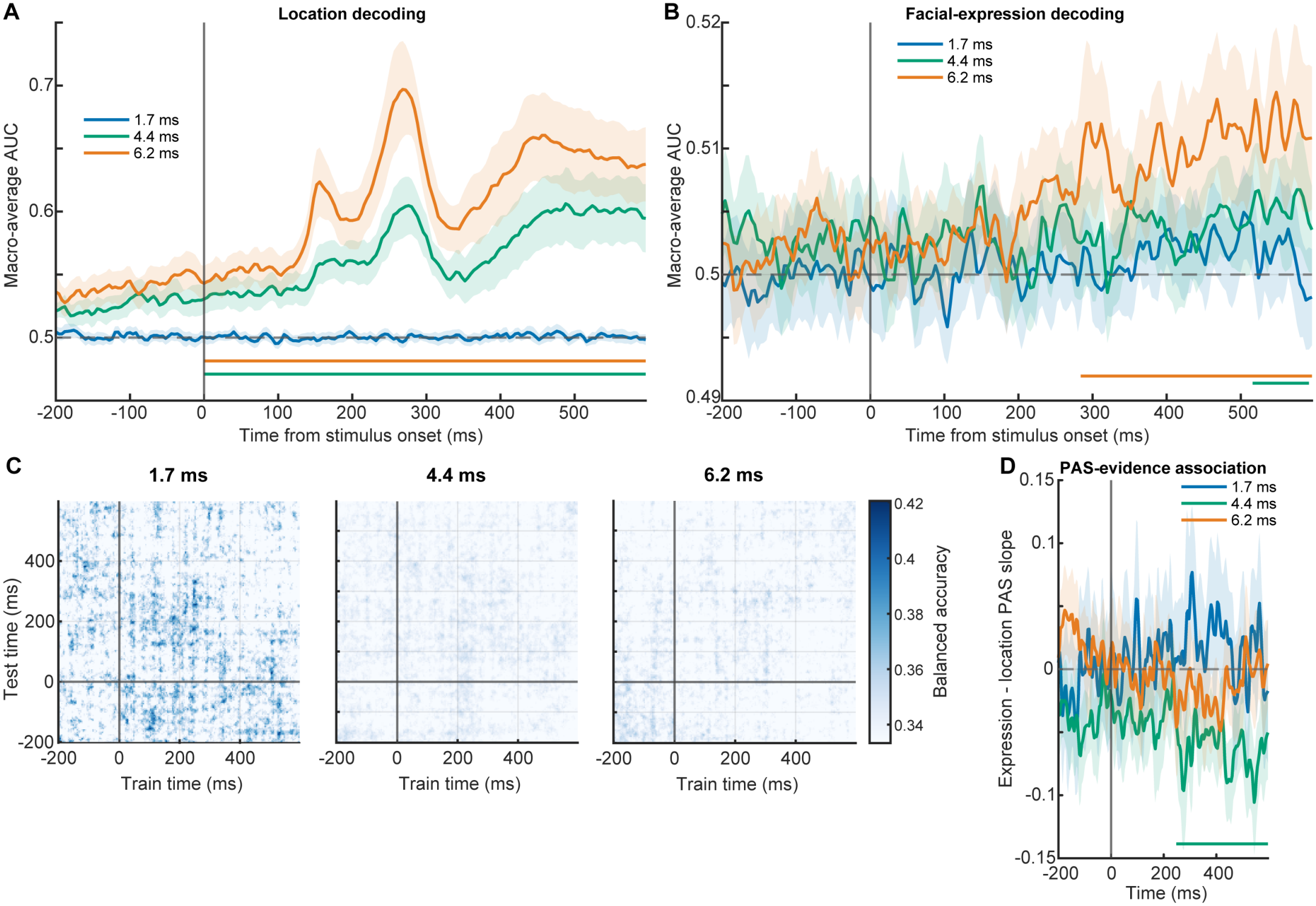
Time-resolved decoding and its relationship with perceptual awareness. **a**, Identity-generalising decoding of intact-face location. **b**, Identity-generalising decoding of facial expression. Lines show mean macro-averaged area under the receiver-operating-characteristic curve across participants, shading shows 95% confidence intervals, and the grey dashed line denotes chance performance. Coloured horizontal bars mark significant temporal clusters after correction across the three exposure durations. **c**, Facial-expression temporal-generalisation matrices for each exposure duration. Classifiers were trained at the times shown on the horizontal axis and tested at the times shown on the vertical axis. Values are balanced accuracies averaged across the matched low-PAS and high-PAS analyses; chance performance was one third. **d**, Difference between the trial-level PAS slopes for expression and location evidence. Negative values indicate a stronger association of PAS with location evidence than with expression evidence. Lines show participant means, shading shows 95% confidence intervals, and coloured horizontal bars indicate significant temporal clusters. Vertical lines mark stimulus onset.

Expression decoding followed a later and more restricted trajectory. No identity-generalising expression cluster survived correction at 1.7 or 4.4 ms. At 6.2 ms, expression decoding was reliably above chance from 348 to 596 ms (p_Holm_ = .0044; Fig. 5b,c). A direct comparison confirmed that identity-generalising expression decoding was stronger at 6.2 than at 4.4 ms, with significant differences from 288 to 324 ms (cluster mass = 26.72, p = .024) and from 440 to 472 ms (cluster mass = 22.91, p = .040). Thus, the emergence of expression decoding that generalised across faces reflected a reliable exposure-dependent increase rather than merely significance at one duration and non-significance at another.

A secondary analysis using trial-wise, stratified folds more closely reproduced the original analysis [8]. Under this less restrictive validation scheme, expression decoding was detected at both 4.4 ms (516–592 ms, *p*_Holm_ = .032) and 6.2 ms (284–596 ms, *p*_Holm_ = .0042; Supplementary Fig. S4). The difference from the original result therefore arose primarily from requiring generalisation to unseen identities: the late 4.4-ms signal was detectable across held-out trials but did not generalise reliably across identities, whereas the 6.2-ms representation survived both tests.

### Subjective clarity was associated more strongly with location than expression evidence

PAS was not provided to the classifiers. We instead asked whether the amount of held-out decision evidence on each trial varied with the participant’s subsequent awareness report. Expression evidence did not vary reliably with PAS at any exposure duration after correction. Location evidence, by contrast, increased with PAS across two extended intervals at 4.4 ms (0–208 ms, *p*_Holm_ = .0048; 240– 596 ms, *p*_Holm_ = .0016) and from 312 to 452 ms at 6.2 ms (*p*_Holm_ = .0042). No relationship was detected at 1.7 ms.

At 4.4 ms, the association with PAS was significantly stronger for location than expression evidence from 248 to 596 ms (*p*_Holm_ = .0018; Fig. 5d). None of the direct duration contrasts for expression evidence was significant. These results argue against a general increase in expression-specific evidence with subjective clarity. Instead, PAS primarily tracked neural information supporting successful localisation or more general stimulus visibility before tracking emotional expression.

### Perceptual awareness was represented locally without robust EPN–LPP information transfer

We next examined information dynamics between the posterior EPN and centroparietal LPP sensor nodes (Fig. 6a,b). Neither node carried reliable information about emotional versus neutral status after correction. Posterior emotion information approached significance at 6.2 ms but did not survive correction across the planned comparisons (*p*_Holm_ = .065).

**Figure 6:**
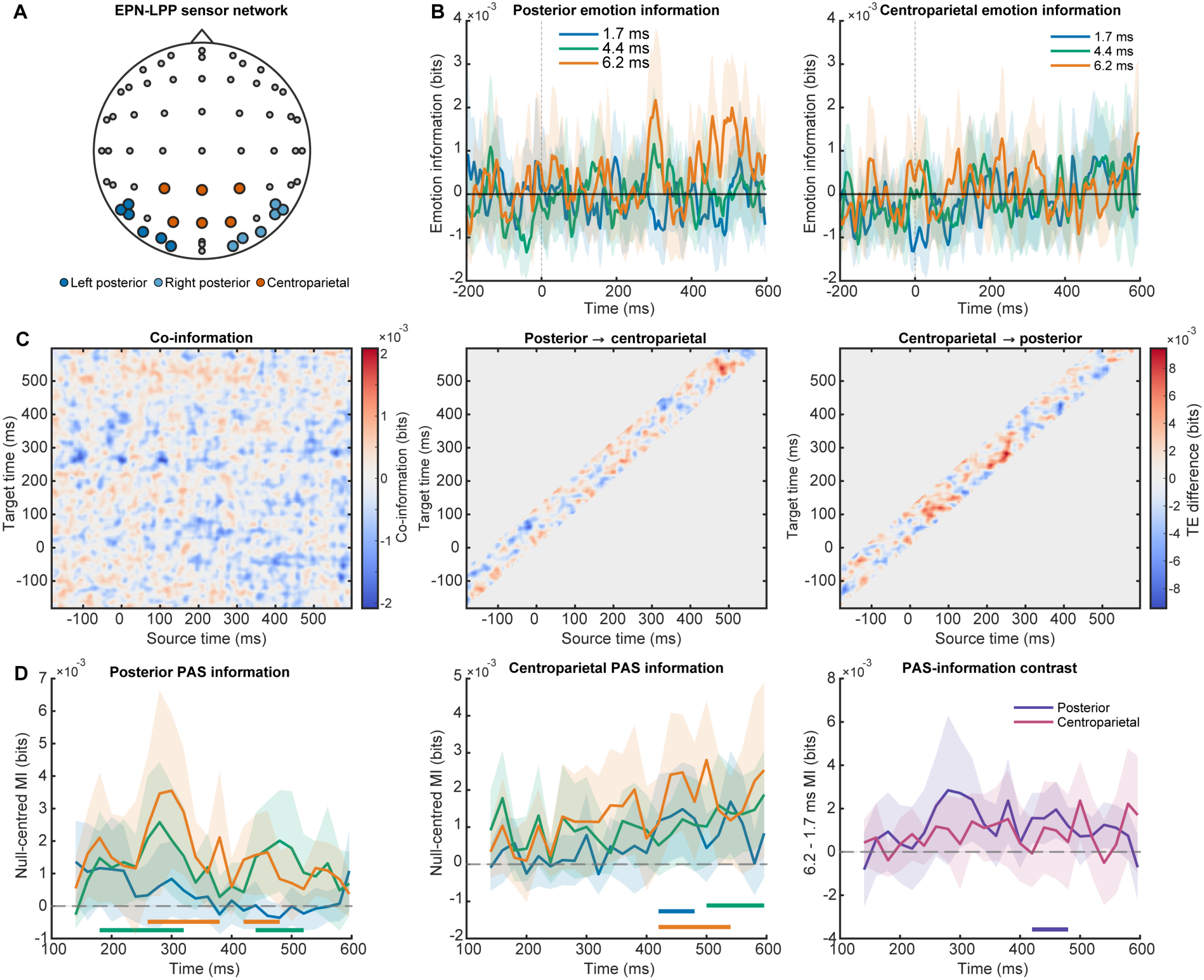
Information represented within and shared between posterior EPN and centroparietal LPP sensor nodes. **a**, Prespecified left-posterior, right-posterior, and centroparietal sensor groupings. The two posterior groupings were combined to define the bilateral EPN node. **b**, Gaussian-copula mutual information about emotional versus neutral status at the posterior and centroparietal nodes for each exposure duration. Lines show participant means and shading shows 95% confidence intervals. **c**, Co-information and emotional-minus-neutral transfer-entropy differences at 6.2 ms. The co-information map represents the net redundancy–synergy balance across posterior and centroparietal time points, with positive values indicating a redundancy-dominant balance and negative values a synergy-dominant balance. Transfer-entropy maps show directed predictive dependence in the posterior-to-centroparietal and centroparietal-to-posterior directions; positive values indicate greater transfer entropy for emotional than neutral faces. Grey regions contain source–target combinations outside the admissible temporal direction or lag range. Black contours, when present, identify significant clusters. **d**, Null-centred mutual information between PAS and posterior or centroparietal activity after adjustment for expression, identity, target side, block, and location accuracy. The rightmost plot shows the direct 6.2-minus-1.7-ms contrast. Coloured horizontal bars mark significant temporal clusters. Information quantities are expressed in bits.

Emotional status also did not reliably alter directed predictive dependence between the two nodes. No posterior-to-centroparietal, centroparietal-to-posterior, directional-interaction, or duration effect survived correction; the smallest Holm-adjusted probability was .554 (Fig. 6c). Thus, although emotional and neutral faces produced different distributed ERPs at 6.2 ms, this difference was not accompanied by detectable directed dependence between the predefined EPN and LPP nodes.

The initial co-information analysis yielded positive, redundancy-dominant clusters at 1.7 ms (*p*_Holm_ = .014) and 4.4 ms (*p*_Holm_ *p* = .049 and .0024). None survived the more stringent paired-label validation analysis, however, and the direct exposure-duration contrasts were also nonsignificant (Fig. 6c). We therefore found no robust evidence that exposure duration altered redundant emotion information shared by the EPN and LPP nodes.

PAS information showed a clearer local pattern (Fig. 6d). After adjustment for expression, identity, target side, block, and location accuracy, posterior activity carried information about PAS at 4.4 ms from 180 to 320 ms (*p*_Holm_ = .0036) and from 440 to 520 ms (*p*_Holm_ = .040). At 6.2 ms, posterior PAS information was present from 260 to 380 ms and from 420 to 480 ms (both *p*_Holm_ = .0080). Centroparietal activity carried PAS information at 1.7 ms from 420 to 480 ms (*p*_Holm_ = .030), at 4.4 ms from 500 to 596 ms (*p*_Holm_ = .0080), and at 6.2 ms from 420 to 540 ms (*p*_Holm_ = .0080). Late posterior PAS information was greater at 6.2 than at 1.7 ms from 420 to 480 ms (*p*_Holm_ = .013); the remaining duration contrasts were not significant. Subjective clarity was therefore represented locally in both posterior and centroparietal activity despite the absence of transfer-entropy effects or reliable exposure-duration differences in EPN-LPP co-information.

### Emotion information emerged in the EPN and centroparietal–occipital predictive dependence changed with exposure

The P1-network analyses examined whether early occipital activity contributed to subsequent emotion-sensitive processing (Fig. 7a,b). The occipital P1 node did not carry reliable information about emotional versus neutral status at any exposure duration. When the posterior EPN node was included, however, it carried reliable emotion information at 6.2 ms from 296 to 320 ms (*p*_Holm_ = .0036). EPN emotion information was also greater at 6.2 than at 1.7 ms from 296 to 312 ms (*p*_Holm_ = .030). No corresponding effect was detected at the LPP node. Emotion information therefore became detectable during the EPN interval at the longest exposure, rather than during the preceding P1 interval.

**Figure 7:**
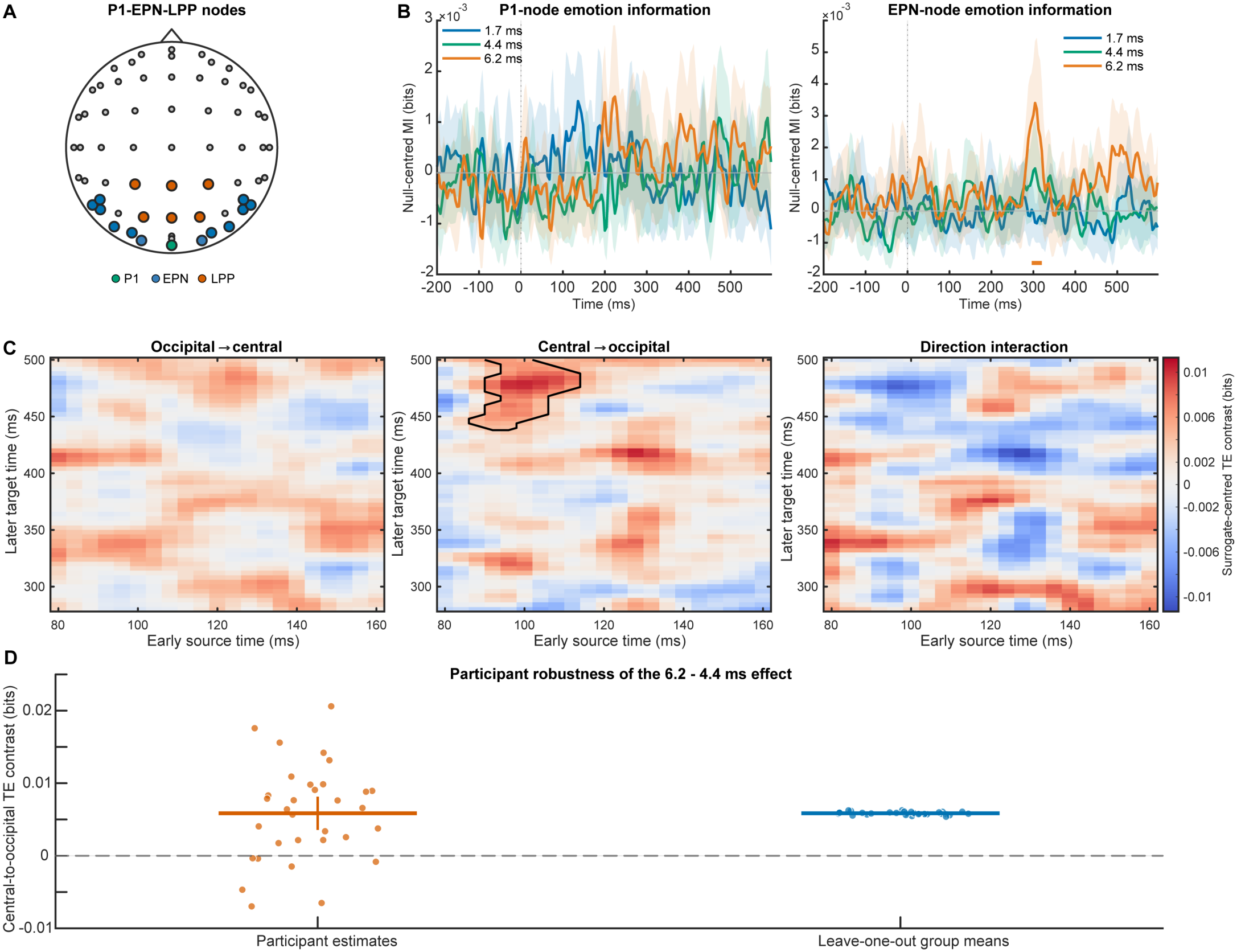
Local emotion information and predictive dependence across P1, EPN, and LPP sensor nodes. **a**, Prespecified occipital P1, posterior EPN, and centroparietal LPP sensor groupings. O1 and O2 were excluded from the EPN node to prevent overlap with the P1 node. **b**, Null-centred mutual information about emotional versus neutral status at the P1 and EPN nodes for each exposure duration. Lines show participant means, shading shows 95% confidence intervals, and coloured horizontal bars mark significant temporal clusters. **c**, Surrogate-centred 6.2-minus-4.4-ms contrasts in the emotional–neutral transfer-entropy effect across the broader P1–LPP analysis windows. Maps show occipital-to-central predictive dependence, central-to-occipital predictive dependence, and their directional interaction. The horizontal axis represents early source time and the vertical axis later target time. Positive values indicate that the emotional–neutral transfer-entropy contrast was more positive at 6.2 than at 4.4 ms. The black contour identifies the significant central-to-occipital cluster. **d**, Participant-level mean transfer-entropy contrasts within that cluster and the corresponding leave-one-participant-out group means. Dots show individual estimates; thick horizontal lines and vertical bars show the group mean and its 95% confidence interval. Information quantities are expressed in bits.

The P1–EPN analyses provided no reliable evidence of co-information or directed predictive dependence between the nodes (Supplementary Fig. S5a). On the other hand, co-information revealed in the P1–LPP model an exposure-dependent change in the emotional-neutral transfer entropy contrast (Fig. 7c; Supplementary Fig. S5b). The emotion effect in predictive dependence from early centroparietal activity to later occipital activity was greater at 6.2 than at 4.4 ms, linking centroparietal source times from 88 to 136 ms with occipital target times from 436 to 500 ms (p_Holm_ = .0090).

Importantly, this effect remained significant after transfer entropy was centred on condition-preserving sourcetrial surrogate estimates. The resulting cluster linked centroparietal activity from 88 to 112 ms with occipital activity from 440 to 500 ms (*p*_Holm_ = .0108). The reverse occipital-to-centroparietal contrast was not significant, but neither was the formal interaction between directions. The effect was therefore detected in the centroparietal-to-occipital analysis, but the data did not establish that this direction was stronger than the reverse direction.

Several checks supported the robustness of this result (Supplementary Table S7). Extending the late analysis interval to 560 ms yielded a nearly identical cluster spanning 84–112 ms by 440–504 ms (*p* = .0026), showing that the effect was not produced by the original 500-ms boundary. The surrogate-centred effect was positive in 25 of 32 participants (mean = 0.00585 bits, SEM = 0.00117, d_z_ = 0.89), and the group mean remained stable when each participant was omitted in turn (range = 0.00538–0.00627 bits; Fig. 7d). The effect was not independently reproduced using Oz and Pz alone or using voltage rather than CSD (all *p*_Holm_ *≥*.402), suggesting that it depended on the distributed CSD representation. The fearful–neutral contrast approached but did not survive correction (*p*_Holm_ = .066), and neither the happy–neutral contrast nor the fearful-versus-happy interaction was significant. The pooled emotion effect therefore could not be attributed specifically to fearful expressions.

Together, these findings distinguish the local representation of emotion from predictive dependence between electrode networks. Emotion information became locally detectable during the EPN interval at 6.2 ms, while the broader P1–LPP analysis revealed an exposure-dependent relationship between early centroparietal and later occipital activity. This pattern is consistent with recurrent processing under the longest exposures.

### Trial-level neural evidence linked perceptual decisions and awareness

Finally, we tested whether neural evidence extracted from theoretically defined EPN and LPP electrode patterns predicted trial-by-trial behaviour. Location could be decoded from both patterns at 4.4 and 6.2 ms (Fig. 8a). For the posterior EPN pattern, mean balanced accuracy was 0.612 at 4.4 ms (*p*_Holm_ = 6.4 *×*10^*−*11^) and 0.671 at 6.2 ms (*p*_Holm_ = 7.5 *×*10^*−*12^). For the centroparietal LPP pattern, accuracy was 0.558 at 4.4 ms (*p*_Holm_ = 1.8*×* 10^*−*7^) and 0.587 at 6.2 ms (*p*_Holm_ = 8.5 *×*10^*−*10^). Neither pattern decoded location at 1.7 ms. Expression decoding was considerably weaker: only the 4.4-ms LPP classifier survived correction (balanced accuracy = 0.346, chance = 0.333, *p*_Holm_ = .020; Fig. 8b).

**Figure 8:**
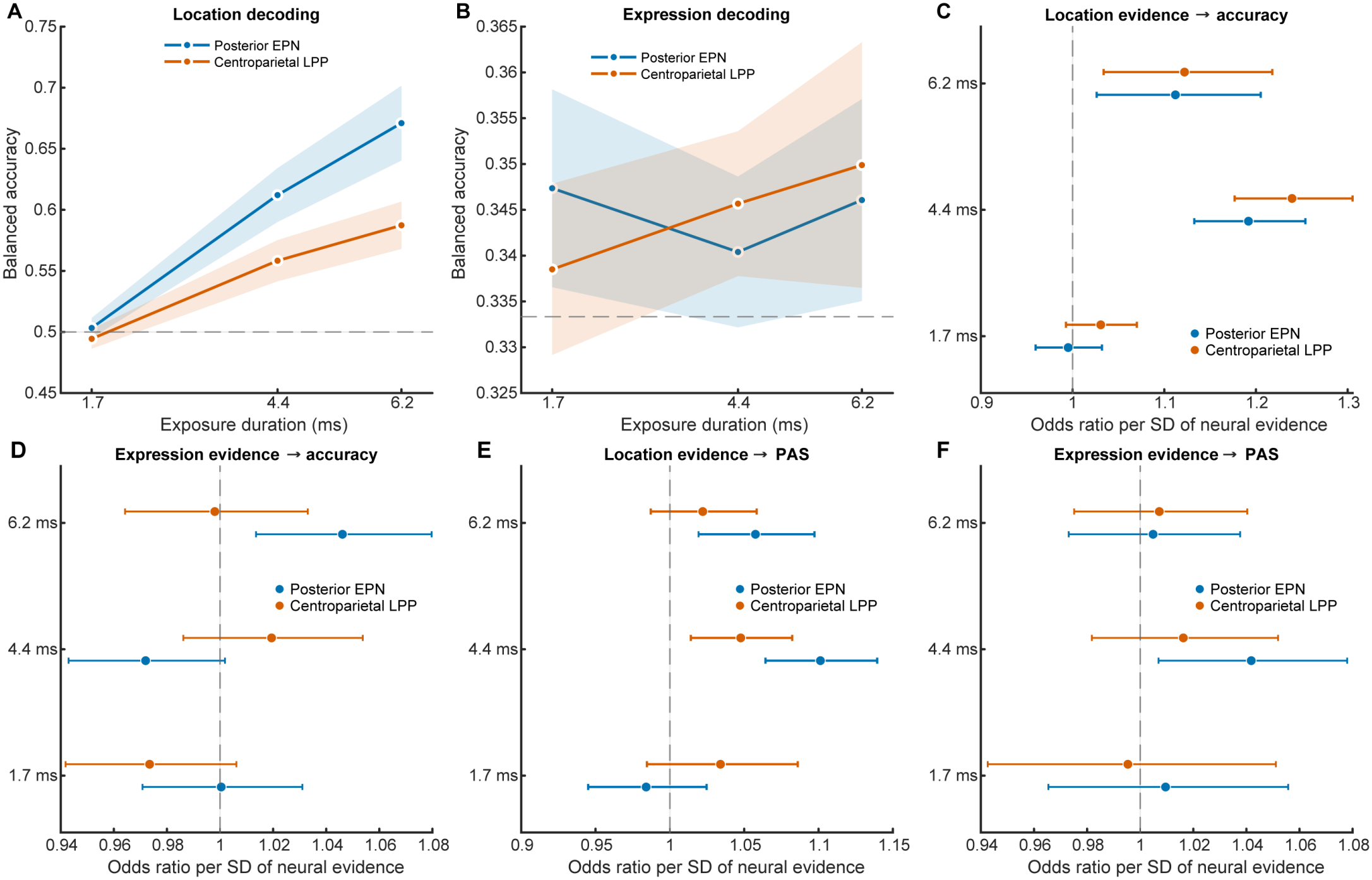
Neural evidence predicting objective decisions and perceptual awareness. **a**, Cross-validated location decoding from posterior EPN and centroparietal LPP electrode patterns. **b**, Corresponding three-class facial-expression decoding. Points show mean balanced accuracy and error bars show 95% confidence intervals; grey dashed lines indicate chance performance. **c**, Associations between location evidence and location accuracy. **d**, Associations between expression evidence and expression accuracy. **e**, Associations between location evidence and PAS. **f**, Associations between expression evidence and PAS. Panels c–f show odds ratios associated with a one-standard-deviation increase in cross-validated neural evidence, with horizontal error bars denoting 95% confidence intervals. The grey dashed line marks an odds ratio of one. Accuracy outcomes were analysed using logistic mixed-effects models, and PAS using participant-wise cumulative-logit models followed by group-level inference. Blue denotes the posterior EPN pattern and orange the centroparietal LPP pattern.

Location evidence robustly predicted location accuracy at both suprathreshold durations (Fig. 8c). Posterior evidence predicted location accuracy at 4.4 ms (OR = 1.192 [1.133, 1.254], *p*_Holm_ = 6.5 *×* 10^*−*11^) and 6.2 ms (OR = 1.112 [1.026, 1.205], *p*_Holm_ = .029). Centroparietal evidence showed the same pattern at 4.4 ms (OR = 1.239 [1.177, 1.305], *p*_Holm_ = 3.2*×* 10^*−*15^) and 6.2 ms (OR = 1.122 [1.034, 1.218], *p*_Holm_ = .023). No location-evidence effect was detected at 1.7 ms.

Despite this modest group-average expression decoding, trial-level posterior expression evidence predicted expression accuracy at 6.2 ms (Fig. 8d). Each standard-deviation increase in evidence was associated with a 4.6% increase in the odds of a correct expression response (odds ratio [OR] = 1.046 [1.014, 1.080], *p*_Holm_ = .031), and this relationship was stronger than at 4.4 ms (*p*_*Holm*_ = .0060). No corresponding relationship was detected for LPP expression evidence.

Crucially, PAS was predicted by location evidence (Fig. 8e) but not by expression evidence (Fig. 8f). The joint PAS tests were significant for posterior location evidence (*p*_Holm_ = 2.8*×* 10^*−*5^) and centroparietal location evidence (*p*_Holm_ = .020), but not for expression evidence from either electrode pattern. Posterior location evidence predicted higher PAS ratings at 4.4 ms (OR = 1.101 [1.064, 1.139], *p*_Holm_ = 1.4 *×*10^*−*5^) and 6.2 ms (OR = 1.058 [1.019, 1.097], *p*_Holm_ = .021). Centroparietal location evidence also predicted PAS at 4.4 ms (OR = 1.048 [1.014, 1.082], *p*_Holm_ = .026). The relationship between posterior location evidence and PAS was stronger at 4.4 than at 1.7 ms (*p*_Holm_ = .00078) and at 6.2 than at 1.7 ms (*p*_Holm_ = .035). Complete EEG cluster and map results are reported in Supplementary Table S8.

## DISCUSSION

The present study shows that increasing visual exposure changed the organisation of facial-expression information rather than simply strengthening a fixed representation. Information supporting intact-face localisation and subjective clarity emerged before expression-related neural patterns generalised to faces excluded from classifier training. With additional exposure, emotional and neutral faces became neurally distinguishable, emotion information became detectable during the EPN interval, and predictive dependence between sensor patterns changed. Local differentiation, representational stability, subjective clarity, and distributed neural dependence therefore followed partly distinct trajectories.

The original study established that localisation and awareness preceded reliable expression processing, whereas stronger expression decoding and emotion-related EPN and LPP effects appeared at 6.2 ms [8]. The present analyses address questions that this sequence left unresolved. Hierarchical Bayesian modelling reproduced the behavioural ordering while characterising the graded development of objective and metacognitive sensitivity. Multivariate analyses reproduced the original decoding pattern and showed that expression representations generalised across faces only with longer exposure. Trial-level analyses related neural evidence to accuracy and perceptual clarity. Information-theoretic analyses distinguished local emotion information from redundancy- or synergy-dominant relationships and directed predictive dependence across P1, EPN, and LPP stages. Together, these analyses characterise how facial-expression information becomes stable, behaviourally relevant, distributed across neural responses, and related to awareness.

Across both experiments, facial attributes became available at different rates. The densely sampled psychometric functions from Experiment 1 showed that intact-face localisation approached high accuracy within a few milliseconds, whereas expression identification was still developing at the longest exposure. Estimated expression thresholds beyond the observed range should not be interpreted as absolute perceptual limits, but their separation from localisation thresholds indicates that locating a coherent face required less visual evidence than identifying its expression. Inversion delayed objective and metacognitive sensitivity, consistent with an early contribution of configural face processing [39–42]. Fearful expressions did not facilitate localisation, arguing against emotional salience accelerating the earliest intact–scrambled discrimination [43, 44]. Experiment 2 confirmed this ordering at three theoretically informative durations and showed that happy and fearful expressions did not become discriminable at identical rates. These findings support distinct exposure-dependent trajectories for localisation and expression processing rather than a common perceptual threshold for all facial attributes.

The relationship between objective performance and PAS also changed with available evidence. PAS initially reflected localisation more strongly, but its relationship with expression accuracy increased with exposure. Conditional mutual information showed distinguishable contributions of location and expression accuracy, whereas the small redundancy-dominant co-information identified only limited information shared among all three reports [24, 45, 46]. Subjective clarity therefore appeared to be anchored initially in the location or general visibility of the intact face and to incorporate expression-related information as that information became more reliable.

The EEG findings revealed a corresponding change in stimulus representation. Small increases in exposure altered the evoked response widely, whereas reliable emotional–neutral differentiation appeared only at the longest exposure, during the late EPN and LPP period. The visual system was therefore sensitive to additional input before evoked activity reliably distinguished emotional content [11, 47]. This places the previously reported ERP effect within the behavioural progression across the experiments [8], although cluster boundaries should not be interpreted as precise neural onset or offset times [48]. Multivariate decoding further showed that expression information detected at 4.4 ms did not generalise reliably to faces excluded from training, whereas decoding at 6.2 ms survived both validation procedures and was significantly stronger than at 4.4 ms in the direct duration comparison. Additional exposure thus produced an expression representation that generalised more reliably across the sampled face identities. Location information generalised earlier, consistent with its faster behavioural emergence. Successful decoding alone therefore does not establish the stability or generality of the underlying representation [49, 50].

Trial-level analyses distinguished information supporting objective decisions from information associated with subjective clarity. Once localisation was reliable, stronger EEG evidence for the intact face’s location was associated with more accurate location judgements and higher PAS ratings. Posterior expression evidence predicted expression accuracy at the longest exposure but was not reliably associated with PAS. This does not imply that subjective clarity was unrelated to expression perception, because PAS shared increasing behavioural information with expression accuracy and expression metacognitive sensitivity became reliable as discrimination improved. Instead, the neural association with PAS was not specific to the decoded expression representation. Subjective clarity may integrate several sources of perceptual and decision-related evidence, with the most reliable dimension exerting the greatest influence. Contributions from decision formation or response confidence cannot be excluded, however, because location and expression were reported through a compound decision before PAS [18, 19, 51, 52].

The information-theoretic analyses separated local emotion sensitivity from dependence between processing stages. Emotion information was detectable during the EPN interval at the longest exposure but not during the preceding P1 interval. The EPN–LPP model showed redundancy-dominant co-information at 1.7 and 4.4 ms, but neither co-information nor transfer entropy changed reliably with exposure. Emotion-related information could therefore become locally detectable over posterior sensors without a corresponding change in the measured dependence between posterior and centroparietal patterns. Importantly, the P1–LPP analysis identified an exposure-dependent change in the emotional–neutral transfer-entropy contrast relating early centroparietal to later occipital activity. Its timing was compatible with recurrent processing at the longest exposure [16, 53, 54], although transfer entropy measures directed predictive dependence and cannot be interpreted as definitive evidence for causal communication [55]. Moreover, the formal directional comparison was nonsignificant, and the effect was not reproduced with single electrodes or voltage rather than distributed current-source-density patterns. It should therefore be interpreted as a change within the selected sensor representation, not necessarily as anatomical feedback between cortical regions.

These findings bear on theories associating awareness with recurrent or globally distributed processing [16, 17]. Behavioural sensitivity, expression generalisation across different faces, local emotion information, subjective clarity, and predictive dependence did not change in parallel. This separation suggests that they capture related but distinguishable aspects of perceptual processing. Sensor-level analyses cannot identify the cortical generators of these effects, establish anatomical communication, or measure global neuronal broadcasting without source reconstruction, and therefore may not distinguish recurrent-processing from global-workspace accounts.

Several considerations further delimit the conclusions. The three EEG durations represented behaviourally distinct stages identified independently in Experiment 1, with 4.4 ms being the shortest duration at which an inversion effect emerged. The P1, EPN, and LPP sensor groups were likewise defined from the previously reported ERP findings [8], reducing circularity but not allowing sensor-level effects to be equated with particular cortical regions. Fearful and happy faces were pooled because the main contrast concerned emotional intensity or salience relative to neutral expressions, consistent with the relevant ERP components [11, 47]. The findings therefore concern emotional–neutral differentiation rather than emotional valence. The compound response may have introduced dependencies between location and expression decisions, limiting their complete separation at the decision stage. Expression decoding was tested across only the limited set of face identities included in the stimulus set. Finally, the transfer-entropy result depended on distributed current-source-density patterns and was not reproduced with alternative signal representations.

Future work should test whether objective performance, subjective experience, and neural information show similar relationships in other perceptual domains. Relevant examples include the emergence of auditory information in awareness [56–62], the representation and metacognitive evaluation of internal physiological signals [63–70], and the integration of visual, tactile, and proprioceptive information in body ownership [71–80]. A common information-theoretic framework could reveal whether local, shared, and directed information follow comparable trajectories as evidence increases [24, 81].

Building on the processing sequence identified in the original study, the present analyses show that increasing exposure did not produce a single threshold or uniformly amplify a fixed representation. Evidence supporting localisation and subjective clarity preceded expression representations that generalised across faces. With additional exposure, emotional and neutral faces became neurally distinguishable, emotion information became locally detectable during the EPN interval, and predictive relationships between successive sensor patterns followed a partly distinct trajectory. Facial-expression processing under minimal vision is therefore better characterised as a progressive change in the content, stability, behavioural relevance, and neural distribution of perceptual information.

## AUTHOR CONTRIBUTIONS

R.C.L. conceived and conceptualised the study. R.C.L., D.C., and A.C. acquired funding. R.C.L., H.R., and D.C. designed the experiments. R.C.L programmed the experiments, collected the data, created the code, analysed the data, and created the figures. R.C.L. wrote the manuscript with input from A.G. and A.C.-J. R.C.L. edited the manuscript with input from all authors.

## DATA AVAILABILITY

The aggregated behavioural data from Experiments 1 and 2, and pre-processed EEG data from Experiment 2, trial-level metadata, participant-inclusion records, and the derived data underlying the figures and statistical analyses will be made publicly available through the Open Science Framework.

## CODE AVAILABILITY

All custom MATLAB, R, and Python code used for behavioural modelling, EEG preprocessing, ERP analyses, multivariate decoding, information-theoretic analyses, statistical inference, and figure generation will be made publicly available through the Open Science Framework. The repository will include instructions for reproducing the analyses and figures, together with software and package version information.

