## Supplementary Information for "The emergence and neural organisation of facial-expression information during ultra-brief vision"

### SUPPLEMENTARY METHODS

#### Bayesian hierarchical signal-detection models

Location and expression decisions were analysed separately with hierarchical meta-d' models implemented in R using hmetad, brms, cmdstanr, and Stan. PAS ratings 1 to 4 formed the ordered type-2 response. The Experiment 1 location model contained the 28 cells formed by seven exposure durations, two face orientations, and two expressions. Its expression model contained the 14 cells formed by exposure duration and orientation. The Experiment 2 location model contained three exposure-duration conditions, whereas its expression model contained the six cells formed by duration and emotional context (fearful versus neutral or happy versus neutral).

Each model estimated condition-specific log metacognitive efficiency, log(M), type-1 sensitivity (d'), type-1 criterion, and PAS thresholds. Participant-varying intercepts were included for log(M), d', and criterion, whereas threshold parameters were condition-specific fixed effects. M-ratio was calculated as exp[log(M)] and meta-d' as d' multiplied by M-ratio. Location d' and meta-d' were divided by $\sqrt{2}$ to express sensitivity under the two-alternative forced-choice convention used in the original study. This rescaling does not affect M-ratio. Because PAS indexed perceptual clarity rather than conventional confidence, meta-d' and M-ratio were interpreted as PAS-based metacognitive sensitivity and efficiency.

Weakly regularising priors were Normal(0, 0.7) for log(M), Normal(0, 1.5) for d', Normal(0, 1) for criterion, Normal(−0.5, 0.75) for the threshold-difference parameters, and Exponential(2) for participant-level standard deviations (**Supplementary Table S2**). Four chains were run with 2,000 warm-up iterations followed by 4,000 sampling iterations per chain. The target acceptance probability was .99 and the maximum tree depth was 15. Estimates and planned contrasts are reported as posterior medians with 95% credible intervals. Fits were accepted when the maximum $\hat{R}$ was no greater than 1.01, minimum bulk effective sample size was at least 400, and no divergent transitions or maximum-tree-depth exceedances occurred.

#### Hierarchical psychometric models for Experiment 1

Accuracy in Experiment 1 was additionally analysed with hierarchical cumulative-Gaussian psychometric functions. Location accuracy was modelled separately for upright-neutral, upright-fearful, inverted-neutral, and inverted-fearful faces. Expression accuracy was modelled separately for upright and inverted faces. The lower asymptote was fixed at .50. The upper asymptote incorporated a participant-varying lapse rate constrained to the interval from 0 to .15. Condition-specific group intercepts and positive slopes were combined with partially pooled participant deviations. Trials sharing a participant, condition, and duration were entered as binomial counts, which yields the same likelihood as the corresponding trial-wise Bernoulli observations.

The primary threshold was $ED_{75}$, the exposure duration at which expected accuracy reached .75 after allowing for lapse. Planned contrasts compared upright and inverted faces and, for localisation, fearful and neutral expressions. Thresholds outside the sampled interval of 0.8 to 6.2 ms were retained as model-based extrapolations rather than interpreted as directly observed perceptual

limits. Priors are reported in **Supplementary Table S4**. The four-chain sampling schedule and diagnostic criteria matched those of the signal-detection models.

#### PAS-related event-related potentials

PAS cells were markedly imbalanced, especially at the shortest exposure, so the primary PAS ERP analysis treated PAS as an ordinal single-trial predictor rather than averaging four equally weighted cells. Within each participant and exposure duration, voltage at every electrode and time sample was regressed on PAS ratings 1 to 4 with an intercept. All sequence-valid trials with a valid PAS report were included. The coefficient quantified the change in voltage, in microvolts, associated with a one-unit increase in PAS. A slope was estimated when at least two PAS levels were represented. Group-level one-sample cluster tests compared the slope maps with zero separately at each duration, and the three maps formed one Holm-corrected family. Four-level PAS averages were used only for descriptive visualisation among participants with observations at every displayed level.

A planned follow-up estimated PAS slopes separately for trials with correct and incorrect location judgements. Trials with missing discrimination or PAS responses were excluded. A participant contributed at a duration only if both accuracy strata contained at least ten trials and two PAS levels. The within-participant correct-minus-incorrect slope map directly tested a PAS by location-accuracy interaction. The 1.7- and 4.4-ms tests, each with 32 participants, formed the confirmatory family. The 6.2-ms analysis was exploratory because only 17 participants met the cell criteria.

#### Cross-condition and temporal-generalisation analyses

Three secondary decoding analyses examined the generality of expression representations. Cross-PAS generalisation trained an expression classifier on PAS 1 to 2 and tested it on PAS 3 to 4, and then reversed the direction, while keeping face identity outside the training set. Cross-duration generalisation trained at 6.2 ms and tested at 4.4 or 1.7 ms, producing train-time by test-time matrices. Finally, time-generalisation matrices for low PAS (1 to 2) and high PAS (3 to 4) were compared after exactly matching training and test counts within expression class and fold. Cells failing the prespecified trial-count requirements were retained as missing rather than estimated from sparse data.

#### Paired-label co-information validation

The posterior-centroparietal co-information analysis was repeated on a 20-ms grid from 140 to 596 ms using a participant-specific paired-label null. Within every matched emotional-neutral pair, the condition labels were randomly exchanged 200 times. The primary statistic was the observed co-information minus the participant's mean shuffled-label co-information. Positive values indicate a redundancy-dominant balance relative to this null, whereas negative values indicate a synergy-dominant balance. Duration-specific maps and direct duration contrasts were analysed in separate planned families.

#### Information shared among behavioural reports

Secondary report-level analyses used the raw location and emotional-status reports rather than accuracy. The information measures were conditioned on presented side, whether an emotional face was shown, and fearful-versus-happy block context to remove dependence directly induced by the stimulus. Measures included conditional mutual information for each report pair, PAS information about the joint reports, and conditional signed co-information. Because location and expression were entered with one compound keypress, dependencies between the two objective reports were interpreted cautiously.

#### PAS and sensor-network dynamics

Identity-grouped binary emotional-versus-neutral classifiers were fitted independently at each sensor node with 10 folds and five repeats. Held-out true-class emotion evidence was regressed continuously on PAS while controlling presented expression, face identity, target side, block, and location accuracy. At least 80 valid trials and two PAS levels were required.

PAS-modulated directed representational coupling was estimated using a PAS by source-evidence interaction to predict later target evidence while controlling current target evidence, source evidence, PAS, and the prespecified nuisance variables. Analyses covered 100 to 596 ms on an 8-ms grid with lags from 16 to 120 ms in 8-ms steps and required at least 60 trials per condition. The mean modulation across emotional and neutral trials was primary; their direct difference tested emotion specificity. For the complementary symbolic analysis, PAS 1 to 2 and PAS 3 to 4 trials were matched within expression, face identity, side, block, and location-accuracy strata. High-minus-low PAS weighted symbolic mutual information was estimated only when at least 20 trials remained in each group.

#### P1-network sensitivity and robustness analyses

The P1-EPN and P1-LPP analyses were repeated over broader component windows to assess sensitivity to the selected temporal boundaries. The broader P1 interval was 80 to 160 ms; later intervals were 260 to 360 ms for the EPN and 280 to 500 ms for the LPP. Both spatial directions were evaluated. The P1-LPP transfer-entropy effect was further examined with participant-level surrogate centring, an extended late boundary, separate fearful and happy contrasts, raw CSD estimates, voltage rather than CSD, published single-electrode definitions, and leave-one-participant-out influence analyses. These analyses assessed the statistical and representational specificity of the planned 6.2-minus-4.4-ms effect.

### SUPPLEMENTARY RESULTS

#### Model convergence and sampling diagnostics

All Bayesian models satisfied the prespecified convergence and sampling criteria (**Supplementary Table S3**; **Supplementary Figs. S1** and **S2**). For the Experiment 1 signal-detection models, maximum $\hat{R}$ was 1.0067 and minimum bulk effective sample size was 622.6. For the Experiment 1 psychometric models, maximum $\hat{R}$ was 1.0030 and minimum bulk effective sample size was 2,358.7. For the primary Experiment 2 signal-detection models, maximum $\hat{R}$ was 1.0055 and minimum bulk effective sample size was 783.1. No model contained divergent transitions or maximum-tree-depth exceedances.

#### Experiment 1 metacognitive efficiency

M-ratio estimates were generally compatible with an efficient relationship between objective performance and PAS once type-1 sensitivity became measurable, although the intervals were broad in the weakest conditions (**Supplementary Fig. S3** and **Supplementary Table S5**). At 4.4 ms, median location M-ratios ranged from 0.98 to 1.14 across orientation and expression conditions. At 6.2 ms, medians ranged from 0.79 to 1.00. Upright-minus-inverted location M-ratio contrasts were negative at 5.3 and 6.2 ms, whereas the corresponding expression-orientation contrasts remained uncertain. These efficiency estimates qualify the larger upright-face advantage in d' and meta-d': inversion changed objective and metacognitive sensitivity more consistently than it changed their ratio.

#### Secondary multivariate analyses

Trial-wise stratified decoding reproduced the main temporal ordering (**Supplementary Fig. S4**). Expression decoding exceeded chance from 516 to 592 ms at 4.4 ms ($p_{Holm}$ = .032) and from 284 to 596 ms at 6.2 ms (*p* = .0042), but not at 1.7 ms. Expression decoding that generalised to held-out identities was present only at 6.2 ms, from 348 to 596 ms (*p* = .0044). A direct comparison confirmed greater identity-generalising expression decoding at 6.2 than at 4.4 ms from 288 to 324 ms (cluster mass = 26.72, *p* = .024) and from 440 to 472 ms (cluster mass = 22.91, *p* = .040). Location decoding was reliable throughout the post-stimulus interval at 4.4 and 6.2 ms, including when identity was held out (all $p_{Holm}$ < .001).

Cross-PAS, cross-duration, and matched low-versus-high PAS temporal-generalisation analyses provided no corrected evidence that expression patterns transferred reliably across these conditions. The only corrected follow-up effect was a more negative expression-minus-location PAS-evidence slope at 4.4 ms from 248 to 596 ms ($p_{Holm}$ = .0018), indicating that PAS tracked location evidence more strongly than expression evidence during this interval.

#### Posterior-centroparietal network validation

The paired-label validation retained positive, redundancy-dominant co-information clusters at 1.7 and 4.4 ms. At 1.7 ms, significant clusters linked posterior times of 340 to 360 ms with centroparietal times of 560 to 596 ms and posterior times of 400 to 440 ms with centroparietal times of 280 to 320 ms ($p_{Holm}$ = .0076 and .0132). At 4.4 ms, clusters linked posterior times of 180 to 260 ms with centroparietal times of 560 to 596 ms and posterior times of 540 to 580 ms with centroparietal times of 420 to 480 ms (*p* = .0006 and .0042). No direct exposure-duration contrast survived correction. These results indicate local redundancy-dominant structure at the two shorter durations, but do not show that this structure changed reliably across durations.

Local multichannel emotion decoding was reliable at 6.2 ms in left-posterior, right-posterior, centroparietal, and frontocentral nodes. By contrast, no directed representational-coupling or weighted symbolic mutual-information effect provided a convergent corrected test of emotion-specific network dependence across the planned connections and durations.

#### PAS information and network dependence

Local PAS mutual information was present in both predefined nodes. Posterior PAS information was significant at 4.4 ms from 180 to 320 ms and 440 to 520 ms, and at 6.2 ms from 260 to 380 ms and 420 to 480 ms. The posterior 6.2-minus-1.7-ms contrast was significant from 420 to 480 ms. These results complement the trial-level decoding analyses by showing that subjective clarity was represented locally in both posterior and centroparietal sensor patterns.

The PAS-modulated network analyses did not yield a consistent family of corrected effects. One emotion-specific duration contrast in centroparietal-frontocentral directed representational coupling survived correction, but it was not accompanied by convergent effects in the corresponding mean coupling or symbolic-dependence analyses. It was therefore treated as exploratory rather than as evidence that PAS systematically altered directed emotion-information flow.

#### P1-EPN and P1-LPP secondary analyses

The P1 node did not show reliable local emotion mutual information, whereas the non-overlapping EPN node carried emotion information at 6.2 ms and showed a larger effect at 6.2 than at 1.7 ms. Neither the narrow nor broader P1-EPN analyses yielded corrected co-information or transfer-entropy effects. These results place the local emergence of emotion information in the later posterior interval rather than in the preceding P1 interval.

In the broader P1-LPP analysis, the emotional-minus-neutral transfer-entropy contrast from early centroparietal activity to later occipital activity was greater at 6.2 than at 4.4 ms. The planned effect linked early times from 88 to 136 ms with later times from 436 to 500 ms ($p_{Holm}$ = .0090). Surrogate-centred analysis retained a cluster spanning 88 to 112 ms and 440 to 500 ms (*p* = .0108), and extending the late boundary yielded a closely overlapping cluster (*p* = .0026). Raw-value reproduction also recovered the effect (*p* < .001). However, it was not reproduced with voltage rather than distributed CSD patterns or with the published single-electrode definitions, and the formal interaction between spatial directions was not significant. Leave-one-participant-out estimates showed that the group effect was not attributable to a single participant (**Supplementary Table S7**).

### SUPPLEMENTARY TABLES

**Supplementary Table S1 | Participant inclusion and analysis samples.** Counts are derived from the fixed inclusion manifests used by each analysis. Scheduled trial counts refer to the task design before missing-response or EEG-epoch exclusions.

| Experiment | Sample stage | Participants, n | Scheduled trials per participant | Eligibility or exclusion criterion |
|---|---|---|---|---|
| Experiment 1 | Available behavioural records | 34 | 1,120 | All participants who completed the main task |
| Experiment 1 | Included in SDT and psychometric models | 31 | 1,120 | Excluded if localisation remained at chance across all exposure durations |
| Experiment 2 | Available behavioural records | 40 | 1,680 | All participants who completed the EEG experiment |
| Experiment 2 | Behaviourally eligible | 39 | 1,680 | Fewer than 5% missing responses for every report required by the analysis |
| Experiment 2 | Included in EEG analyses | 32 | | Retained after the prespecified EEG quality-control review |

**Supplementary Table S2 | Bayesian hierarchical signal-detection model priors.** The table reports priors for condition effects and participant-level variation, together with transformations used to obtain M-ratio and meta-d'.

| Parameter | Prior or definition | Model specification |
|---|---|---|
| Condition effects on log(M) | Normal(0, 0.7) | Condition-specific fixed effects with participant-varying intercepts |
| Condition effects on d' | Normal(0, 1.5) | Condition-specific fixed effects with participant-varying intercepts |
| Condition effects on type-1 criterion | Normal(0, 1) | Condition-specific fixed effects with participant-varying intercepts |
| PAS threshold differences | Normal(-0.5, 0.75) | Condition-specific fixed effects |
| Participant-level standard deviations | Exponential(2) | Applied to participant-varying log(M), d', and criterion intercepts |
| M-ratio | exp[log(M)] | Metacognitive efficiency |
| Meta-d' | d' × M-ratio | PAS-based metacognitive sensitivity |
| Location-task sensitivity scaling | d' and meta-d' divided by √2 | Two-alternative forced-choice convention; M-ratio unchanged |

**Supplementary Table S3 | MCMC convergence and sampling diagnostics.** Reported quantities include maximum R̂, minimum bulk effective sample size, divergent transitions, and maximum-tree-depth exceedances for every fitted model. There were no divergences and all models passed.

| Experiment | Model | Task | Sample | Maximum R̂ | Minimum bulk ESS | Diagnostic outcome |
|---|---|---|---|---|---|---|
| Experiment 1 | Hierarchical SDT | Location | Primary behavioural sample | 1.0067 | 623.000 | Passed |
| Experiment 1 | Hierarchical SDT | Expression | Primary behavioural sample | 1.0026 | 1126.000 | Passed |
| Experiment 1 | Psychometric function | Location | Primary behavioural sample | 1.0030 | 2359.000 | Passed |
| Experiment 1 | Psychometric function | Expression | Primary behavioural sample | 1.0014 | 3709.000 | Passed |
| Experiment 2 | Hierarchical SDT | Location | EEG-matched sample | 1.0055 | 783.000 | Passed |
| Experiment 2 | Hierarchical SDT | Expression | EEG-matched sample | 1.0045 | 803.000 | Passed |
| Experiment 2 | Hierarchical SDT | Location | Behaviourally eligible sample | 1.0081 | 784.000 | Passed |
| Experiment 2 | Hierarchical SDT | Expression | Behaviourally eligible sample | 1.0039 | 643.000 | Passed |

**Supplementary Table S4 | Hierarchical psychometric-function priors.** The lapse parameter was constrained to remain below .15, and slopes were constrained to be positive through their log-scale parameterisation.

| Parameter | Prior | Model specification |
|---|---|---|
| Condition-level intercept | Normal(0, 1.5) | Condition-level cumulative-Gaussian intercept |
| Condition-level log-slope | Normal(log(1/1.5), 0.7) | Positive slope represented on the log scale |
| Participant intercept SD | half-Normal(0, 1.0) | Partial pooling of participant intercepts |
| Participant log-slope SD | half-Normal(0, 0.5) | Partial pooling of participant slopes |
| Condition-level lapse logit | Normal(-1.87, 1) | Group lapse parameter transformed to the interval 0 to 0.15 |
| Participant lapse-logit SD | half-Normal(0, 0.7) | Participant variation in lapse |
| Standardised participant effects | Normal(0, 1) | Non-centred participant effects |

**Supplementary Table S5 | Full Experiment 1 behavioural estimates and planned contrasts.** The table combines condition estimates and planned contrasts from the Bayesian signal-detection and hierarchical psychometric models.

| Analysis | Condition or contrast | Measure | Posterior median [95% CrI] | Posterior probability |
|---|---|---|---|---|
| Location SDT | Estimate: 0.8 ms; Upright; Neutral | log(M) | -1.710 [-2.635, -0.924] | 1.0000 |
| Location SDT | Estimate: 0.8 ms; Upright; Neutral | M-ratio | 0.181 [0.072, 0.397] | 1.0000 |
| Location SDT | Estimate: 0.8 ms; Upright; Neutral | d-prime | -0.085 [-0.273, 0.082] | 0.8320 |
| Location SDT | Estimate: 0.8 ms; Upright; Neutral | meta-d-prime | -0.014 [-0.069, 0.015] | 0.8320 |
| Location SDT | Estimate: 0.8 ms; Upright; Neutral | Type-1 criterion | 0.017 [-0.082, 0.115] | 0.6350 |
| Location SDT | Estimate: 0.8 ms; Upright; Fearful | log(M) | -0.603 [-1.442, 0.004] | 0.9740 |
| Location SDT | Estimate: 0.8 ms; Upright; Fearful | M-ratio | 0.547 [0.236, 1.004] | 0.9740 |
| Location SDT | Estimate: 0.8 ms; Upright; Fearful | d-prime | -0.009 [-0.191, 0.155] | 0.5420 |
| Location SDT | Estimate: 0.8 ms; Upright; Fearful | meta-d-prime | -0.004 [-0.120, 0.094] | 0.5420 |

| | | | | |
|---|---|---|---|---|
| Location SDT | Estimate: 0.8 ms; Upright; Fearful | Type-1 criterion | 0.059 [-0.039, 0.158] | 0.8820 |
| Location SDT | Estimate: 0.8 ms; Inverted; Neutral | log(M) | -0.806 [-1.767, -0.121] | 0.9910 |
| Location SDT | Estimate: 0.8 ms; Inverted; Neutral | M-ratio | 0.447 [0.171, 0.886] | 0.9910 |
| Location SDT | Estimate: 0.8 ms; Inverted; Neutral | d-prime | -0.109 [-0.290, 0.060] | 0.8940 |
| Location SDT | Estimate: 0.8 ms; Inverted; Neutral | meta-d-prime | -0.045 [-0.175, 0.026] | 0.8940 |
| Location SDT | Estimate: 0.8 ms; Inverted; Neutral | Type-1 criterion | 0.058 [-0.039, 0.157] | 0.8760 |
| Location SDT | Estimate: 0.8 ms; Inverted; Fearful | log(M) | -1.221 [-2.133, -0.480] | 1.0000 |
| Location SDT | Estimate: 0.8 ms; Inverted; Fearful | M-ratio | 0.295 [0.118, 0.619] | 1.0000 |
| Location SDT | Estimate: 0.8 ms; Inverted; Fearful | d-prime | -0.083 [-0.270, 0.087] | 0.8230 |
| Location SDT | Estimate: 0.8 ms; Inverted; Fearful | meta-d-prime | -0.022 [-0.104, 0.028] | 0.8230 |
| Location SDT | Estimate: 0.8 ms; Inverted; Fearful | Type-1 criterion | 0.107 [0.008, 0.205] | 0.9830 |
| Location SDT | Estimate: 1.7 ms; Upright; Neutral | log(M) | -0.343 [-0.994, 0.156] | 0.9020 |
| Location SDT | Estimate: 1.7 ms; Upright; Neutral | M-ratio | 0.709 [0.370, 1.169] | 0.9020 |
| Location SDT | Estimate: 1.7 ms; Upright; Neutral | d-prime | 0.050 [-0.134, 0.217] | 0.7100 |
| Location SDT | Estimate: 1.7 ms; Upright; Neutral | meta-d-prime | 0.033 [-0.101, 0.166] | 0.7100 |
| Location SDT | Estimate: 1.7 ms; Upright; Neutral | Type-1 criterion | 0.037 [-0.062, 0.134] | 0.7750 |
| Location SDT | Estimate: 1.7 ms; Upright; Fearful | log(M) | -0.097 [-0.655, 0.353] | 0.6490 |
| Location SDT | Estimate: 1.7 ms; Upright; Fearful | M-ratio | 0.907 [0.519, 1.423] | 0.6490 |
| Location SDT | Estimate: 1.7 ms; Upright; Fearful | d-prime | 0.014 [-0.165, 0.177] | 0.5640 |
| Location SDT | Estimate: 1.7 ms; Upright; Fearful | meta-d-prime | 0.012 [-0.158, 0.171] | 0.5640 |
| Location SDT | Estimate: 1.7 ms; Upright; Fearful | Type-1 criterion | 0.039 [-0.060, 0.137] | 0.7830 |
| Location SDT | Estimate: 1.7 ms; Inverted; Neutral | log(M) | 0.016 [-0.445, 0.420] | 0.5290 |
| Location SDT | Estimate: 1.7 ms; Inverted; Neutral | M-ratio | 1.016 [0.641, 1.522] | 0.5290 |
| Location SDT | Estimate: 1.7 ms; Inverted; Neutral | d-prime | 0.104 [-0.077, 0.270] | 0.8780 |
| Location SDT | Estimate: 1.7 ms; Inverted; Neutral | meta-d-prime | 0.103 [-0.081, 0.288] | 0.8780 |
| Location SDT | Estimate: 1.7 ms; Inverted; Neutral | Type-1 criterion | 0.048 [-0.052, 0.143] | 0.8340 |
| Location SDT | Estimate: 1.7 ms; Inverted; Fearful | log(M) | 0.061 [-0.392, 0.454] | 0.6120 |
| Location SDT | Estimate: 1.7 ms; Inverted; Fearful | M-ratio | 1.063 [0.676, 1.574] | 0.6120 |
| Location SDT | Estimate: 1.7 ms; Inverted; Fearful | d-prime | 0.078 [-0.100, 0.241] | 0.8130 |
| Location SDT | Estimate: 1.7 ms; Inverted; Fearful | meta-d-prime | 0.081 [-0.109, 0.264] | 0.8130 |
| Location SDT | Estimate: 1.7 ms; Inverted; Fearful | Type-1 criterion | 0.004 [-0.095, 0.102] | 0.5350 |
| Location SDT | Estimate: 2.6 ms; Upright; Neutral | log(M) | -0.046 [-0.412, 0.295] | 0.5970 |
| Location SDT | Estimate: 2.6 ms; Upright; Neutral | M-ratio | 0.955 [0.663, 1.343] | 0.5970 |
| Location SDT | Estimate: 2.6 ms; Upright; Neutral | d-prime | 0.466 [0.282, 0.637] | 1.0000 |
| Location SDT | Estimate: 2.6 ms; Upright; Neutral | meta-d-prime | 0.441 [0.249, 0.682] | 1.0000 |
| Location SDT | Estimate: 2.6 ms; Upright; Neutral | Type-1 criterion | -0.010 [-0.111, 0.088] | 0.5740 |
| Location SDT | Estimate: 2.6 ms; Upright; Fearful | log(M) | -0.017 [-0.383, 0.318] | 0.5370 |
| Location SDT | Estimate: 2.6 ms; Upright; Fearful | M-ratio | 0.983 [0.682, 1.374] | 0.5370 |
| Location SDT | Estimate: 2.6 ms; Upright; Fearful | d-prime | 0.473 [0.287, 0.641] | 1.0000 |
| Location SDT | Estimate: 2.6 ms; Upright; Fearful | meta-d-prime | 0.458 [0.261, 0.706] | 1.0000 |
| Location SDT | Estimate: 2.6 ms; Upright; Fearful | Type-1 criterion | 0.019 [-0.081, 0.118] | 0.6440 |
| Location SDT | Estimate: 2.6 ms; Inverted; Neutral | log(M) | 0.129 [-0.224, 0.458] | 0.7690 |
| Location SDT | Estimate: 2.6 ms; Inverted; Neutral | M-ratio | 1.138 [0.800, 1.582] | 0.7690 |
| Location SDT | Estimate: 2.6 ms; Inverted; Neutral | d-prime | 0.445 [0.259, 0.616] | 1.0000 |
| Location SDT | Estimate: 2.6 ms; Inverted; Neutral | meta-d-prime | 0.499 [0.278, 0.773] | 1.0000 |
| Location SDT | Estimate: 2.6 ms; Inverted; Neutral | Type-1 criterion | 0.089 [-0.011, 0.190] | 0.9590 |
| Location SDT | Estimate: 2.6 ms; Inverted; Fearful | log(M) | 0.154 [-0.202, 0.493] | 0.8020 |
| Location SDT | Estimate: 2.6 ms; Inverted; Fearful | M-ratio | 1.167 [0.817, 1.637] | 0.8020 |
| Location SDT | Estimate: 2.6 ms; Inverted; Fearful | d-prime | 0.357 [0.173, 0.523] | 1.0000 |
| Location SDT | Estimate: 2.6 ms; Inverted; Fearful | meta-d-prime | 0.409 [0.195, 0.666] | 1.0000 |

| | | | | |
|---|---|---|---|---|
| Location SDT | Estimate: 2.6 ms; Inverted; Fearful | Type-1 criterion | 0.056 [-0.045, 0.155] | 0.8630 |
| Location SDT | Estimate: 3.5 ms; Upright; Neutral | log(M) | -0.146 [-0.471, 0.167] | 0.8170 |
| Location SDT | Estimate: 3.5 ms; Upright; Neutral | M-ratio | 0.864 [0.624, 1.181] | 0.8170 |
| Location SDT | Estimate: 3.5 ms; Upright; Neutral | d-prime | 0.941 [0.750, 1.117] | 1.0000 |
| Location SDT | Estimate: 3.5 ms; Upright; Neutral | meta-d-prime | 0.808 [0.560, 1.135] | 1.0000 |
| Location SDT | Estimate: 3.5 ms; Upright; Neutral | Type-1 criterion | 0.083 [-0.020, 0.186] | 0.9420 |
| Location SDT | Estimate: 3.5 ms; Upright; Fearful | log(M) | 0.165 [-0.142, 0.465] | 0.8550 |
| Location SDT | Estimate: 3.5 ms; Upright; Fearful | M-ratio | 1.180 [0.868, 1.591] | 0.8550 |
| Location SDT | Estimate: 3.5 ms; Upright; Fearful | d-prime | 0.885 [0.699, 1.060] | 1.0000 |
| Location SDT | Estimate: 3.5 ms; Upright; Fearful | meta-d-prime | 1.037 [0.725, 1.446] | 1.0000 |
| Location SDT | Estimate: 3.5 ms; Upright; Fearful | Type-1 criterion | -0.065 [-0.169, 0.038] | 0.8910 |
| Location SDT | Estimate: 3.5 ms; Inverted; Neutral | log(M) | 0.265 [-0.047, 0.567] | 0.9520 |
| Location SDT | Estimate: 3.5 ms; Inverted; Neutral | M-ratio | 1.303 [0.954, 1.762] | 0.9520 |
| Location SDT | Estimate: 3.5 ms; Inverted; Neutral | d-prime | 0.723 [0.535, 0.898] | 1.0000 |
| Location SDT | Estimate: 3.5 ms; Inverted; Neutral | meta-d-prime | 0.936 [0.634, 1.323] | 1.0000 |
| Location SDT | Estimate: 3.5 ms; Inverted; Neutral | Type-1 criterion | 0.049 [-0.052, 0.149] | 0.8300 |
| Location SDT | Estimate: 3.5 ms; Inverted; Fearful | log(M) | 0.103 [-0.225, 0.417] | 0.7350 |
| Location SDT | Estimate: 3.5 ms; Inverted; Fearful | M-ratio | 1.109 [0.798, 1.517] | 0.7350 |
| Location SDT | Estimate: 3.5 ms; Inverted; Fearful | d-prime | 0.625 [0.437, 0.797] | 1.0000 |
| Location SDT | Estimate: 3.5 ms; Inverted; Fearful | meta-d-prime | 0.688 [0.443, 0.998] | 1.0000 |
| Location SDT | Estimate: 3.5 ms; Inverted; Fearful | Type-1 criterion | 0.050 [-0.053, 0.151] | 0.8290 |
| Location SDT | Estimate: 4.4 ms; Upright; Neutral | log(M) | -0.017 [-0.309, 0.270] | 0.5480 |
| Location SDT | Estimate: 4.4 ms; Upright; Neutral | M-ratio | 0.983 [0.734, 1.310] | 0.5480 |
| Location SDT | Estimate: 4.4 ms; Upright; Neutral | d-prime | 1.694 [1.490, 1.885] | 1.0000 |
| Location SDT | Estimate: 4.4 ms; Upright; Neutral | meta-d-prime | 1.661 [1.223, 2.230] | 1.0000 |
| Location SDT | Estimate: 4.4 ms; Upright; Neutral | Type-1 criterion | 0.102 [-0.013, 0.218] | 0.9590 |
| Location SDT | Estimate: 4.4 ms; Upright; Fearful | log(M) | 0.062 [-0.233, 0.344] | 0.6600 |
| Location SDT | Estimate: 4.4 ms; Upright; Fearful | M-ratio | 1.064 [0.792, 1.410] | 0.6600 |
| Location SDT | Estimate: 4.4 ms; Upright; Fearful | d-prime | 1.557 [1.357, 1.743] | 1.0000 |
| Location SDT | Estimate: 4.4 ms; Upright; Fearful | meta-d-prime | 1.650 [1.215, 2.204] | 1.0000 |
| Location SDT | Estimate: 4.4 ms; Upright; Fearful | Type-1 criterion | 0.025 [-0.085, 0.137] | 0.6660 |
| Location SDT | Estimate: 4.4 ms; Inverted; Neutral | log(M) | 0.079 [-0.221, 0.371] | 0.7000 |
| Location SDT | Estimate: 4.4 ms; Inverted; Neutral | M-ratio | 1.082 [0.802, 1.449] | 0.7000 |
| Location SDT | Estimate: 4.4 ms; Inverted; Neutral | d-prime | 1.185 [0.988, 1.364] | 1.0000 |
| Location SDT | Estimate: 4.4 ms; Inverted; Neutral | meta-d-prime | 1.276 [0.921, 1.741] | 1.0000 |
| Location SDT | Estimate: 4.4 ms; Inverted; Neutral | Type-1 criterion | 0.159 [0.051, 0.264] | 0.9980 |
| Location SDT | Estimate: 4.4 ms; Inverted; Fearful | log(M) | 0.127 [-0.170, 0.419] | 0.7960 |
| Location SDT | Estimate: 4.4 ms; Inverted; Fearful | M-ratio | 1.135 [0.844, 1.520] | 0.7960 |
| Location SDT | Estimate: 4.4 ms; Inverted; Fearful | d-prime | 1.157 [0.961, 1.338] | 1.0000 |
| Location SDT | Estimate: 4.4 ms; Inverted; Fearful | meta-d-prime | 1.311 [0.947, 1.782] | 1.0000 |
| Location SDT | Estimate: 4.4 ms; Inverted; Fearful | Type-1 criterion | 0.129 [0.020, 0.233] | 0.9910 |
| Location SDT | Estimate: 5.3 ms; Upright; Neutral | log(M) | -0.169 [-0.466, 0.115] | 0.8770 |
| Location SDT | Estimate: 5.3 ms; Upright; Neutral | M-ratio | 0.844 [0.627, 1.122] | 0.8770 |
| Location SDT | Estimate: 5.3 ms; Upright; Neutral | d-prime | 1.955 [1.746, 2.151] | 1.0000 |
| Location SDT | Estimate: 5.3 ms; Upright; Neutral | meta-d-prime | 1.647 [1.210, 2.198] | 1.0000 |
| Location SDT | Estimate: 5.3 ms; Upright; Neutral | Type-1 criterion | 0.008 [-0.113, 0.131] | 0.5510 |
| Location SDT | Estimate: 5.3 ms; Upright; Fearful | log(M) | -0.087 [-0.377, 0.195] | 0.7270 |
| Location SDT | Estimate: 5.3 ms; Upright; Fearful | M-ratio | 0.917 [0.686, 1.216] | 0.7270 |
| Location SDT | Estimate: 5.3 ms; Upright; Fearful | d-prime | 2.123 [1.912, 2.329] | 1.0000 |
| Location SDT | Estimate: 5.3 ms; Upright; Fearful | meta-d-prime | 1.942 [1.443, 2.588] | 1.0000 |
| Location SDT | Estimate: 5.3 ms; Upright; Fearful | Type-1 criterion | -0.024 [-0.152, 0.104] | 0.6440 |

| | | | | |
|---|---|---|---|---|
| Location SDT | Estimate: 5.3 ms; Inverted; Neutral | log(M) | 0.070 [-0.227, 0.359] | 0.6780 |
| Location SDT | Estimate: 5.3 ms; Inverted; Neutral | M-ratio | 1.072 [0.797, 1.431] | 0.6780 |
| Location SDT | Estimate: 5.3 ms; Inverted; Neutral | d-prime | 1.344 [1.147, 1.531] | 1.0000 |
| Location SDT | Estimate: 5.3 ms; Inverted; Neutral | meta-d-prime | 1.435 [1.047, 1.942] | 1.0000 |
| Location SDT | Estimate: 5.3 ms; Inverted; Neutral | Type-1 criterion | 0.133 [0.021, 0.242] | 0.9900 |
| Location SDT | Estimate: 5.3 ms; Inverted; Fearful | log(M) | 0.091 [-0.203, 0.379] | 0.7320 |
| Location SDT | Estimate: 5.3 ms; Inverted; Fearful | M-ratio | 1.095 [0.816, 1.461] | 0.7320 |
| Location SDT | Estimate: 5.3 ms; Inverted; Fearful | d-prime | 1.307 [1.111, 1.492] | 1.0000 |
| Location SDT | Estimate: 5.3 ms; Inverted; Fearful | meta-d-prime | 1.427 [1.043, 1.929] | 1.0000 |
| Location SDT | Estimate: 5.3 ms; Inverted; Fearful | Type-1 criterion | 0.145 [0.035, 0.252] | 0.9960 |
| Location SDT | Estimate: 6.2 ms; Upright; Neutral | log(M) | -0.230 [-0.525, 0.053] | 0.9440 |
| Location SDT | Estimate: 6.2 ms; Upright; Neutral | M-ratio | 0.794 [0.592, 1.054] | 0.9440 |
| Location SDT | Estimate: 6.2 ms; Upright; Neutral | d-prime | 2.482 [2.245, 2.713] | 1.0000 |
| Location SDT | Estimate: 6.2 ms; Upright; Neutral | meta-d-prime | 1.965 [1.464, 2.614] | 1.0000 |
| Location SDT | Estimate: 6.2 ms; Upright; Neutral | Type-1 criterion | 0.015 [-0.129, 0.156] | 0.5810 |
| Location SDT | Estimate: 6.2 ms; Upright; Fearful | log(M) | -0.181 [-0.476, 0.103] | 0.8930 |
| Location SDT | Estimate: 6.2 ms; Upright; Fearful | M-ratio | 0.834 [0.621, 1.109] | 0.8930 |
| Location SDT | Estimate: 6.2 ms; Upright; Fearful | d-prime | 2.252 [2.033, 2.468] | 1.0000 |
| Location SDT | Estimate: 6.2 ms; Upright; Fearful | meta-d-prime | 1.878 [1.392, 2.503] | 1.0000 |
| Location SDT | Estimate: 6.2 ms; Upright; Fearful | Type-1 criterion | 0.045 [-0.090, 0.175] | 0.7510 |
| Location SDT | Estimate: 6.2 ms; Inverted; Neutral | log(M) | -0.091 [-0.383, 0.195] | 0.7300 |
| Location SDT | Estimate: 6.2 ms; Inverted; Neutral | M-ratio | 0.913 [0.682, 1.216] | 0.7300 |
| Location SDT | Estimate: 6.2 ms; Inverted; Neutral | d-prime | 1.817 [1.611, 2.012] | 1.0000 |
| Location SDT | Estimate: 6.2 ms; Inverted; Neutral | meta-d-prime | 1.655 [1.226, 2.216] | 1.0000 |
| Location SDT | Estimate: 6.2 ms; Inverted; Neutral | Type-1 criterion | 0.138 [0.021, 0.257] | 0.9880 |
| Location SDT | Estimate: 6.2 ms; Inverted; Fearful | log(M) | -0.005 [-0.299, 0.279] | 0.5130 |
| Location SDT | Estimate: 6.2 ms; Inverted; Fearful | M-ratio | 0.995 [0.741, 1.322] | 0.5130 |
| Location SDT | Estimate: 6.2 ms; Inverted; Fearful | d-prime | 1.702 [1.492, 1.896] | 1.0000 |
| Location SDT | Estimate: 6.2 ms; Inverted; Fearful | meta-d-prime | 1.690 [1.245, 2.251] | 1.0000 |
| Location SDT | Estimate: 6.2 ms; Inverted; Fearful | Type-1 criterion | 0.137 [0.023, 0.252] | 0.9900 |
| Location SDT | Contrast: 0.8 ms; Neutral; upright - inverted | d-prime | 0.024 [-0.117, 0.164] | 0.6280 |
| Location SDT | Contrast: 0.8 ms; Neutral; upright - inverted | meta-d-prime | 0.027 [-0.034, 0.142] | 0.8110 |
| Location SDT | Contrast: 0.8 ms; Neutral; upright - inverted | M-ratio | -0.257 [-0.700, 0.074] | 0.9350 |
| Location SDT | Contrast: 0.8 ms; Neutral; upright - inverted | log(M) | -0.892 [-2.016, 0.294] | 0.9350 |
| Location SDT | Contrast: 0.8 ms; Neutral; upright - inverted | Type-1 criterion | -0.041 [-0.140, 0.058] | 0.7910 |
| Location SDT | Contrast: 0.8 ms; Fearful; upright - inverted | d-prime | 0.072 [-0.062, 0.209] | 0.8470 |
| Location SDT | Contrast: 0.8 ms; Fearful; upright - inverted | meta-d-prime | 0.020 [-0.071, 0.105] | 0.7100 |
| Location SDT | Contrast: 0.8 ms; Fearful; upright - inverted | M-ratio | 0.243 [-0.171, 0.724] | 0.8680 |
| Location SDT | Contrast: 0.8 ms; Fearful; upright - inverted | log(M) | 0.606 [-0.463, 1.689] | 0.8680 |
| Location SDT | Contrast: 0.8 ms; Fearful; upright - inverted | Type-1 criterion | -0.047 [-0.148, 0.054] | 0.8240 |
| Location SDT | Contrast: 0.8 ms; Upright; fearful - neutral | d-prime | 0.076 [-0.063, 0.215] | 0.8560 |
| Location SDT | Contrast: 0.8 ms; Upright; fearful - neutral | meta-d-prime | 0.013 [-0.087, 0.096] | 0.6290 |

| | | | | |
|---|---|---|---|---|
| Location SDT | Contrast: 0.8 ms; Upright; fearful - neutral | M-ratio | 0.355 [0.0004, 0.816] | 0.9750 |
| Location SDT | Contrast: 0.8 ms; Upright; fearful - neutral | log(M) | 1.092 [0.002, 2.184] | 0.9750 |
| Location SDT | Contrast: 0.8 ms; Upright; fearful - neutral | Type-1 criterion | 0.042 [-0.057, 0.144] | 0.7990 |
| Location SDT | Contrast: 0.8 ms; Inverted; fearful - neutral | d-prime | 0.026 [-0.114, 0.163] | 0.6480 |
| Location SDT | Contrast: 0.8 ms; Inverted; fearful - neutral | meta-d-prime | 0.021 [-0.050, 0.129] | 0.7300 |
| Location SDT | Contrast: 0.8 ms; Inverted; fearful - neutral | M-ratio | -0.144 [-0.610, 0.260] | 0.7640 |
| Location SDT | Contrast: 0.8 ms; Inverted; fearful - neutral | log(M) | -0.406 [-1.524, 0.741] | 0.7640 |
| Location SDT | Contrast: 0.8 ms; Inverted; fearful - neutral | Type-1 criterion | 0.048 [-0.052, 0.149] | 0.8320 |
| Location SDT | Contrast: 0.8 ms; upright - inverted | d-prime | 0.048 [-0.052, 0.149] | 0.8310 |
| Location SDT | Contrast: 0.8 ms; fearful - neutral | d-prime | 0.051 [-0.048, 0.150] | 0.8460 |
| Location SDT | Contrast: 0.8 ms; upright - inverted | meta-d-prime | 0.026 [-0.024, 0.087] | 0.8550 |
| Location SDT | Contrast: 0.8 ms; fearful - neutral | meta-d-prime | 0.018 [-0.042, 0.076] | 0.7590 |
| Location SDT | Contrast: 0.8 ms; upright - inverted | M-ratio | -0.010 [-0.316, 0.280] | 0.5270 |
| Location SDT | Contrast: 0.8 ms; fearful - neutral | M-ratio | 0.107 [-0.179, 0.412] | 0.7740 |
| Location SDT | Contrast: 0.8 ms; upright - inverted | log(M) | -0.144 [-0.927, 0.661] | 0.6400 |
| Location SDT | Contrast: 0.8 ms; fearful - neutral | log(M) | 0.346 [-0.438, 1.144] | 0.8110 |
| Location SDT | Contrast: 0.8 ms; upright - inverted | Type-1 criterion | -0.044 [-0.115, 0.027] | 0.8920 |
| Location SDT | Contrast: 0.8 ms; fearful - neutral | Type-1 criterion | 0.045 [-0.026, 0.116] | 0.8990 |
| Location SDT | Contrast: 1.7 ms; Neutral; upright - inverted | d-prime | -0.055 [-0.188, 0.077] | 0.7950 |
| Location SDT | Contrast: 1.7 ms; Neutral; upright - inverted | meta-d-prime | -0.069 [-0.204, 0.059] | 0.8680 |
| Location SDT | Contrast: 1.7 ms; Neutral; upright - inverted | M-ratio | -0.302 [-0.855, 0.205] | 0.8810 |
| Location SDT | Contrast: 1.7 ms; Neutral; upright - inverted | log(M) | -0.354 [-1.053, 0.236] | 0.8810 |
| Location SDT | Contrast: 1.7 ms; Neutral; upright - inverted | Type-1 criterion | -0.011 [-0.112, 0.090] | 0.5820 |
| Location SDT | Contrast: 1.7 ms; Fearful; upright - inverted | d-prime | -0.064 [-0.192, 0.061] | 0.8430 |
| Location SDT | Contrast: 1.7 ms; Fearful; upright - inverted | meta-d-prime | -0.069 [-0.209, 0.061] | 0.8540 |
| Location SDT | Contrast: 1.7 ms; Fearful; upright - inverted | M-ratio | -0.150 [-0.711, 0.388] | 0.7170 |
| Location SDT | Contrast: 1.7 ms; Fearful; upright - inverted | log(M) | -0.153 [-0.759, 0.384] | 0.7170 |
| Location SDT | Contrast: 1.7 ms; Fearful; upright - inverted | Type-1 criterion | 0.035 [-0.065, 0.134] | 0.7520 |
| Location SDT | Contrast: 1.7 ms; Upright; fearful - neutral | d-prime | -0.036 [-0.164, 0.094] | 0.7020 |
| Location SDT | Contrast: 1.7 ms; Upright; fearful - neutral | meta-d-prime | -0.022 [-0.146, 0.096] | 0.6510 |
| Location SDT | Contrast: 1.7 ms; Upright; fearful - neutral | M-ratio | 0.196 [-0.329, 0.754] | 0.7690 |
| Location SDT | Contrast: 1.7 ms; Upright; fearful - neutral | log(M) | 0.247 [-0.421, 0.977] | 0.7690 |
| Location SDT | Contrast: 1.7 ms; Upright; fearful - neutral | Type-1 criterion | 0.001 [-0.097, 0.102] | 0.5110 |

| | | | | |
|---|---|---|---|---|
| Location SDT | Contrast: 1.7 ms; Inverted; fearful - neutral | d-prime | -0.026 [-0.154, 0.104] | 0.6560 |
| Location SDT | Contrast: 1.7 ms; Inverted; fearful - neutral | meta-d-prime | -0.022 [-0.161, 0.113] | 0.6320 |
| Location SDT | Contrast: 1.7 ms; Inverted; fearful - neutral | M-ratio | 0.046 [-0.467, 0.565] | 0.5700 |
| Location SDT | Contrast: 1.7 ms; Inverted; fearful - neutral | log(M) | 0.045 [-0.447, 0.543] | 0.5700 |
| Location SDT | Contrast: 1.7 ms; Inverted; fearful - neutral | Type-1 criterion | -0.043 [-0.143, 0.056] | 0.8040 |
| Location SDT | Contrast: 1.7 ms; upright - inverted | d-prime | -0.060 [-0.151, 0.032] | 0.9030 |
| Location SDT | Contrast: 1.7 ms; fearful - neutral | d-prime | -0.031 [-0.122, 0.062] | 0.7450 |
| Location SDT | Contrast: 1.7 ms; upright - inverted | meta-d-prime | -0.069 [-0.170, 0.026] | 0.9280 |
| Location SDT | Contrast: 1.7 ms; fearful - neutral | meta-d-prime | -0.023 [-0.115, 0.067] | 0.6910 |
| Location SDT | Contrast: 1.7 ms; upright - inverted | M-ratio | -0.229 [-0.629, 0.148] | 0.8880 |
| Location SDT | Contrast: 1.7 ms; fearful - neutral | M-ratio | 0.120 [-0.246, 0.503] | 0.7450 |
| Location SDT | Contrast: 1.7 ms; upright - inverted | log(M) | -0.262 [-0.722, 0.155] | 0.8940 |
| Location SDT | Contrast: 1.7 ms; fearful - neutral | log(M) | 0.145 [-0.268, 0.584] | 0.7550 |
| Location SDT | Contrast: 1.7 ms; upright - inverted | Type-1 criterion | 0.012 [-0.061, 0.083] | 0.6330 |
| Location SDT | Contrast: 1.7 ms; fearful - neutral | Type-1 criterion | -0.021 [-0.091, 0.050] | 0.7210 |
| Location SDT | Contrast: 2.6 ms; Neutral; upright - inverted | d-prime | 0.021 [-0.119, 0.160] | 0.6200 |
| Location SDT | Contrast: 2.6 ms; Neutral; upright - inverted | meta-d-prime | -0.057 [-0.230, 0.096] | 0.7670 |
| Location SDT | Contrast: 2.6 ms; Neutral; upright - inverted | M-ratio | -0.177 [-0.544, 0.158] | 0.8500 |
| Location SDT | Contrast: 2.6 ms; Neutral; upright - inverted | log(M) | -0.171 [-0.503, 0.146] | 0.8500 |
| Location SDT | Contrast: 2.6 ms; Neutral; upright - inverted | Type-1 criterion | -0.099 [-0.203, 0.005] | 0.9690 |
| Location SDT | Contrast: 2.6 ms; Fearful; upright - inverted | d-prime | 0.116 [-0.026, 0.258] | 0.9470 |
| Location SDT | Contrast: 2.6 ms; Fearful; upright - inverted | meta-d-prime | 0.050 [-0.110, 0.208] | 0.7400 |
| Location SDT | Contrast: 2.6 ms; Fearful; upright - inverted | M-ratio | -0.184 [-0.571, 0.167] | 0.8430 |
| Location SDT | Contrast: 2.6 ms; Fearful; upright - inverted | log(M) | -0.172 [-0.510, 0.160] | 0.8430 |
| Location SDT | Contrast: 2.6 ms; Fearful; upright - inverted | Type-1 criterion | -0.038 [-0.140, 0.065] | 0.7620 |
| Location SDT | Contrast: 2.6 ms; Upright; fearful - neutral | d-prime | 0.006 [-0.137, 0.148] | 0.5350 |
| Location SDT | Contrast: 2.6 ms; Upright; fearful - neutral | meta-d-prime | 0.017 [-0.137, 0.175] | 0.5900 |
| Location SDT | Contrast: 2.6 ms; Upright; fearful - neutral | M-ratio | 0.026 [-0.310, 0.363] | 0.5650 |
| Location SDT | Contrast: 2.6 ms; Upright; fearful - neutral | log(M) | 0.027 [-0.316, 0.371] | 0.5650 |
| Location SDT | Contrast: 2.6 ms; Upright; fearful - neutral | Type-1 criterion | 0.029 [-0.075, 0.133] | 0.7070 |
| Location SDT | Contrast: 2.6 ms; Inverted; fearful - neutral | d-prime | -0.089 [-0.227, 0.048] | 0.8950 |
| Location SDT | Contrast: 2.6 ms; Inverted; fearful - neutral | meta-d-prime | -0.090 [-0.258, 0.061] | 0.8780 |
| Location SDT | Contrast: 2.6 ms; Inverted; fearful - neutral | M-ratio | 0.028 [-0.340, 0.405] | 0.5650 |

| | | | | |
|---|---|---|---|---|
| Location SDT | Contrast: 2.6 ms; Inverted; fearful - neutral | log(M) | 0.025 [-0.296, 0.338] | 0.5650 |
| Location SDT | Contrast: 2.6 ms; Inverted; fearful - neutral | Type-1 criterion | -0.033 [-0.135, 0.070] | 0.7370 |
| Location SDT | Contrast: 2.6 ms; upright - inverted | d-prime | 0.069 [-0.030, 0.169] | 0.9120 |
| Location SDT | Contrast: 2.6 ms; fearful - neutral | d-prime | -0.041 [-0.142, 0.055] | 0.7930 |
| Location SDT | Contrast: 2.6 ms; upright - inverted | meta-d-prime | -0.003 [-0.124, 0.108] | 0.5220 |
| Location SDT | Contrast: 2.6 ms; fearful - neutral | meta-d-prime | -0.037 [-0.151, 0.073] | 0.7490 |
| Location SDT | Contrast: 2.6 ms; upright - inverted | M-ratio | -0.179 [-0.454, 0.063] | 0.9260 |
| Location SDT | Contrast: 2.6 ms; fearful - neutral | M-ratio | 0.027 [-0.219, 0.278] | 0.5900 |
| Location SDT | Contrast: 2.6 ms; upright - inverted | log(M) | -0.171 [-0.409, 0.060] | 0.9260 |
| Location SDT | Contrast: 2.6 ms; fearful - neutral | log(M) | 0.026 [-0.203, 0.254] | 0.5860 |
| Location SDT | Contrast: 2.6 ms; upright - inverted | Type-1 criterion | -0.068 [-0.140, 0.004] | 0.9680 |
| Location SDT | Contrast: 2.6 ms; fearful - neutral | Type-1 criterion | -0.002 [-0.074, 0.070] | 0.5230 |
| Location SDT | Contrast: 3.5 ms; Neutral; upright - inverted | d-prime | 0.219 [0.063, 0.372] | 0.9970 |
| Location SDT | Contrast: 3.5 ms; Neutral; upright - inverted | meta-d-prime | -0.124 [-0.339, 0.059] | 0.9050 |
| Location SDT | Contrast: 3.5 ms; Neutral; upright - inverted | M-ratio | -0.436 [-0.759, -0.179] | 1.0000 |
| Location SDT | Contrast: 3.5 ms; Neutral; upright - inverted | log(M) | -0.410 [-0.657, -0.175] | 1.0000 |
| Location SDT | Contrast: 3.5 ms; Neutral; upright - inverted | Type-1 criterion | 0.034 [-0.074, 0.139] | 0.7310 |
| Location SDT | Contrast: 3.5 ms; Fearful; upright - inverted | d-prime | 0.260 [0.111, 0.408] | 1.0000 |
| Location SDT | Contrast: 3.5 ms; Fearful; upright - inverted | meta-d-prime | 0.347 [0.173, 0.578] | 1.0000 |
| Location SDT | Contrast: 3.5 ms; Fearful; upright - inverted | M-ratio | 0.069 [-0.203, 0.350] | 0.6960 |
| Location SDT | Contrast: 3.5 ms; Fearful; upright - inverted | log(M) | 0.061 [-0.172, 0.303] | 0.6960 |
| Location SDT | Contrast: 3.5 ms; Fearful; upright - inverted | Type-1 criterion | -0.115 [-0.221, -0.005] | 0.9800 |
| Location SDT | Contrast: 3.5 ms; Upright; fearful - neutral | d-prime | -0.057 [-0.211, 0.100] | 0.7610 |
| Location SDT | Contrast: 3.5 ms; Upright; fearful - neutral | meta-d-prime | 0.226 [0.037, 0.452] | 0.9910 |
| Location SDT | Contrast: 3.5 ms; Upright; fearful - neutral | M-ratio | 0.312 [0.083, 0.584] | 0.9970 |
| Location SDT | Contrast: 3.5 ms; Upright; fearful - neutral | log(M) | 0.311 [0.083, 0.549] | 0.9970 |
| Location SDT | Contrast: 3.5 ms; Upright; fearful - neutral | Type-1 criterion | -0.148 [-0.256, -0.039] | 0.9960 |
| Location SDT | Contrast: 3.5 ms; Inverted; fearful - neutral | d-prime | -0.097 [-0.246, 0.050] | 0.9010 |
| Location SDT | Contrast: 3.5 ms; Inverted; fearful - neutral | meta-d-prime | -0.245 [-0.452, -0.077] | 0.9980 |
| Location SDT | Contrast: 3.5 ms; Inverted; fearful - neutral | M-ratio | -0.191 [-0.512, 0.097] | 0.9050 |
| Location SDT | Contrast: 3.5 ms; Inverted; fearful - neutral | log(M) | -0.161 [-0.409, 0.081] | 0.9050 |
| Location SDT | Contrast: 3.5 ms; Inverted; fearful - neutral | Type-1 criterion | 0.0006 [-0.107, 0.105] | 0.5050 |
| Location SDT | Contrast: 3.5 ms; upright - inverted | d-prime | 0.239 [0.132, 0.345] | 1.0000 |
| Location SDT | Contrast: 3.5 ms; fearful - neutral | d-prime | -0.077 [-0.183, 0.029] | 0.9230 |
| Location SDT | Contrast: 3.5 ms; upright - inverted | meta-d-prime | 0.112 [-0.019, 0.251] | 0.9530 |

| | | | | |
|---|---|---|---|---|
| Location SDT | Contrast: 3.5 ms; fearful - neutral | meta-d-prime | -0.009 [-0.138, 0.118] | 0.5570 |
| Location SDT | Contrast: 3.5 ms; upright - inverted | M-ratio | -0.183 [-0.393, -0.0006] | 0.9750 |
| Location SDT | Contrast: 3.5 ms; fearful - neutral | M-ratio | 0.059 [-0.127, 0.253] | 0.7410 |
| Location SDT | Contrast: 3.5 ms; upright - inverted | log(M) | -0.174 [-0.342, -0.007] | 0.9790 |
| Location SDT | Contrast: 3.5 ms; fearful - neutral | log(M) | 0.074 [-0.092, 0.242] | 0.8150 |
| Location SDT | Contrast: 3.5 ms; upright - inverted | Type-1 criterion | -0.040 [-0.116, 0.036] | 0.8520 |
| Location SDT | Contrast: 3.5 ms; fearful - neutral | Type-1 criterion | -0.074 [-0.149, 0.002] | 0.9720 |
| Location SDT | Contrast: 4.4 ms; Neutral; upright - inverted | d-prime | 0.509 [0.331, 0.690] | 1.0000 |
| Location SDT | Contrast: 4.4 ms; Neutral; upright - inverted | meta-d-prime | 0.380 [0.162, 0.649] | 1.0000 |
| Location SDT | Contrast: 4.4 ms; Neutral; upright - inverted | M-ratio | -0.099 [-0.297, 0.085] | 0.8580 |
| Location SDT | Contrast: 4.4 ms; Neutral; upright - inverted | log(M) | -0.097 [-0.269, 0.082] | 0.8580 |
| Location SDT | Contrast: 4.4 ms; Neutral; upright - inverted | Type-1 criterion | -0.055 [-0.179, 0.067] | 0.8110 |
| Location SDT | Contrast: 4.4 ms; Fearful; upright - inverted | d-prime | 0.399 [0.225, 0.574] | 1.0000 |
| Location SDT | Contrast: 4.4 ms; Fearful; upright - inverted | meta-d-prime | 0.336 [0.122, 0.588] | 0.9990 |
| Location SDT | Contrast: 4.4 ms; Fearful; upright - inverted | M-ratio | -0.074 [-0.276, 0.119] | 0.7760 |
| Location SDT | Contrast: 4.4 ms; Fearful; upright - inverted | log(M) | -0.068 [-0.242, 0.107] | 0.7760 |
| Location SDT | Contrast: 4.4 ms; Fearful; upright - inverted | Type-1 criterion | -0.103 [-0.223, 0.018] | 0.9520 |
| Location SDT | Contrast: 4.4 ms; Upright; fearful - neutral | d-prime | -0.137 [-0.322, 0.046] | 0.9290 |
| Location SDT | Contrast: 4.4 ms; Upright; fearful - neutral | meta-d-prime | -0.012 [-0.242, 0.214] | 0.5400 |
| Location SDT | Contrast: 4.4 ms; Upright; fearful - neutral | M-ratio | 0.077 [-0.086, 0.254] | 0.8280 |
| Location SDT | Contrast: 4.4 ms; Upright; fearful - neutral | log(M) | 0.077 [-0.083, 0.238] | 0.8280 |
| Location SDT | Contrast: 4.4 ms; Upright; fearful - neutral | Type-1 criterion | -0.077 [-0.207, 0.053] | 0.8780 |
| Location SDT | Contrast: 4.4 ms; Inverted; fearful - neutral | d-prime | -0.028 [-0.193, 0.143] | 0.6290 |
| Location SDT | Contrast: 4.4 ms; Inverted; fearful - neutral | meta-d-prime | 0.032 [-0.171, 0.236] | 0.6260 |
| Location SDT | Contrast: 4.4 ms; Inverted; fearful - neutral | M-ratio | 0.054 [-0.156, 0.270] | 0.6960 |
| Location SDT | Contrast: 4.4 ms; Inverted; fearful - neutral | log(M) | 0.049 [-0.139, 0.233] | 0.6960 |
| Location SDT | Contrast: 4.4 ms; Inverted; fearful - neutral | Type-1 criterion | -0.030 [-0.148, 0.087] | 0.6970 |
| Location SDT | Contrast: 4.4 ms; upright - inverted | d-prime | 0.455 [0.329, 0.583] | 1.0000 |
| Location SDT | Contrast: 4.4 ms; fearful - neutral | d-prime | -0.082 [-0.207, 0.041] | 0.9030 |
| Location SDT | Contrast: 4.4 ms; upright - inverted | meta-d-prime | 0.359 [0.195, 0.564] | 1.0000 |
| Location SDT | Contrast: 4.4 ms; fearful - neutral | meta-d-prime | 0.010 [-0.144, 0.165] | 0.5550 |
| Location SDT | Contrast: 4.4 ms; upright - inverted | M-ratio | -0.086 [-0.235, 0.047] | 0.8980 |
| Location SDT | Contrast: 4.4 ms; fearful - neutral | M-ratio | 0.065 [-0.064, 0.207] | 0.8380 |
| Location SDT | Contrast: 4.4 ms; upright - inverted | log(M) | -0.082 [-0.207, 0.044] | 0.8980 |
| Location SDT | Contrast: 4.4 ms; fearful - neutral | log(M) | 0.063 [-0.059, 0.186] | 0.8400 |

| | | | | |
|---|---|---|---|---|
| Location SDT | Contrast: 4.4 ms; upright - inverted | Type-1 criterion | -0.079 [-0.166, 0.006] | 0.9660 |
| Location SDT | Contrast: 4.4 ms; fearful - neutral | Type-1 criterion | -0.054 [-0.140, 0.033] | 0.8870 |
| Location SDT | Contrast: 5.3 ms; Neutral; upright - inverted | d-prime | 0.611 [0.424, 0.799] | 1.0000 |
| Location SDT | Contrast: 5.3 ms; Neutral; upright - inverted | meta-d-prime | 0.210 [-0.022, 0.464] | 0.9620 |
| Location SDT | Contrast: 5.3 ms; Neutral; upright - inverted | M-ratio | -0.226 [-0.427, -0.063] | 0.9970 |
| Location SDT | Contrast: 5.3 ms; Neutral; upright - inverted | log(M) | -0.239 [-0.410, -0.068] | 0.9970 |
| Location SDT | Contrast: 5.3 ms; Neutral; upright - inverted | Type-1 criterion | -0.124 [-0.258, 0.009] | 0.9650 |
| Location SDT | Contrast: 5.3 ms; Fearful; upright - inverted | d-prime | 0.817 [0.620, 1.012] | 1.0000 |
| Location SDT | Contrast: 5.3 ms; Fearful; upright - inverted | meta-d-prime | 0.509 [0.270, 0.808] | 1.0000 |
| Location SDT | Contrast: 5.3 ms; Fearful; upright - inverted | M-ratio | -0.179 [-0.370, -0.014] | 0.9830 |
| Location SDT | Contrast: 5.3 ms; Fearful; upright - inverted | log(M) | -0.180 [-0.341, -0.014] | 0.9830 |
| Location SDT | Contrast: 5.3 ms; Fearful; upright - inverted | Type-1 criterion | -0.169 [-0.305, -0.033] | 0.9920 |
| Location SDT | Contrast: 5.3 ms; Upright; fearful - neutral | d-prime | 0.168 [-0.040, 0.375] | 0.9420 |
| Location SDT | Contrast: 5.3 ms; Upright; fearful - neutral | meta-d-prime | 0.293 [0.053, 0.578] | 0.9920 |
| Location SDT | Contrast: 5.3 ms; Upright; fearful - neutral | M-ratio | 0.072 [-0.066, 0.222] | 0.8480 |
| Location SDT | Contrast: 5.3 ms; Upright; fearful - neutral | log(M) | 0.082 [-0.074, 0.241] | 0.8480 |
| Location SDT | Contrast: 5.3 ms; Upright; fearful - neutral | Type-1 criterion | -0.033 [-0.179, 0.116] | 0.6710 |
| Location SDT | Contrast: 5.3 ms; Inverted; fearful - neutral | d-prime | -0.037 [-0.209, 0.137] | 0.6640 |
| Location SDT | Contrast: 5.3 ms; Inverted; fearful - neutral | meta-d-prime | -0.007 [-0.219, 0.206] | 0.5280 |
| Location SDT | Contrast: 5.3 ms; Inverted; fearful - neutral | M-ratio | 0.025 [-0.168, 0.222] | 0.6050 |
| Location SDT | Contrast: 5.3 ms; Inverted; fearful - neutral | log(M) | 0.023 [-0.149, 0.198] | 0.6050 |
| Location SDT | Contrast: 5.3 ms; Inverted; fearful - neutral | Type-1 criterion | 0.013 [-0.109, 0.131] | 0.5860 |
| Location SDT | Contrast: 5.3 ms; upright - inverted | d-prime | 0.714 [0.581, 0.848] | 1.0000 |
| Location SDT | Contrast: 5.3 ms; fearful - neutral | d-prime | 0.066 [-0.070, 0.200] | 0.8310 |
| Location SDT | Contrast: 5.3 ms; upright - inverted | meta-d-prime | 0.360 [0.184, 0.570] | 1.0000 |
| Location SDT | Contrast: 5.3 ms; fearful - neutral | meta-d-prime | 0.142 [-0.015, 0.324] | 0.9620 |
| Location SDT | Contrast: 5.3 ms; upright - inverted | M-ratio | -0.203 [-0.351, -0.084] | 1.0000 |
| Location SDT | Contrast: 5.3 ms; fearful - neutral | M-ratio | 0.047 [-0.071, 0.172] | 0.7940 |
| Location SDT | Contrast: 5.3 ms; upright - inverted | log(M) | -0.209 [-0.331, -0.090] | 1.0000 |
| Location SDT | Contrast: 5.3 ms; fearful - neutral | log(M) | 0.052 [-0.065, 0.171] | 0.8130 |
| Location SDT | Contrast: 5.3 ms; upright - inverted | Type-1 criterion | -0.147 [-0.243, -0.051] | 0.9990 |
| Location SDT | Contrast: 5.3 ms; fearful - neutral | Type-1 criterion | -0.010 [-0.106, 0.086] | 0.5810 |
| Location SDT | Contrast: 6.2 ms; Neutral; upright - inverted | d-prime | 0.666 [0.438, 0.899] | 1.0000 |

| | | | | |
|---|---|---|---|---|
| Location SDT | Contrast: 6.2 ms; Neutral; upright - inverted | meta-d-prime | 0.308 [0.060, 0.600] | 0.9920 |
| Location SDT | Contrast: 6.2 ms; Neutral; upright - inverted | M-ratio | -0.119 [-0.274, 0.018] | 0.9560 |
| Location SDT | Contrast: 6.2 ms; Neutral; upright - inverted | log(M) | -0.140 [-0.303, 0.022] | 0.9560 |
| Location SDT | Contrast: 6.2 ms; Neutral; upright - inverted | Type-1 criterion | -0.124 [-0.281, 0.035] | 0.9370 |
| Location SDT | Contrast: 6.2 ms; Fearful; upright - inverted | d-prime | 0.552 [0.337, 0.767] | 1.0000 |
| Location SDT | Contrast: 6.2 ms; Fearful; upright - inverted | meta-d-prime | 0.187 [-0.059, 0.460] | 0.9320 |
| Location SDT | Contrast: 6.2 ms; Fearful; upright - inverted | M-ratio | -0.159 [-0.329, -0.010] | 0.9820 |
| Location SDT | Contrast: 6.2 ms; Fearful; upright - inverted | log(M) | -0.176 [-0.339, -0.011] | 0.9820 |
| Location SDT | Contrast: 6.2 ms; Fearful; upright - inverted | Type-1 criterion | -0.093 [-0.238, 0.052] | 0.8940 |
| Location SDT | Contrast: 6.2 ms; Upright; fearful - neutral | d-prime | -0.229 [-0.476, 0.014] | 0.9680 |
| Location SDT | Contrast: 6.2 ms; Upright; fearful - neutral | meta-d-prime | -0.090 [-0.359, 0.169] | 0.7560 |
| Location SDT | Contrast: 6.2 ms; Upright; fearful - neutral | M-ratio | 0.040 [-0.090, 0.174] | 0.7310 |
| Location SDT | Contrast: 6.2 ms; Upright; fearful - neutral | log(M) | 0.049 [-0.110, 0.207] | 0.7310 |
| Location SDT | Contrast: 6.2 ms; Upright; fearful - neutral | Type-1 criterion | 0.030 [-0.139, 0.198] | 0.6340 |
| Location SDT | Contrast: 6.2 ms; Inverted; fearful - neutral | d-prime | -0.114 [-0.307, 0.081] | 0.8710 |
| Location SDT | Contrast: 6.2 ms; Inverted; fearful - neutral | meta-d-prime | 0.032 [-0.193, 0.265] | 0.6100 |
| Location SDT | Contrast: 6.2 ms; Inverted; fearful - neutral | M-ratio | 0.079 [-0.073, 0.245] | 0.8480 |
| Location SDT | Contrast: 6.2 ms; Inverted; fearful - neutral | log(M) | 0.084 [-0.076, 0.245] | 0.8480 |
| Location SDT | Contrast: 6.2 ms; Inverted; fearful - neutral | Type-1 criterion | -0.0006 [-0.136, 0.132] | 0.5030 |
| Location SDT | Contrast: 6.2 ms; upright - inverted | d-prime | 0.609 [0.453, 0.769] | 1.0000 |
| Location SDT | Contrast: 6.2 ms; fearful - neutral | d-prime | -0.171 [-0.328, -0.015] | 0.9850 |
| Location SDT | Contrast: 6.2 ms; upright - inverted | meta-d-prime | 0.248 [0.070, 0.460] | 0.9970 |
| Location SDT | Contrast: 6.2 ms; fearful - neutral | meta-d-prime | -0.029 [-0.202, 0.143] | 0.6350 |
| Location SDT | Contrast: 6.2 ms; upright - inverted | M-ratio | -0.140 [-0.260, -0.036] | 0.9960 |
| Location SDT | Contrast: 6.2 ms; fearful - neutral | M-ratio | 0.060 [-0.040, 0.167] | 0.8800 |
| Location SDT | Contrast: 6.2 ms; upright - inverted | log(M) | -0.159 [-0.273, -0.041] | 0.9960 |
| Location SDT | Contrast: 6.2 ms; fearful - neutral | log(M) | 0.067 [-0.046, 0.179] | 0.8760 |
| Location SDT | Contrast: 6.2 ms; upright - inverted | Type-1 criterion | -0.108 [-0.215, -0.002] | 0.9770 |
| Location SDT | Contrast: 6.2 ms; fearful - neutral | Type-1 criterion | 0.014 [-0.095, 0.123] | 0.6030 |
| Location SDT | Contrast: 1.7 ms; Upright; Neutral; 1.7 - 0.8 ms | d-prime | 0.135 [-0.002, 0.272] | 0.9730 |
| Location SDT | Contrast: 1.7 ms; Upright; Neutral; 1.7 - 0.8 ms | meta-d-prime | 0.051 [-0.065, 0.166] | 0.8340 |
| Location SDT | Contrast: 1.7 ms; Upright; Neutral; 1.7 - 0.8 ms | M-ratio | 0.519 [0.141, 0.993] | 0.9970 |

| | | | | |
|---|---|---|---|---|
| Location SDT | Contrast: 1.7 ms; Upright; Neutral; 1.7 - 0.8 ms | log(M) | 1.355 [0.389, 2.398] | 0.9970 |
| Location SDT | Contrast: 1.7 ms; Upright; Neutral; 1.7 - 0.8 ms | Type-1 criterion | 0.019 [-0.080, 0.118] | 0.6500 |
| Location SDT | Contrast: 1.7 ms; Upright; Fearful; 1.7 - 0.8 ms | d-prime | 0.024 [-0.107, 0.152] | 0.6400 |
| Location SDT | Contrast: 1.7 ms; Upright; Fearful; 1.7 - 0.8 ms | meta-d-prime | 0.018 [-0.106, 0.141] | 0.6280 |
| Location SDT | Contrast: 1.7 ms; Upright; Fearful; 1.7 - 0.8 ms | M-ratio | 0.356 [-0.193, 0.937] | 0.9000 |
| Location SDT | Contrast: 1.7 ms; Upright; Fearful; 1.7 - 0.8 ms | log(M) | 0.503 [-0.269, 1.414] | 0.9000 |
| Location SDT | Contrast: 1.7 ms; Upright; Fearful; 1.7 - 0.8 ms | Type-1 criterion | -0.022 [-0.121, 0.078] | 0.6610 |
| Location SDT | Contrast: 1.7 ms; Inverted; Neutral; 1.7 - 0.8 ms | d-prime | 0.213 [0.080, 0.347] | 0.9990 |
| Location SDT | Contrast: 1.7 ms; Inverted; Neutral; 1.7 - 0.8 ms | meta-d-prime | 0.154 [0.009, 0.312] | 0.9800 |
| Location SDT | Contrast: 1.7 ms; Inverted; Neutral; 1.7 - 0.8 ms | M-ratio | 0.559 [0.047, 1.119] | 0.9840 |
| Location SDT | Contrast: 1.7 ms; Inverted; Neutral; 1.7 - 0.8 ms | log(M) | 0.816 [0.060, 1.822] | 0.9840 |
| Location SDT | Contrast: 1.7 ms; Inverted; Neutral; 1.7 - 0.8 ms | Type-1 criterion | -0.010 [-0.112, 0.090] | 0.5830 |
| Location SDT | Contrast: 1.7 ms; Inverted; Fearful; 1.7 - 0.8 ms | d-prime | 0.161 [0.027, 0.296] | 0.9910 |
| Location SDT | Contrast: 1.7 ms; Inverted; Fearful; 1.7 - 0.8 ms | meta-d-prime | 0.107 [-0.054, 0.265] | 0.9170 |
| Location SDT | Contrast: 1.7 ms; Inverted; Fearful; 1.7 - 0.8 ms | M-ratio | 0.754 [0.306, 1.282] | 0.9990 |
| Location SDT | Contrast: 1.7 ms; Inverted; Fearful; 1.7 - 0.8 ms | log(M) | 1.269 [0.486, 2.239] | 0.9990 |
| Location SDT | Contrast: 1.7 ms; Inverted; Fearful; 1.7 - 0.8 ms | Type-1 criterion | -0.103 [-0.205, -0.003] | 0.9770 |
| Location SDT | Contrast: 2.6 ms; Upright; Neutral; 2.6 - 1.7 ms | d-prime | 0.417 [0.277, 0.556] | 1.0000 |
| Location SDT | Contrast: 2.6 ms; Upright; Neutral; 2.6 - 1.7 ms | meta-d-prime | 0.409 [0.256, 0.610] | 1.0000 |
| Location SDT | Contrast: 2.6 ms; Upright; Neutral; 2.6 - 1.7 ms | M-ratio | 0.240 [-0.191, 0.699] | 0.8620 |
| Location SDT | Contrast: 2.6 ms; Upright; Neutral; 2.6 - 1.7 ms | log(M) | 0.293 [-0.207, 0.956] | 0.8620 |
| Location SDT | Contrast: 2.6 ms; Upright; Neutral; 2.6 - 1.7 ms | Type-1 criterion | -0.046 [-0.148, 0.053] | 0.8180 |
| Location SDT | Contrast: 2.6 ms; Upright; Fearful; 2.6 - 1.7 ms | d-prime | 0.460 [0.322, 0.595] | 1.0000 |
| Location SDT | Contrast: 2.6 ms; Upright; Fearful; 2.6 - 1.7 ms | meta-d-prime | 0.450 [0.291, 0.660] | 1.0000 |
| Location SDT | Contrast: 2.6 ms; Upright; Fearful; 2.6 - 1.7 ms | M-ratio | 0.070 [-0.395, 0.535] | 0.6230 |
| Location SDT | Contrast: 2.6 ms; Upright; Fearful; 2.6 - 1.7 ms | log(M) | 0.075 [-0.372, 0.642] | 0.6230 |
| Location SDT | Contrast: 2.6 ms; Upright; Fearful; 2.6 - 1.7 ms | Type-1 criterion | -0.020 [-0.122, 0.079] | 0.6490 |
| Location SDT | Contrast: 2.6 ms; Inverted; Neutral; 2.6 - 1.7 ms | d-prime | 0.341 [0.206, 0.476] | 1.0000 |
| Location SDT | Contrast: 2.6 ms; Inverted; Neutral; 2.6 - 1.7 ms | meta-d-prime | 0.397 [0.240, 0.605] | 1.0000 |

| | | | | |
|---|---|---|---|---|
| Location SDT | Contrast: 2.6 ms; Inverted; Neutral; 2.6 - 1.7 ms | M-ratio | 0.119 [-0.331, 0.566] | 0.7020 |
| Location SDT | Contrast: 2.6 ms; Inverted; Neutral; 2.6 - 1.7 ms | log(M) | 0.111 [-0.283, 0.560] | 0.7020 |
| Location SDT | Contrast: 2.6 ms; Inverted; Neutral; 2.6 - 1.7 ms | Type-1 criterion | 0.042 [-0.061, 0.144] | 0.7860 |
| Location SDT | Contrast: 2.6 ms; Inverted; Fearful; 2.6 - 1.7 ms | d-prime | 0.278 [0.143, 0.410] | 1.0000 |
| Location SDT | Contrast: 2.6 ms; Inverted; Fearful; 2.6 - 1.7 ms | meta-d-prime | 0.330 [0.180, 0.525] | 1.0000 |
| Location SDT | Contrast: 2.6 ms; Inverted; Fearful; 2.6 - 1.7 ms | M-ratio | 0.100 [-0.360, 0.575] | 0.6710 |
| Location SDT | Contrast: 2.6 ms; Inverted; Fearful; 2.6 - 1.7 ms | log(M) | 0.091 [-0.304, 0.540] | 0.6710 |
| Location SDT | Contrast: 2.6 ms; Inverted; Fearful; 2.6 - 1.7 ms | Type-1 criterion | 0.052 [-0.049, 0.153] | 0.8440 |
| Location SDT | Contrast: 3.5 ms; Upright; Neutral; 3.5 - 2.6 ms | d-prime | 0.475 [0.325, 0.625] | 1.0000 |
| Location SDT | Contrast: 3.5 ms; Upright; Neutral; 3.5 - 2.6 ms | meta-d-prime | 0.367 [0.188, 0.586] | 1.0000 |
| Location SDT | Contrast: 3.5 ms; Upright; Neutral; 3.5 - 2.6 ms | M-ratio | -0.092 [-0.390, 0.184] | 0.7460 |
| Location SDT | Contrast: 3.5 ms; Upright; Neutral; 3.5 - 2.6 ms | log(M) | -0.103 [-0.399, 0.213] | 0.7460 |
| Location SDT | Contrast: 3.5 ms; Upright; Neutral; 3.5 - 2.6 ms | Type-1 criterion | 0.093 [-0.015, 0.199] | 0.9550 |
| Location SDT | Contrast: 3.5 ms; Upright; Fearful; 3.5 - 2.6 ms | d-prime | 0.412 [0.266, 0.560] | 1.0000 |
| Location SDT | Contrast: 3.5 ms; Upright; Fearful; 3.5 - 2.6 ms | meta-d-prime | 0.575 [0.373, 0.845] | 1.0000 |
| Location SDT | Contrast: 3.5 ms; Upright; Fearful; 3.5 - 2.6 ms | M-ratio | 0.195 [-0.094, 0.500] | 0.9100 |
| Location SDT | Contrast: 3.5 ms; Upright; Fearful; 3.5 - 2.6 ms | log(M) | 0.182 [-0.084, 0.467] | 0.9100 |
| Location SDT | Contrast: 3.5 ms; Upright; Fearful; 3.5 - 2.6 ms | Type-1 criterion | -0.084 [-0.188, 0.022] | 0.9430 |
| Location SDT | Contrast: 3.5 ms; Inverted; Neutral; 3.5 - 2.6 ms | d-prime | 0.278 [0.133, 0.424] | 1.0000 |
| Location SDT | Contrast: 3.5 ms; Inverted; Neutral; 3.5 - 2.6 ms | meta-d-prime | 0.433 [0.253, 0.668] | 1.0000 |
| Location SDT | Contrast: 3.5 ms; Inverted; Neutral; 3.5 - 2.6 ms | M-ratio | 0.164 [-0.158, 0.505] | 0.8420 |
| Location SDT | Contrast: 3.5 ms; Inverted; Neutral; 3.5 - 2.6 ms | log(M) | 0.135 [-0.125, 0.413] | 0.8420 |
| Location SDT | Contrast: 3.5 ms; Inverted; Neutral; 3.5 - 2.6 ms | Type-1 criterion | -0.040 [-0.144, 0.064] | 0.7720 |
| Location SDT | Contrast: 3.5 ms; Inverted; Fearful; 3.5 - 2.6 ms | d-prime | 0.269 [0.128, 0.410] | 1.0000 |
| Location SDT | Contrast: 3.5 ms; Inverted; Fearful; 3.5 - 2.6 ms | meta-d-prime | 0.278 [0.117, 0.471] | 1.0000 |
| Location SDT | Contrast: 3.5 ms; Inverted; Fearful; 3.5 - 2.6 ms | M-ratio | -0.058 [-0.412, 0.278] | 0.6330 |
| Location SDT | Contrast: 3.5 ms; Inverted; Fearful; 3.5 - 2.6 ms | log(M) | -0.051 [-0.343, 0.251] | 0.6330 |
| Location SDT | Contrast: 3.5 ms; Inverted; Fearful; 3.5 - 2.6 ms | Type-1 criterion | -0.007 [-0.112, 0.096] | 0.5530 |
| Location SDT | Contrast: 4.4 ms; Upright; Neutral; 4.4 - 3.5 ms | d-prime | 0.754 [0.584, 0.925] | 1.0000 |

| | | | | |
|---|---|---|---|---|
| Location SDT | Contrast: 4.4 ms; Upright; Neutral; 4.4 - 3.5 ms | meta-d-prime | 0.848 [0.579, 1.211] | 1.0000 |
| Location SDT | Contrast: 4.4 ms; Upright; Neutral; 4.4 - 3.5 ms | M-ratio | 0.118 [-0.073, 0.319] | 0.8850 |
| Location SDT | Contrast: 4.4 ms; Upright; Neutral; 4.4 - 3.5 ms | log(M) | 0.128 [-0.076, 0.343] | 0.8850 |
| Location SDT | Contrast: 4.4 ms; Upright; Neutral; 4.4 - 3.5 ms | Type-1 criterion | 0.019 [-0.101, 0.143] | 0.6200 |
| Location SDT | Contrast: 4.4 ms; Upright; Fearful; 4.4 - 3.5 ms | d-prime | 0.673 [0.507, 0.838] | 1.0000 |
| Location SDT | Contrast: 4.4 ms; Upright; Fearful; 4.4 - 3.5 ms | meta-d-prime | 0.607 [0.376, 0.900] | 1.0000 |
| Location SDT | Contrast: 4.4 ms; Upright; Fearful; 4.4 - 3.5 ms | M-ratio | -0.118 [-0.344, 0.089] | 0.8700 |
| Location SDT | Contrast: 4.4 ms; Upright; Fearful; 4.4 - 3.5 ms | log(M) | -0.105 [-0.289, 0.082] | 0.8700 |
| Location SDT | Contrast: 4.4 ms; Upright; Fearful; 4.4 - 3.5 ms | Type-1 criterion | 0.090 [-0.029, 0.208] | 0.9350 |
| Location SDT | Contrast: 4.4 ms; Inverted; Neutral; 4.4 - 3.5 ms | d-prime | 0.462 [0.301, 0.623] | 1.0000 |
| Location SDT | Contrast: 4.4 ms; Inverted; Neutral; 4.4 - 3.5 ms | meta-d-prime | 0.338 [0.144, 0.575] | 1.0000 |
| Location SDT | Contrast: 4.4 ms; Inverted; Neutral; 4.4 - 3.5 ms | M-ratio | -0.220 [-0.501, 0.025] | 0.9600 |
| Location SDT | Contrast: 4.4 ms; Inverted; Neutral; 4.4 - 3.5 ms | log(M) | -0.186 [-0.392, 0.022] | 0.9600 |
| Location SDT | Contrast: 4.4 ms; Inverted; Neutral; 4.4 - 3.5 ms | Type-1 criterion | 0.109 [-0.005, 0.222] | 0.9700 |
| Location SDT | Contrast: 4.4 ms; Inverted; Fearful; 4.4 - 3.5 ms | d-prime | 0.533 [0.373, 0.689] | 1.0000 |
| Location SDT | Contrast: 4.4 ms; Inverted; Fearful; 4.4 - 3.5 ms | meta-d-prime | 0.619 [0.404, 0.903] | 1.0000 |
| Location SDT | Contrast: 4.4 ms; Inverted; Fearful; 4.4 - 3.5 ms | M-ratio | 0.025 [-0.234, 0.289] | 0.5760 |
| Location SDT | Contrast: 4.4 ms; Inverted; Fearful; 4.4 - 3.5 ms | log(M) | 0.022 [-0.199, 0.256] | 0.5760 |
| Location SDT | Contrast: 4.4 ms; Inverted; Fearful; 4.4 - 3.5 ms | Type-1 criterion | 0.078 [-0.031, 0.190] | 0.9200 |
| Location SDT | Contrast: 5.3 ms; Upright; Neutral; 5.3 - 4.4 ms | d-prime | 0.261 [0.066, 0.452] | 0.9970 |
| Location SDT | Contrast: 5.3 ms; Upright; Neutral; 5.3 - 4.4 ms | meta-d-prime | -0.016 [-0.254, 0.219] | 0.5520 |
| Location SDT | Contrast: 5.3 ms; Upright; Neutral; 5.3 - 4.4 ms | M-ratio | -0.138 [-0.303, 0.006] | 0.9690 |
| Location SDT | Contrast: 5.3 ms; Upright; Neutral; 5.3 - 4.4 ms | log(M) | -0.153 [-0.316, 0.006] | 0.9690 |
| Location SDT | Contrast: 5.3 ms; Upright; Neutral; 5.3 - 4.4 ms | Type-1 criterion | -0.094 [-0.231, 0.043] | 0.9100 |
| Location SDT | Contrast: 5.3 ms; Upright; Fearful; 5.3 - 4.4 ms | d-prime | 0.567 [0.367, 0.763] | 1.0000 |
| Location SDT | Contrast: 5.3 ms; Upright; Fearful; 5.3 - 4.4 ms | meta-d-prime | 0.288 [0.055, 0.559] | 0.9920 |
| Location SDT | Contrast: 5.3 ms; Upright; Fearful; 5.3 - 4.4 ms | M-ratio | -0.145 [-0.320, 0.011] | 0.9670 |
| Location SDT | Contrast: 5.3 ms; Upright; Fearful; 5.3 - 4.4 ms | log(M) | -0.148 [-0.303, 0.011] | 0.9670 |
| Location SDT | Contrast: 5.3 ms; Upright; Fearful; 5.3 - 4.4 ms | Type-1 criterion | -0.049 [-0.188, 0.089] | 0.7560 |

| | | | | |
|---|---|---|---|---|
| Location SDT | Contrast: 5.3 ms; Inverted; Neutral; 5.3 - 4.4 ms | d-prime | 0.159 [-0.016, 0.331] | 0.9650 |
| Location SDT | Contrast: 5.3 ms; Inverted; Neutral; 5.3 - 4.4 ms | meta-d-prime | 0.156 [-0.050, 0.385] | 0.9310 |
| Location SDT | Contrast: 5.3 ms; Inverted; Neutral; 5.3 - 4.4 ms | M-ratio | -0.011 [-0.216, 0.197] | 0.5440 |
| Location SDT | Contrast: 5.3 ms; Inverted; Neutral; 5.3 - 4.4 ms | log(M) | -0.010 [-0.194, 0.179] | 0.5440 |
| Location SDT | Contrast: 5.3 ms; Inverted; Neutral; 5.3 - 4.4 ms | Type-1 criterion | -0.026 [-0.143, 0.094] | 0.6660 |
| Location SDT | Contrast: 5.3 ms; Inverted; Fearful; 5.3 - 4.4 ms | d-prime | 0.149 [-0.021, 0.321] | 0.9570 |
| Location SDT | Contrast: 5.3 ms; Inverted; Fearful; 5.3 - 4.4 ms | meta-d-prime | 0.116 [-0.084, 0.333] | 0.8760 |
| Location SDT | Contrast: 5.3 ms; Inverted; Fearful; 5.3 - 4.4 ms | M-ratio | -0.040 [-0.248, 0.164] | 0.6490 |
| Location SDT | Contrast: 5.3 ms; Inverted; Fearful; 5.3 - 4.4 ms | log(M) | -0.036 [-0.214, 0.145] | 0.6490 |
| Location SDT | Contrast: 5.3 ms; Inverted; Fearful; 5.3 - 4.4 ms | Type-1 criterion | 0.017 [-0.100, 0.131] | 0.6110 |
| Location SDT | Contrast: 6.2 ms; Upright; Neutral; 6.2 - 5.3 ms | d-prime | 0.527 [0.295, 0.759] | 1.0000 |
| Location SDT | Contrast: 6.2 ms; Upright; Neutral; 6.2 - 5.3 ms | meta-d-prime | 0.319 [0.067, 0.604] | 0.9940 |
| Location SDT | Contrast: 6.2 ms; Upright; Neutral; 6.2 - 5.3 ms | M-ratio | -0.049 [-0.188, 0.083] | 0.7710 |
| Location SDT | Contrast: 6.2 ms; Upright; Neutral; 6.2 - 5.3 ms | log(M) | -0.061 [-0.223, 0.101] | 0.7710 |
| Location SDT | Contrast: 6.2 ms; Upright; Neutral; 6.2 - 5.3 ms | Type-1 criterion | 0.007 [-0.156, 0.168] | 0.5310 |
| Location SDT | Contrast: 6.2 ms; Upright; Fearful; 6.2 - 5.3 ms | d-prime | 0.130 [-0.090, 0.353] | 0.8790 |
| Location SDT | Contrast: 6.2 ms; Upright; Fearful; 6.2 - 5.3 ms | meta-d-prime | -0.064 [-0.325, 0.189] | 0.6910 |
| Location SDT | Contrast: 6.2 ms; Upright; Fearful; 6.2 - 5.3 ms | M-ratio | -0.082 [-0.225, 0.052] | 0.8830 |
| Location SDT | Contrast: 6.2 ms; Upright; Fearful; 6.2 - 5.3 ms | log(M) | -0.094 [-0.246, 0.060] | 0.8830 |
| Location SDT | Contrast: 6.2 ms; Upright; Fearful; 6.2 - 5.3 ms | Type-1 criterion | 0.069 [-0.089, 0.223] | 0.8090 |
| Location SDT | Contrast: 6.2 ms; Inverted; Neutral; 6.2 - 5.3 ms | d-prime | 0.472 [0.286, 0.658] | 1.0000 |
| Location SDT | Contrast: 6.2 ms; Inverted; Neutral; 6.2 - 5.3 ms | meta-d-prime | 0.218 [0.002, 0.463] | 0.9760 |
| Location SDT | Contrast: 6.2 ms; Inverted; Neutral; 6.2 - 5.3 ms | M-ratio | -0.156 [-0.350, 0.013] | 0.9660 |
| Location SDT | Contrast: 6.2 ms; Inverted; Neutral; 6.2 - 5.3 ms | log(M) | -0.159 [-0.332, 0.014] | 0.9660 |
| Location SDT | Contrast: 6.2 ms; Inverted; Neutral; 6.2 - 5.3 ms | Type-1 criterion | 0.006 [-0.125, 0.137] | 0.5340 |
| Location SDT | Contrast: 6.2 ms; Inverted; Fearful; 6.2 - 5.3 ms | d-prime | 0.396 [0.214, 0.577] | 1.0000 |
| Location SDT | Contrast: 6.2 ms; Inverted; Fearful; 6.2 - 5.3 ms | meta-d-prime | 0.258 [0.042, 0.508] | 0.9900 |
| Location SDT | Contrast: 6.2 ms; Inverted; Fearful; 6.2 - 5.3 ms | M-ratio | -0.102 [-0.293, 0.073] | 0.8700 |
| Location SDT | Contrast: 6.2 ms; Inverted; Fearful; 6.2 - 5.3 ms | log(M) | -0.098 [-0.265, 0.069] | 0.8700 |

| | | | | |
|---|---|---|---|---|
| Location SDT | Contrast: 6.2 ms; Inverted; Fearful; 6.2 - 5.3 ms | Type-1 criterion | -0.007 [-0.131, 0.118] | 0.5470 |
| Expression SDT | Estimate: 0.8 ms; Upright | log(M) | -0.651 [-1.749, 0.303] | 0.9080 |
| Expression SDT | Estimate: 0.8 ms; Upright | M-ratio | 0.521 [0.174, 1.354] | 0.9080 |
| Expression SDT | Estimate: 0.8 ms; Upright | d-prime | 0.016 [-0.084, 0.119] | 0.6190 |
| Expression SDT | Estimate: 0.8 ms; Upright | meta-d-prime | 0.007 [-0.063, 0.071] | 0.6190 |
| Expression SDT | Estimate: 0.8 ms; Upright | Type-1 criterion | 0.218 [0.083, 0.349] | 0.9990 |
| Expression SDT | Estimate: 0.8 ms; Inverted | log(M) | -0.687 [-1.776, 0.223] | 0.9280 |
| Expression SDT | Estimate: 0.8 ms; Inverted | M-ratio | 0.503 [0.169, 1.250] | 0.9280 |
| Expression SDT | Estimate: 0.8 ms; Inverted | d-prime | -0.023 [-0.126, 0.080] | 0.6740 |
| Expression SDT | Estimate: 0.8 ms; Inverted | meta-d-prime | -0.010 [-0.080, 0.046] | 0.6740 |
| Expression SDT | Estimate: 0.8 ms; Inverted | Type-1 criterion | 0.349 [0.215, 0.483] | 1.0000 |
| Expression SDT | Estimate: 1.7 ms; Upright | log(M) | -0.580 [-1.659, 0.301] | 0.8930 |
| Expression SDT | Estimate: 1.7 ms; Upright | M-ratio | 0.560 [0.190, 1.351] | 0.8930 |
| Expression SDT | Estimate: 1.7 ms; Upright | d-prime | -0.073 [-0.170, 0.024] | 0.9290 |
| Expression SDT | Estimate: 1.7 ms; Upright | meta-d-prime | -0.038 [-0.133, 0.012] | 0.9290 |
| Expression SDT | Estimate: 1.7 ms; Upright | Type-1 criterion | 0.216 [0.081, 0.347] | 0.9990 |
| Expression SDT | Estimate: 1.7 ms; Inverted | log(M) | -0.745 [-1.807, 0.178] | 0.9400 |
| Expression SDT | Estimate: 1.7 ms; Inverted | M-ratio | 0.475 [0.164, 1.195] | 0.9400 |
| Expression SDT | Estimate: 1.7 ms; Inverted | d-prime | -0.014 [-0.113, 0.094] | 0.6070 |
| Expression SDT | Estimate: 1.7 ms; Inverted | meta-d-prime | -0.006 [-0.083, 0.041] | 0.6070 |
| Expression SDT | Estimate: 1.7 ms; Inverted | Type-1 criterion | 0.420 [0.285, 0.553] | 1.0000 |
| Expression SDT | Estimate: 2.6 ms; Upright | log(M) | -0.752 [-1.910, 0.302] | 0.9200 |
| Expression SDT | Estimate: 2.6 ms; Upright | M-ratio | 0.471 [0.148, 1.353] | 0.9200 |
| Expression SDT | Estimate: 2.6 ms; Upright | d-prime | -0.027 [-0.125, 0.083] | 0.6860 |
| Expression SDT | Estimate: 2.6 ms; Upright | meta-d-prime | -0.011 [-0.109, 0.032] | 0.6860 |
| Expression SDT | Estimate: 2.6 ms; Upright | Type-1 criterion | 0.205 [0.071, 0.337] | 0.9980 |
| Expression SDT | Estimate: 2.6 ms; Inverted | log(M) | -0.604 [-1.659, 0.306] | 0.9020 |
| Expression SDT | Estimate: 2.6 ms; Inverted | M-ratio | 0.547 [0.190, 1.357] | 0.9020 |
| Expression SDT | Estimate: 2.6 ms; Inverted | d-prime | 0.040 [-0.059, 0.147] | 0.7780 |

| | | | | |
|---|---|---|---|---|
| Expression SDT | Estimate: 2.6 ms; Inverted | meta-d-prime | 0.019 [-0.044, 0.087] | 0.7780 |
| Expression SDT | Estimate: 2.6 ms; Inverted | Type-1 criterion | 0.309 [0.173, 0.442] | 1.0000 |
| Expression SDT | Estimate: 3.5 ms; Upright | log(M) | -0.050 [-0.777, 0.595] | 0.5600 |
| Expression SDT | Estimate: 3.5 ms; Upright | M-ratio | 0.951 [0.460, 1.813] | 0.5600 |
| Expression SDT | Estimate: 3.5 ms; Upright | d-prime | 0.134 [0.044, 0.238] | 0.9980 |
| Expression SDT | Estimate: 3.5 ms; Upright | meta-d-prime | 0.125 [0.043, 0.237] | 0.9980 |
| Expression SDT | Estimate: 3.5 ms; Upright | Type-1 criterion | 0.178 [0.045, 0.311] | 0.9950 |
| Expression SDT | Estimate: 3.5 ms; Inverted | log(M) | -0.337 [-1.373, 0.421] | 0.7980 |
| Expression SDT | Estimate: 3.5 ms; Inverted | M-ratio | 0.714 [0.253, 1.524] | 0.7980 |
| Expression SDT | Estimate: 3.5 ms; Inverted | d-prime | 0.033 [-0.059, 0.127] | 0.7570 |
| Expression SDT | Estimate: 3.5 ms; Inverted | meta-d-prime | 0.021 [-0.045, 0.101] | 0.7570 |
| Expression SDT | Estimate: 3.5 ms; Inverted | Type-1 criterion | 0.377 [0.242, 0.509] | 1.0000 |
| Expression SDT | Estimate: 4.4 ms; Upright | log(M) | -0.212 [-0.783, 0.319] | 0.7790 |
| Expression SDT | Estimate: 4.4 ms; Upright | M-ratio | 0.809 [0.457, 1.376] | 0.7790 |
| Expression SDT | Estimate: 4.4 ms; Upright | d-prime | 0.267 [0.162, 0.377] | 1.0000 |
| Expression SDT | Estimate: 4.4 ms; Upright | meta-d-prime | 0.214 [0.121, 0.344] | 1.0000 |
| Expression SDT | Estimate: 4.4 ms; Upright | Type-1 criterion | 0.142 [0.009, 0.275] | 0.9810 |
| Expression SDT | Estimate: 4.4 ms; Inverted | log(M) | -0.363 [-1.264, 0.342] | 0.8340 |
| Expression SDT | Estimate: 4.4 ms; Inverted | M-ratio | 0.695 [0.283, 1.407] | 0.8340 |
| Expression SDT | Estimate: 4.4 ms; Inverted | d-prime | 0.104 [0.012, 0.207] | 0.9870 |
| Expression SDT | Estimate: 4.4 ms; Inverted | meta-d-prime | 0.069 [0.008, 0.160] | 0.9870 |
| Expression SDT | Estimate: 4.4 ms; Inverted | Type-1 criterion | 0.332 [0.198, 0.465] | 1.0000 |
| Expression SDT | Estimate: 5.3 ms; Upright | log(M) | 0.012 [-0.433, 0.448] | 0.5210 |
| Expression SDT | Estimate: 5.3 ms; Upright | M-ratio | 1.012 [0.649, 1.565] | 0.5210 |
| Expression SDT | Estimate: 5.3 ms; Upright | d-prime | 0.382 [0.276, 0.490] | 1.0000 |
| Expression SDT | Estimate: 5.3 ms; Upright | meta-d-prime | 0.385 [0.252, 0.569] | 1.0000 |
| Expression SDT | Estimate: 5.3 ms; Upright | Type-1 criterion | 0.046 [-0.087, 0.178] | 0.7510 |
| Expression SDT | Estimate: 5.3 ms; Inverted | log(M) | -0.017 [-0.777, 0.662] | 0.5210 |
| Expression SDT | Estimate: 5.3 ms; Inverted | M-ratio | 0.983 [0.460, 1.939] | 0.5210 |

| | | | | |
|---|---|---|---|---|
| Expression SDT | Estimate: 5.3 ms; Inverted | d-prime | 0.116 [0.017, 0.221] | 0.9910 |
| Expression SDT | Estimate: 5.3 ms; Inverted | meta-d-prime | 0.110 [0.021, 0.215] | 0.9910 |
| Expression SDT | Estimate: 5.3 ms; Inverted | Type-1 criterion | 0.312 [0.178, 0.444] | 1.0000 |
| Expression SDT | Estimate: 6.2 ms; Upright | log(M) | -0.330 [-0.808, 0.106] | 0.9290 |
| Expression SDT | Estimate: 6.2 ms; Upright | M-ratio | 0.719 [0.446, 1.111] | 0.9290 |
| Expression SDT | Estimate: 6.2 ms; Upright | d-prime | 0.458 [0.347, 0.567] | 1.0000 |
| Expression SDT | Estimate: 6.2 ms; Upright | meta-d-prime | 0.327 [0.207, 0.488] | 1.0000 |
| Expression SDT | Estimate: 6.2 ms; Upright | Type-1 criterion | -0.049 [-0.182, 0.084] | 0.7710 |
| Expression SDT | Estimate: 6.2 ms; Inverted | log(M) | -0.128 [-0.754, 0.453] | 0.6680 |
| Expression SDT | Estimate: 6.2 ms; Inverted | M-ratio | 0.880 [0.471, 1.573] | 0.6680 |
| Expression SDT | Estimate: 6.2 ms; Inverted | d-prime | 0.219 [0.109, 0.331] | 1.0000 |
| Expression SDT | Estimate: 6.2 ms; Inverted | meta-d-prime | 0.188 [0.101, 0.308] | 1.0000 |
| Expression SDT | Estimate: 6.2 ms; Inverted | Type-1 criterion | 0.272 [0.140, 0.403] | 1.0000 |
| Expression SDT | Contrast: 0.8 ms; upright - inverted | d-prime | 0.039 [-0.092, 0.174] | 0.7160 |
| Expression SDT | Contrast: 0.8 ms; upright - inverted | meta-d-prime | 0.019 [-0.063, 0.102] | 0.6990 |
| Expression SDT | Contrast: 0.8 ms; upright - inverted | M-ratio | 0.021 [-0.770, 0.869] | 0.5230 |
| Expression SDT | Contrast: 0.8 ms; upright - inverted | log(M) | 0.043 [-1.355, 1.429] | 0.5230 |
| Expression SDT | Contrast: 0.8 ms; upright - inverted | Type-1 criterion | -0.131 [-0.204, -0.058] | 1.0000 |
| Expression SDT | Contrast: 1.7 ms; upright - inverted | d-prime | -0.058 [-0.194, 0.069] | 0.8160 |
| Expression SDT | Contrast: 1.7 ms; upright - inverted | meta-d-prime | -0.032 [-0.132, 0.054] | 0.7990 |
| Expression SDT | Contrast: 1.7 ms; upright - inverted | M-ratio | 0.081 [-0.706, 0.904] | 0.5930 |
| Expression SDT | Contrast: 1.7 ms; upright - inverted | log(M) | 0.163 [-1.246, 1.526] | 0.5930 |
| Expression SDT | Contrast: 1.7 ms; upright - inverted | Type-1 criterion | -0.204 [-0.277, -0.130] | 1.0000 |
| Expression SDT | Contrast: 2.6 ms; upright - inverted | d-prime | -0.065 [-0.202, 0.071] | 0.8320 |
| Expression SDT | Contrast: 2.6 ms; upright - inverted | meta-d-prime | -0.035 [-0.139, 0.039] | 0.8350 |
| Expression SDT | Contrast: 2.6 ms; upright - inverted | M-ratio | -0.069 [-0.884, 0.789] | 0.5820 |
| Expression SDT | Contrast: 2.6 ms; upright - inverted | log(M) | -0.141 [-1.528, 1.242] | 0.5820 |
| Expression SDT | Contrast: 2.6 ms; upright - inverted | Type-1 criterion | -0.105 [-0.177, -0.033] | 0.9980 |
| Expression SDT | Contrast: 3.5 ms; upright - inverted | d-prime | 0.103 [-0.016, 0.228] | 0.9520 |

| | | | | |
|---|---|---|---|---|
| Expression SDT | Contrast: 3.5 ms; upright - inverted | meta-d-prime | 0.103 [0.003, 0.224] | 0.9780 |
| Expression SDT | Contrast: 3.5 ms; upright - inverted | M-ratio | 0.239 [-0.646, 1.164] | 0.7200 |
| Expression SDT | Contrast: 3.5 ms; upright - inverted | log(M) | 0.296 [-0.707, 1.487] | 0.7200 |
| Expression SDT | Contrast: 3.5 ms; upright - inverted | Type-1 criterion | -0.199 [-0.272, -0.127] | 1.0000 |
| Expression SDT | Contrast: 4.4 ms; upright - inverted | d-prime | 0.162 [0.029, 0.298] | 0.9910 |
| Expression SDT | Contrast: 4.4 ms; upright - inverted | meta-d-prime | 0.143 [0.032, 0.275] | 0.9940 |
| Expression SDT | Contrast: 4.4 ms; upright - inverted | M-ratio | 0.114 [-0.634, 0.769] | 0.6350 |
| Expression SDT | Contrast: 4.4 ms; upright - inverted | log(M) | 0.155 [-0.704, 1.129] | 0.6350 |
| Expression SDT | Contrast: 4.4 ms; upright - inverted | Type-1 criterion | -0.189 [-0.262, -0.116] | 1.0000 |
| Expression SDT | Contrast: 5.3 ms; upright - inverted | d-prime | 0.266 [0.128, 0.402] | 1.0000 |
| Expression SDT | Contrast: 5.3 ms; upright - inverted | meta-d-prime | 0.274 [0.135, 0.454] | 1.0000 |
| Expression SDT | Contrast: 5.3 ms; upright - inverted | M-ratio | 0.026 [-0.898, 0.714] | 0.5280 |
| Expression SDT | Contrast: 5.3 ms; upright - inverted | log(M) | 0.026 [-0.697, 0.843] | 0.5280 |
| Expression SDT | Contrast: 5.3 ms; upright - inverted | Type-1 criterion | -0.267 [-0.338, -0.195] | 1.0000 |
| Expression SDT | Contrast: 6.2 ms; upright - inverted | d-prime | 0.238 [0.091, 0.385] | 0.9990 |
| Expression SDT | Contrast: 6.2 ms; upright - inverted | meta-d-prime | 0.137 [0.007, 0.288] | 0.9820 |
| Expression SDT | Contrast: 6.2 ms; upright - inverted | M-ratio | -0.158 [-0.827, 0.336] | 0.7280 |
| Expression SDT | Contrast: 6.2 ms; upright - inverted | log(M) | -0.200 [-0.853, 0.464] | 0.7280 |
| Expression SDT | Contrast: 6.2 ms; upright - inverted | Type-1 criterion | -0.321 [-0.393, -0.249] | 1.0000 |
| Expression SDT | Contrast: 1.7 ms; Upright; 1.7 - 0.8 ms | d-prime | -0.088 [-0.220, 0.039] | 0.9130 |
| Expression SDT | Contrast: 1.7 ms; Upright; 1.7 - 0.8 ms | meta-d-prime | -0.048 [-0.150, 0.033] | 0.8900 |
| Expression SDT | Contrast: 1.7 ms; Upright; 1.7 - 0.8 ms | M-ratio | 0.033 [-0.829, 0.859] | 0.5370 |
| Expression SDT | Contrast: 1.7 ms; Upright; 1.7 - 0.8 ms | log(M) | 0.061 [-1.351, 1.450] | 0.5370 |
| Expression SDT | Contrast: 1.7 ms; Upright; 1.7 - 0.8 ms | Type-1 criterion | -0.002 [-0.075, 0.069] | 0.5230 |
| Expression SDT | Contrast: 1.7 ms; Inverted; 1.7 - 0.8 ms | d-prime | 0.009 [-0.125, 0.151] | 0.5540 |
| Expression SDT | Contrast: 1.7 ms; Inverted; 1.7 - 0.8 ms | meta-d-prime | 0.003 [-0.084, 0.081] | 0.5360 |
| Expression SDT | Contrast: 1.7 ms; Inverted; 1.7 - 0.8 ms | M-ratio | -0.025 [-0.794, 0.726] | 0.5310 |
| Expression SDT | Contrast: 1.7 ms; Inverted; 1.7 - 0.8 ms | log(M) | -0.056 [-1.417, 1.332] | 0.5310 |
| Expression SDT | Contrast: 1.7 ms; Inverted; 1.7 - 0.8 ms | Type-1 criterion | 0.071 [-0.004, 0.143] | 0.9690 |

| | | | | |
|---|---|---|---|---|
| Expression SDT | Contrast: 2.6 ms; Upright; 2.6 - 1.7 ms | d-prime | 0.047 [-0.082, 0.184] | 0.7590 |
| Expression SDT | Contrast: 2.6 ms; Upright; 2.6 - 1.7 ms | meta-d-prime | 0.026 [-0.076, 0.125] | 0.7330 |
| Expression SDT | Contrast: 2.6 ms; Upright; 2.6 - 1.7 ms | M-ratio | -0.086 [-0.901, 0.806] | 0.5900 |
| Expression SDT | Contrast: 2.6 ms; Upright; 2.6 - 1.7 ms | log(M) | -0.175 [-1.595, 1.288] | 0.5900 |
| Expression SDT | Contrast: 2.6 ms; Upright; 2.6 - 1.7 ms | Type-1 criterion | -0.011 [-0.083, 0.061] | 0.6190 |
| Expression SDT | Contrast: 2.6 ms; Inverted; 2.6 - 1.7 ms | d-prime | 0.054 [-0.080, 0.188] | 0.7860 |
| Expression SDT | Contrast: 2.6 ms; Inverted; 2.6 - 1.7 ms | meta-d-prime | 0.028 [-0.045, 0.118] | 0.7900 |
| Expression SDT | Contrast: 2.6 ms; Inverted; 2.6 - 1.7 ms | M-ratio | 0.071 [-0.687, 0.892] | 0.5840 |
| Expression SDT | Contrast: 2.6 ms; Inverted; 2.6 - 1.7 ms | log(M) | 0.145 [-1.196, 1.503] | 0.5840 |
| Expression SDT | Contrast: 2.6 ms; Inverted; 2.6 - 1.7 ms | Type-1 criterion | -0.110 [-0.184, -0.036] | 0.9990 |
| Expression SDT | Contrast: 3.5 ms; Upright; 3.5 - 2.6 ms | d-prime | 0.161 [0.030, 0.296] | 0.9920 |
| Expression SDT | Contrast: 3.5 ms; Upright; 3.5 - 2.6 ms | meta-d-prime | 0.143 [0.049, 0.275] | 0.9990 |
| Expression SDT | Contrast: 3.5 ms; Upright; 3.5 - 2.6 ms | M-ratio | 0.463 [-0.484, 1.341] | 0.8720 |
| Expression SDT | Contrast: 3.5 ms; Upright; 3.5 - 2.6 ms | log(M) | 0.703 [-0.535, 1.970] | 0.8720 |
| Expression SDT | Contrast: 3.5 ms; Upright; 3.5 - 2.6 ms | Type-1 criterion | -0.026 [-0.099, 0.045] | 0.7650 |
| Expression SDT | Contrast: 3.5 ms; Inverted; 3.5 - 2.6 ms | d-prime | -0.007 [-0.140, 0.121] | 0.5440 |
| Expression SDT | Contrast: 3.5 ms; Inverted; 3.5 - 2.6 ms | meta-d-prime | 0.002 [-0.084, 0.095] | 0.5220 |
| Expression SDT | Contrast: 3.5 ms; Inverted; 3.5 - 2.6 ms | M-ratio | 0.147 [-0.724, 1.000] | 0.6520 |
| Expression SDT | Contrast: 3.5 ms; Inverted; 3.5 - 2.6 ms | log(M) | 0.247 [-1.063, 1.502] | 0.6520 |
| Expression SDT | Contrast: 3.5 ms; Inverted; 3.5 - 2.6 ms | Type-1 criterion | 0.068 [-0.005, 0.140] | 0.9660 |
| Expression SDT | Contrast: 4.4 ms; Upright; 4.4 - 3.5 ms | d-prime | 0.132 [-0.005, 0.265] | 0.9700 |
| Expression SDT | Contrast: 4.4 ms; Upright; 4.4 - 3.5 ms | meta-d-prime | 0.088 [-0.030, 0.219] | 0.9320 |
| Expression SDT | Contrast: 4.4 ms; Upright; 4.4 - 3.5 ms | M-ratio | -0.135 [-1.008, 0.533] | 0.6530 |
| Expression SDT | Contrast: 4.4 ms; Upright; 4.4 - 3.5 ms | log(M) | -0.157 [-0.961, 0.661] | 0.6530 |
| Expression SDT | Contrast: 4.4 ms; Upright; 4.4 - 3.5 ms | Type-1 criterion | -0.035 [-0.107, 0.036] | 0.8320 |
| Expression SDT | Contrast: 4.4 ms; Inverted; 4.4 - 3.5 ms | d-prime | 0.072 [-0.049, 0.198] | 0.8790 |
| Expression SDT | Contrast: 4.4 ms; Inverted; 4.4 - 3.5 ms | meta-d-prime | 0.048 [-0.042, 0.150] | 0.8610 |
| Expression SDT | Contrast: 4.4 ms; Inverted; 4.4 - 3.5 ms | M-ratio | -0.011 [-0.883, 0.782] | 0.5100 |
| Expression SDT | Contrast: 4.4 ms; Inverted; 4.4 - 3.5 ms | log(M) | -0.016 [-1.148, 1.199] | 0.5100 |

| | | | | |
|---|---|---|---|---|
| Expression SDT | Contrast: 4.4 ms; Inverted; 4.4 - 3.5 ms | Type-1 criterion | -0.045 [-0.118, 0.027] | 0.8880 |
| Expression SDT | Contrast: 5.3 ms; Upright; 5.3 - 4.4 ms | d-prime | 0.115 [-0.028, 0.254] | 0.9420 |
| Expression SDT | Contrast: 5.3 ms; Upright; 5.3 - 4.4 ms | meta-d-prime | 0.169 [0.036, 0.330] | 0.9930 |
| Expression SDT | Contrast: 5.3 ms; Upright; 5.3 - 4.4 ms | M-ratio | 0.198 [-0.347, 0.739] | 0.7830 |
| Expression SDT | Contrast: 5.3 ms; Upright; 5.3 - 4.4 ms | log(M) | 0.225 [-0.346, 0.826] | 0.7830 |
| Expression SDT | Contrast: 5.3 ms; Upright; 5.3 - 4.4 ms | Type-1 criterion | -0.097 [-0.170, -0.024] | 0.9960 |
| Expression SDT | Contrast: 5.3 ms; Inverted; 5.3 - 4.4 ms | d-prime | 0.011 [-0.120, 0.143] | 0.5650 |
| Expression SDT | Contrast: 5.3 ms; Inverted; 5.3 - 4.4 ms | meta-d-prime | 0.039 [-0.069, 0.148] | 0.7670 |
| Expression SDT | Contrast: 5.3 ms; Inverted; 5.3 - 4.4 ms | M-ratio | 0.288 [-0.529, 1.273] | 0.7610 |
| Expression SDT | Contrast: 5.3 ms; Inverted; 5.3 - 4.4 ms | log(M) | 0.354 [-0.623, 1.381] | 0.7610 |
| Expression SDT | Contrast: 5.3 ms; Inverted; 5.3 - 4.4 ms | Type-1 criterion | -0.019 [-0.092, 0.053] | 0.6960 |
| Expression SDT | Contrast: 6.2 ms; Upright; 6.2 - 5.3 ms | d-prime | 0.075 [-0.067, 0.215] | 0.8480 |
| Expression SDT | Contrast: 6.2 ms; Upright; 6.2 - 5.3 ms | meta-d-prime | -0.058 [-0.209, 0.082] | 0.7960 |
| Expression SDT | Contrast: 6.2 ms; Upright; 6.2 - 5.3 ms | M-ratio | -0.288 [-0.792, 0.115] | 0.9250 |
| Expression SDT | Contrast: 6.2 ms; Upright; 6.2 - 5.3 ms | log(M) | -0.343 [-0.845, 0.137] | 0.9250 |
| Expression SDT | Contrast: 6.2 ms; Upright; 6.2 - 5.3 ms | Type-1 criterion | -0.094 [-0.168, -0.022] | 0.9950 |
| Expression SDT | Contrast: 6.2 ms; Inverted; 6.2 - 5.3 ms | d-prime | 0.102 [-0.039, 0.242] | 0.9260 |
| Expression SDT | Contrast: 6.2 ms; Inverted; 6.2 - 5.3 ms | meta-d-prime | 0.078 [-0.035, 0.208] | 0.9120 |
| Expression SDT | Contrast: 6.2 ms; Inverted; 6.2 - 5.3 ms | M-ratio | -0.102 [-1.079, 0.696] | 0.6040 |
| Expression SDT | Contrast: 6.2 ms; Inverted; 6.2 - 5.3 ms | log(M) | -0.112 [-0.977, 0.777] | 0.6040 |
| Expression SDT | Contrast: 6.2 ms; Inverted; 6.2 - 5.3 ms | Type-1 criterion | -0.040 [-0.114, 0.033] | 0.8580 |
| Location psychometric | Estimate: Upright; Neutral | Psychometric midpoint (ms) | 3.297 [3.024, 3.580] | |
| Location psychometric | Estimate: Upright; Neutral | Psychometric scale (ms) | 1.040 [0.912, 1.171] | |
| Location psychometric | Estimate: Upright; Neutral | $ED_{75}$ (ms) | 3.383 [3.108, 3.668] | |
| Location psychometric | Estimate: Upright; Fearful | Psychometric midpoint (ms) | 3.324 [3.051, 3.605] | |
| Location psychometric | Estimate: Upright; Fearful | Psychometric scale (ms) | 1.074 [0.931, 1.234] | |
| Location psychometric | Estimate: Upright; Fearful | $ED_{75}$ (ms) | 3.413 [3.136, 3.695] | |
| Location psychometric | Estimate: Inverted; Neutral | Psychometric midpoint (ms) | 3.760 [3.320, 4.220] | |
| Location psychometric | Estimate: Inverted; Neutral | Psychometric scale (ms) | 1.577 [1.359, 1.841] | |

| | | | | |
|---|---|---|---|---|
| Location psychometric | Estimate: Inverted; Neutral | $ED_{75}$ (ms) | 3.892 [3.444, 4.363] | |
| Location psychometric | Estimate: Inverted; Fearful | Psychometric midpoint (ms) | 3.985 [3.490, 4.489] | |
| Location psychometric | Estimate: Inverted; Fearful | Psychometric scale (ms) | 1.563 [1.378, 1.784] | |
| Location psychometric | Estimate: Inverted; Fearful | $ED_{75}$ (ms) | 4.115 [3.617, 4.624] | |
| Location psychometric | Estimate: Overall | Lapse rate | 0.030 [0.014, 0.054] | |
| Location psychometric | Contrast: neutral: inverted - upright | $ED_{75}$ (ms) | 0.508 [-0.003, 1.048] | 0.9740 |
| Location psychometric | Contrast: fearful: inverted - upright | $ED_{75}$ (ms) | 0.702 [0.141, 1.272] | 0.9930 |
| Location psychometric | Contrast: upright: fearful - neutral | $ED_{75}$ (ms) | 0.032 [-0.368, 0.413] | 0.5670 |
| Location psychometric | Contrast: inverted: fearful - neutral | $ED_{75}$ (ms) | 0.221 [-0.440, 0.878] | 0.7520 |
| Location psychometric | Contrast: inverted - upright | $ED_{75}$ (ms) | 0.606 [0.216, 0.998] | 0.9990 |
| Location psychometric | Contrast: fearful - neutral | $ED_{75}$ (ms) | 0.126 [-0.260, 0.509] | 0.7400 |
| Expression psychometric | Estimate: Upright | Psychometric midpoint (ms) | 8.933 [7.801, 10.963] | |
| Expression psychometric | Estimate: Upright | Psychometric scale (ms) | 2.715 [2.066, 3.940] | |
| Expression psychometric | Estimate: Upright | $ED_{75}$ (ms) | 9.172 [8.038, 11.309] | |
| Expression psychometric | Estimate: Inverted | Psychometric midpoint (ms) | 11.222 [8.758, 16.808] | |
| Expression psychometric | Estimate: Inverted | Psychometric scale (ms) | 2.989 [1.731, 5.504] | |
| Expression psychometric | Estimate: Inverted | $ED_{75}$ (ms) | 11.492 [8.926, 17.279] | |
| Expression psychometric | Estimate: Overall | Lapse rate | 0.026 [0.004, 0.094] | |
| Expression psychometric | Contrast: inverted - upright | $ED_{75}$ (ms) | 2.271 [-0.952, 8.196] | 0.9110 |

**Supplementary Table S6 | Full Experiment 2 behavioural and report-information estimates.** The table combines Bayesian signal-detection estimates, planned contrasts, and information-theoretic results relating PAS to objective reports.

| Sample | Analysis | Condition or contrast | Measure | Result |
|---|---|---|---|---|
| EEG-matched sample (n = 32) | Location SDT | Estimate: 1.7 ms | log(M) | -1.045 [-1.527, -0.524]; Pr = 1.000 |
| EEG-matched sample (n = 32) | Location SDT | Estimate: 1.7 ms | M-ratio | 0.352 [0.217, 0.592]; Pr = 1.000 |
| EEG-matched sample (n = 32) | Location SDT | Estimate: 1.7 ms | d-prime | 0.084 [-0.045, 0.204]; Pr = 0.906 |
| EEG-matched sample (n = 32) | Location SDT | Estimate: 1.7 ms | meta-d-prime | 0.029 [-0.016, 0.083]; Pr = 0.906 |
| EEG-matched sample (n = 32) | Location SDT | Estimate: 1.7 ms | Type-1 criterion | -0.143 [-0.213, -0.073]; Pr = 1.000 |
| EEG-matched sample (n = 32) | Location SDT | Estimate: 4.4 ms | log(M) | -0.625 [-1.045, -0.153]; Pr = 0.994 |

| | | | | |
|---|---|---|---|---|
| EEG-matched sample (n = 32) | Location SDT | Estimate: 4.4 ms | M-ratio | 0.535 [0.352, 0.858]; Pr = 0.994 |
| EEG-matched sample (n = 32) | Location SDT | Estimate: 4.4 ms | d-prime | 1.746 [1.613, 1.867]; Pr = 1.000 |
| EEG-matched sample (n = 32) | Location SDT | Estimate: 4.4 ms | meta-d-prime | 0.934 [0.611, 1.501]; Pr = 1.000 |
| EEG-matched sample (n = 32) | Location SDT | Estimate: 4.4 ms | Type-1 criterion | -0.061 [-0.132, 0.012]; Pr = 0.949 |
| EEG-matched sample (n = 32) | Location SDT | Estimate: 6.2 ms | log(M) | -0.761 [-1.183, -0.283]; Pr = 0.998 |
| EEG-matched sample (n = 32) | Location SDT | Estimate: 6.2 ms | M-ratio | 0.467 [0.306, 0.754]; Pr = 0.998 |
| EEG-matched sample (n = 32) | Location SDT | Estimate: 6.2 ms | d-prime | 2.651 [2.514, 2.779]; Pr = 1.000 |
| EEG-matched sample (n = 32) | Location SDT | Estimate: 6.2 ms | meta-d-prime | 1.238 [0.809, 1.995]; Pr = 1.000 |
| EEG-matched sample (n = 32) | Location SDT | Estimate: 6.2 ms | Type-1 criterion | -0.056 [-0.131, 0.022]; Pr = 0.926 |
| EEG-matched sample (n = 32) | Location SDT | Contrast: 4.4 ms - 1.7 ms | log(M) | 0.415 [0.172, 0.707]; Pr = 1.000 |
| EEG-matched sample (n = 32) | Location SDT | Contrast: 4.4 ms - 1.7 ms | M-ratio | 0.180 [0.075, 0.345]; Pr = 1.000 |
| EEG-matched sample (n = 32) | Location SDT | Contrast: 4.4 ms - 1.7 ms | d-prime | 1.661 [1.618, 1.706]; Pr = 1.000 |
| EEG-matched sample (n = 32) | Location SDT | Contrast: 4.4 ms - 1.7 ms | meta-d-prime | 0.906 [0.594, 1.450]; Pr = 1.000 |
| EEG-matched sample (n = 32) | Location SDT | Contrast: 4.4 ms - 1.7 ms | Type-1 criterion | 0.083 [0.052, 0.114]; Pr = 1.000 |
| EEG-matched sample (n = 32) | Location SDT | Contrast: 6.2 ms - 1.7 ms | log(M) | 0.280 [0.036, 0.572]; Pr = 0.989 |
| EEG-matched sample (n = 32) | Location SDT | Contrast: 6.2 ms - 1.7 ms | M-ratio | 0.112 [0.016, 0.251]; Pr = 0.989 |
| EEG-matched sample (n = 32) | Location SDT | Contrast: 6.2 ms - 1.7 ms | d-prime | 2.567 [2.509, 2.624]; Pr = 1.000 |
| EEG-matched sample (n = 32) | Location SDT | Contrast: 6.2 ms - 1.7 ms | meta-d-prime | 1.209 [0.793, 1.943]; Pr = 1.000 |
| EEG-matched sample (n = 32) | Location SDT | Contrast: 6.2 ms - 1.7 ms | Type-1 criterion | 0.087 [0.048, 0.127]; Pr = 1.000 |
| EEG-matched sample (n = 32) | Location SDT | Contrast: 6.2 ms - 4.4 ms | log(M) | -0.136 [-0.181, -0.090]; Pr = 1.000 |
| EEG-matched sample (n = 32) | Location SDT | Contrast: 6.2 ms - 4.4 ms | M-ratio | -0.067 [-0.117, -0.038]; Pr = 1.000 |
| EEG-matched sample (n = 32) | Location SDT | Contrast: 6.2 ms - 4.4 ms | d-prime | 0.905 [0.843, 0.968]; Pr = 1.000 |
| EEG-matched sample (n = 32) | Location SDT | Contrast: 6.2 ms - 4.4 ms | meta-d-prime | 0.305 [0.194, 0.499]; Pr = 1.000 |
| EEG-matched sample (n = 32) | Location SDT | Contrast: 6.2 ms - 4.4 ms | Type-1 criterion | 0.005 [-0.038, 0.046]; Pr = 0.581 |
| EEG-matched sample (n = 32) | Expression SDT | Estimate: 1.7 ms; Fearful | log(M) | -1.739 [-2.598, -1.029]; Pr = 1.000 |
| EEG-matched sample (n = 32) | Expression SDT | Estimate: 1.7 ms; Fearful | M-ratio | 0.176 [0.074, 0.357]; Pr = 1.000 |
| EEG-matched sample (n = 32) | Expression SDT | Estimate: 1.7 ms; Fearful | d-prime | 0.036 [-0.081, 0.151]; Pr = 0.732 |
| EEG-matched sample (n = 32) | Expression SDT | Estimate: 1.7 ms; Fearful | meta-d-prime | 0.006 [-0.016, 0.033]; Pr = 0.732 |
| EEG-matched sample (n = 32) | Expression SDT | Estimate: 1.7 ms; Fearful | Type-1 criterion | 0.163 [0.044, 0.283]; Pr = 0.996 |

| | | | | |
|---|---|---|---|---|
| EEG-matched sample (n = 32) | Expression SDT | Estimate: 4.4 ms; Fearful | log(M) | -0.007 [-0.466, 0.487]; Pr = 0.513 |
| EEG-matched sample (n = 32) | Expression SDT | Estimate: 4.4 ms; Fearful | M-ratio | 0.993 [0.627, 1.628]; Pr = 0.513 |
| EEG-matched sample (n = 32) | Expression SDT | Estimate: 4.4 ms; Fearful | d-prime | 0.145 [0.032, 0.254]; Pr = 0.993 |
| EEG-matched sample (n = 32) | Expression SDT | Estimate: 4.4 ms; Fearful | meta-d-prime | 0.141 [0.031, 0.299]; Pr = 0.993 |
| EEG-matched sample (n = 32) | Expression SDT | Estimate: 4.4 ms; Fearful | Type-1 criterion | 0.081 [-0.039, 0.200]; Pr = 0.908 |
| EEG-matched sample (n = 32) | Expression SDT | Estimate: 6.2 ms; Fearful | log(M) | -0.088 [-0.526, 0.393]; Pr = 0.653 |
| EEG-matched sample (n = 32) | Expression SDT | Estimate: 6.2 ms; Fearful | M-ratio | 0.915 [0.591, 1.482]; Pr = 0.653 |
| EEG-matched sample (n = 32) | Expression SDT | Estimate: 6.2 ms; Fearful | d-prime | 0.535 [0.418, 0.649]; Pr = 1.000 |
| EEG-matched sample (n = 32) | Expression SDT | Estimate: 6.2 ms; Fearful | meta-d-prime | 0.488 [0.301, 0.809]; Pr = 1.000 |
| EEG-matched sample (n = 32) | Expression SDT | Estimate: 6.2 ms; Fearful | Type-1 criterion | 0.028 [-0.092, 0.148]; Pr = 0.677 |
| EEG-matched sample (n = 32) | Expression SDT | Estimate: 1.7 ms; Happy | log(M) | -1.849 [-2.701, -1.126]; Pr = 1.000 |
| EEG-matched sample (n = 32) | Expression SDT | Estimate: 1.7 ms; Happy | M-ratio | 0.157 [0.067, 0.324]; Pr = 1.000 |
| EEG-matched sample (n = 32) | Expression SDT | Estimate: 1.7 ms; Happy | d-prime | 0.003 [-0.115, 0.117]; Pr = 0.522 |
| EEG-matched sample (n = 32) | Expression SDT | Estimate: 1.7 ms; Happy | meta-d-prime | 0.0004 [-0.021, 0.022]; Pr = 0.522 |
| EEG-matched sample (n = 32) | Expression SDT | Estimate: 1.7 ms; Happy | Type-1 criterion | 0.134 [0.015, 0.253]; Pr = 0.987 |
| EEG-matched sample (n = 32) | Expression SDT | Estimate: 4.4 ms; Happy | log(M) | -0.050 [-0.490, 0.429]; Pr = 0.585 |
| EEG-matched sample (n = 32) | Expression SDT | Estimate: 4.4 ms; Happy | M-ratio | 0.951 [0.613, 1.536]; Pr = 0.585 |
| EEG-matched sample (n = 32) | Expression SDT | Estimate: 4.4 ms; Happy | d-prime | 0.416 [0.300, 0.528]; Pr = 1.000 |
| EEG-matched sample (n = 32) | Expression SDT | Estimate: 4.4 ms; Happy | meta-d-prime | 0.394 [0.232, 0.666]; Pr = 1.000 |
| EEG-matched sample (n = 32) | Expression SDT | Estimate: 4.4 ms; Happy | Type-1 criterion | -0.111 [-0.230, 0.008]; Pr = 0.966 |
| EEG-matched sample (n = 32) | Expression SDT | Estimate: 6.2 ms; Happy | log(M) | -0.084 [-0.507, 0.387]; Pr = 0.647 |
| EEG-matched sample (n = 32) | Expression SDT | Estimate: 6.2 ms; Happy | M-ratio | 0.919 [0.602, 1.472]; Pr = 0.647 |
| EEG-matched sample (n = 32) | Expression SDT | Estimate: 6.2 ms; Happy | d-prime | 1.190 [1.072, 1.306]; Pr = 1.000 |
| EEG-matched sample (n = 32) | Expression SDT | Estimate: 6.2 ms; Happy | meta-d-prime | 1.095 [0.706, 1.768]; Pr = 1.000 |
| EEG-matched sample (n = 32) | Expression SDT | Estimate: 6.2 ms; Happy | Type-1 criterion | -0.214 [-0.333, -0.095]; Pr = 0.999 |
| EEG-matched sample (n = 32) | Expression SDT | Contrast: Fearful; 4.4 ms - 1.7 ms | log(M) | 1.728 [1.101, 2.572]; Pr = 1.000 |
| EEG-matched sample (n = 32) | Expression SDT | Contrast: Fearful; 4.4 ms - 1.7 ms | M-ratio | 0.809 [0.475, 1.400]; Pr = 1.000 |
| EEG-matched sample (n = 32) | Expression SDT | Contrast: Fearful; 4.4 ms - 1.7 ms | d-prime | 0.109 [0.040, 0.178]; Pr = 0.999 |

| | | | | |
|---|---|---|---|---|
| EEG-matched sample (n = 32) | Expression SDT | Contrast: Fearful; 4.4 ms - 1.7 ms | meta-d-prime | 0.135 [0.039, 0.278]; Pr = 0.996 |
| EEG-matched sample (n = 32) | Expression SDT | Contrast: Fearful; 4.4 ms - 1.7 ms | Type-1 criterion | -0.082 [-0.120, -0.044]; Pr = 1.000 |
| EEG-matched sample (n = 32) | Expression SDT | Contrast: Fearful; 6.2 ms - 1.7 ms | log(M) | 1.647 [1.034, 2.494]; Pr = 1.000 |
| EEG-matched sample (n = 32) | Expression SDT | Contrast: Fearful; 6.2 ms - 1.7 ms | M-ratio | 0.732 [0.439, 1.252]; Pr = 1.000 |
| EEG-matched sample (n = 32) | Expression SDT | Contrast: Fearful; 6.2 ms - 1.7 ms | d-prime | 0.499 [0.422, 0.574]; Pr = 1.000 |
| EEG-matched sample (n = 32) | Expression SDT | Contrast: Fearful; 6.2 ms - 1.7 ms | meta-d-prime | 0.482 [0.302, 0.794]; Pr = 1.000 |
| EEG-matched sample (n = 32) | Expression SDT | Contrast: Fearful; 6.2 ms - 1.7 ms | Type-1 criterion | -0.135 [-0.174, -0.097]; Pr = 1.000 |
| EEG-matched sample (n = 32) | Expression SDT | Contrast: Fearful; 6.2 ms - 4.4 ms | log(M) | -0.083 [-0.306, 0.156]; Pr = 0.759 |
| EEG-matched sample (n = 32) | Expression SDT | Contrast: Fearful; 6.2 ms - 4.4 ms | M-ratio | -0.077 [-0.339, 0.149]; Pr = 0.759 |
| EEG-matched sample (n = 32) | Expression SDT | Contrast: Fearful; 6.2 ms - 4.4 ms | d-prime | 0.391 [0.321, 0.460]; Pr = 1.000 |
| EEG-matched sample (n = 32) | Expression SDT | Contrast: Fearful; 6.2 ms - 4.4 ms | meta-d-prime | 0.345 [0.218, 0.572]; Pr = 1.000 |
| EEG-matched sample (n = 32) | Expression SDT | Contrast: Fearful; 6.2 ms - 4.4 ms | Type-1 criterion | -0.053 [-0.091, -0.016]; Pr = 0.998 |
| EEG-matched sample (n = 32) | Expression SDT | Contrast: Happy; 4.4 ms - 1.7 ms | log(M) | 1.795 [1.167, 2.636]; Pr = 1.000 |
| EEG-matched sample (n = 32) | Expression SDT | Contrast: Happy; 4.4 ms - 1.7 ms | M-ratio | 0.786 [0.476, 1.322]; Pr = 1.000 |
| EEG-matched sample (n = 32) | Expression SDT | Contrast: Happy; 4.4 ms - 1.7 ms | d-prime | 0.413 [0.340, 0.486]; Pr = 1.000 |
| EEG-matched sample (n = 32) | Expression SDT | Contrast: Happy; 4.4 ms - 1.7 ms | meta-d-prime | 0.393 [0.238, 0.657]; Pr = 1.000 |
| EEG-matched sample (n = 32) | Expression SDT | Contrast: Happy; 4.4 ms - 1.7 ms | Type-1 criterion | -0.246 [-0.284, -0.207]; Pr = 1.000 |
| EEG-matched sample (n = 32) | Expression SDT | Contrast: Happy; 6.2 ms - 1.7 ms | log(M) | 1.758 [1.139, 2.595]; Pr = 1.000 |
| EEG-matched sample (n = 32) | Expression SDT | Contrast: Happy; 6.2 ms - 1.7 ms | M-ratio | 0.753 [0.467, 1.259]; Pr = 1.000 |
| EEG-matched sample (n = 32) | Expression SDT | Contrast: Happy; 6.2 ms - 1.7 ms | d-prime | 1.188 [1.109, 1.265]; Pr = 1.000 |
| EEG-matched sample (n = 32) | Expression SDT | Contrast: Happy; 6.2 ms - 1.7 ms | meta-d-prime | 1.094 [0.709, 1.763]; Pr = 1.000 |
| EEG-matched sample (n = 32) | Expression SDT | Contrast: Happy; 6.2 ms - 1.7 ms | Type-1 criterion | -0.348 [-0.387, -0.309]; Pr = 1.000 |
| EEG-matched sample (n = 32) | Expression SDT | Contrast: Happy; 6.2 ms - 4.4 ms | log(M) | -0.035 [-0.178, 0.114]; Pr = 0.677 |

| | | | | |
|---|---|---|---|---|
| EEG-matched sample (n = 32) | Expression SDT | Contrast: Happy; 6.2 ms - 4.4 ms | M-ratio | -0.032 [-0.190, 0.108]; Pr = 0.677 |
| EEG-matched sample (n = 32) | Expression SDT | Contrast: Happy; 6.2 ms - 4.4 ms | d-prime | 0.775 [0.701, 0.848]; Pr = 1.000 |
| EEG-matched sample (n = 32) | Expression SDT | Contrast: Happy; 6.2 ms - 4.4 ms | meta-d-prime | 0.699 [0.453, 1.127]; Pr = 1.000 |
| EEG-matched sample (n = 32) | Expression SDT | Contrast: Happy; 6.2 ms - 4.4 ms | Type-1 criterion | -0.102 [-0.141, -0.062]; Pr = 1.000 |
| EEG-matched sample (n = 32) | Expression SDT | Contrast: Happy - fearful; 1.7 ms: happy - fearful | log(M) | -0.106 [-1.119, 0.905]; Pr = 0.586 |
| EEG-matched sample (n = 32) | Expression SDT | Contrast: Happy - fearful; 1.7 ms: happy - fearful | M-ratio | -0.017 [-0.200, 0.154]; Pr = 0.586 |
| EEG-matched sample (n = 32) | Expression SDT | Contrast: Happy - fearful; 1.7 ms: happy - fearful | d-prime | -0.033 [-0.109, 0.043]; Pr = 0.806 |
| EEG-matched sample (n = 32) | Expression SDT | Contrast: Happy - fearful; 1.7 ms: happy - fearful | meta-d-prime | -0.006 [-0.026, 0.011]; Pr = 0.772 |
| EEG-matched sample (n = 32) | Expression SDT | Contrast: Happy - fearful; 1.7 ms: happy - fearful | Type-1 criterion | -0.029 [-0.067, 0.009]; Pr = 0.932 |
| EEG-matched sample (n = 32) | Expression SDT | Contrast: Happy - fearful; 4.4 ms: happy - fearful | log(M) | -0.044 [-0.269, 0.191]; Pr = 0.647 |
| EEG-matched sample (n = 32) | Expression SDT | Contrast: Happy - fearful; 4.4 ms: happy - fearful | M-ratio | -0.042 [-0.294, 0.190]; Pr = 0.647 |
| EEG-matched sample (n = 32) | Expression SDT | Contrast: Happy - fearful; 4.4 ms: happy - fearful | d-prime | 0.271 [0.205, 0.336]; Pr = 1.000 |
| EEG-matched sample (n = 32) | Expression SDT | Contrast: Happy - fearful; 4.4 ms: happy - fearful | meta-d-prime | 0.251 [0.154, 0.416]; Pr = 1.000 |
| EEG-matched sample (n = 32) | Expression SDT | Contrast: Happy - fearful; 4.4 ms: happy - fearful | Type-1 criterion | -0.193 [-0.231, -0.155]; Pr = 1.000 |
| EEG-matched sample (n = 32) | Expression SDT | Contrast: Happy - fearful; 6.2 ms: happy - fearful | log(M) | 0.004 [-0.130, 0.143]; Pr = 0.522 |
| EEG-matched sample (n = 32) | Expression SDT | Contrast: Happy - fearful; 6.2 ms: happy - fearful | M-ratio | 0.004 [-0.133, 0.137]; Pr = 0.522 |
| EEG-matched sample (n = 32) | Expression SDT | Contrast: Happy - fearful; 6.2 ms: happy - fearful | d-prime | 0.655 [0.579, 0.731]; Pr = 1.000 |
| EEG-matched sample (n = 32) | Expression SDT | Contrast: Happy - fearful; 6.2 ms: happy - fearful | meta-d-prime | 0.604 [0.389, 0.977]; Pr = 1.000 |
| EEG-matched sample (n = 32) | Expression SDT | Contrast: Happy - fearful; 6.2 ms: happy - fearful | Type-1 criterion | -0.242 [-0.281, -0.203]; Pr = 1.000 |
| Behaviourally eligible sample (n = 39) | Location SDT | Estimate: 1.7 ms | log(M) | -0.725 [-1.108, -0.312]; Pr = 0.999 |
| Behaviourally eligible sample (n = 39) | Location SDT | Estimate: 1.7 ms | M-ratio | 0.484 [0.330, 0.732]; Pr = 0.999 |

| | | | | |
|---|---|---|---|---|
| Behaviourally eligible sample (n = 39) | Location SDT | Estimate: 1.7 ms | d-prime | 0.107 [-0.009, 0.224]; Pr = 0.964 |
| Behaviourally eligible sample (n = 39) | Location SDT | Estimate: 1.7 ms | meta-d-prime | 0.051 [-0.004, 0.124]; Pr = 0.964 |
| Behaviourally eligible sample (n = 39) | Location SDT | Estimate: 1.7 ms | Type-1 criterion | -0.095 [-0.152, -0.037]; Pr = 0.999 |
| Behaviourally eligible sample (n = 39) | Location SDT | Estimate: 4.4 ms | log(M) | -0.639 [-1.009, -0.245]; Pr = 0.999 |
| Behaviourally eligible sample (n = 39) | Location SDT | Estimate: 4.4 ms | M-ratio | 0.528 [0.365, 0.783]; Pr = 0.999 |
| Behaviourally eligible sample (n = 39) | Location SDT | Estimate: 4.4 ms | d-prime | 1.689 [1.569, 1.809]; Pr = 1.000 |
| Behaviourally eligible sample (n = 39) | Location SDT | Estimate: 4.4 ms | meta-d-prime | 0.891 [0.611, 1.327]; Pr = 1.000 |
| Behaviourally eligible sample (n = 39) | Location SDT | Estimate: 4.4 ms | Type-1 criterion | -0.023 [-0.082, 0.037]; Pr = 0.782 |
| Behaviourally eligible sample (n = 39) | Location SDT | Estimate: 6.2 ms | log(M) | -0.828 [-1.196, -0.431]; Pr = 1.000 |
| Behaviourally eligible sample (n = 39) | Location SDT | Estimate: 6.2 ms | M-ratio | 0.437 [0.302, 0.650]; Pr = 1.000 |
| Behaviourally eligible sample (n = 39) | Location SDT | Estimate: 6.2 ms | d-prime | 2.617 [2.494, 2.742]; Pr = 1.000 |
| Behaviourally eligible sample (n = 39) | Location SDT | Estimate: 6.2 ms | meta-d-prime | 1.143 [0.789, 1.698]; Pr = 1.000 |
| Behaviourally eligible sample (n = 39) | Location SDT | Estimate: 6.2 ms | Type-1 criterion | -0.011 [-0.074, 0.054]; Pr = 0.642 |
| Behaviourally eligible sample (n = 39) | Location SDT | Contrast: 4.4 ms - 1.7 ms | log(M) | 0.084 [-0.042, 0.223]; Pr = 0.898 |
| Behaviourally eligible sample (n = 39) | Location SDT | Contrast: 4.4 ms - 1.7 ms | M-ratio | 0.042 [-0.022, 0.118]; Pr = 0.898 |
| Behaviourally eligible sample (n = 39) | Location SDT | Contrast: 4.4 ms - 1.7 ms | d-prime | 1.581 [1.543, 1.620]; Pr = 1.000 |
| Behaviourally eligible sample (n = 39) | Location SDT | Contrast: 4.4 ms - 1.7 ms | meta-d-prime | 0.839 [0.580, 1.243]; Pr = 1.000 |
| Behaviourally eligible sample (n = 39) | Location SDT | Contrast: 4.4 ms - 1.7 ms | Type-1 criterion | 0.072 [0.045, 0.100]; Pr = 1.000 |
| Behaviourally eligible sample (n = 39) | Location SDT | Contrast: 6.2 ms - 1.7 ms | log(M) | -0.105 [-0.234, 0.034]; Pr = 0.933 |

| | | | | |
|---|---|---|---|---|
| Behaviourally eligible sample (n = 39) | Location SDT | Contrast: 6.2 ms - 1.7 ms | M-ratio | -0.048 [-0.124, 0.015]; Pr = 0.933 |
| Behaviourally eligible sample (n = 39) | Location SDT | Contrast: 6.2 ms - 1.7 ms | d-prime | 2.510 [2.459, 2.561]; Pr = 1.000 |
| Behaviourally eligible sample (n = 39) | Location SDT | Contrast: 6.2 ms - 1.7 ms | meta-d-prime | 1.091 [0.753, 1.621]; Pr = 1.000 |
| Behaviourally eligible sample (n = 39) | Location SDT | Contrast: 6.2 ms - 1.7 ms | Type-1 criterion | 0.084 [0.048, 0.119]; Pr = 1.000 |
| Behaviourally eligible sample (n = 39) | Location SDT | Contrast: 6.2 ms - 4.4 ms | log(M) | -0.189 [-0.229, -0.149]; Pr = 1.000 |
| Behaviourally eligible sample (n = 39) | Location SDT | Contrast: 6.2 ms - 4.4 ms | M-ratio | -0.090 [-0.140, -0.059]; Pr = 1.000 |
| Behaviourally eligible sample (n = 39) | Location SDT | Contrast: 6.2 ms - 4.4 ms | d-prime | 0.928 [0.873, 0.983]; Pr = 1.000 |
| Behaviourally eligible sample (n = 39) | Location SDT | Contrast: 6.2 ms - 4.4 ms | meta-d-prime | 0.251 [0.169, 0.383]; Pr = 1.000 |
| Behaviourally eligible sample (n = 39) | Location SDT | Contrast: 6.2 ms - 4.4 ms | Type-1 criterion | 0.011 [-0.026, 0.050]; Pr = 0.724 |
| Behaviourally eligible sample (n = 39) | Expression SDT | Estimate: 1.7 ms; Fearful | log(M) | -1.986 [-2.814, -1.302]; Pr = 1.000 |
| Behaviourally eligible sample (n = 39) | Expression SDT | Estimate: 1.7 ms; Fearful | M-ratio | 0.137 [0.060, 0.272]; Pr = 1.000 |
| Behaviourally eligible sample (n = 39) | Expression SDT | Estimate: 1.7 ms; Fearful | d-prime | -0.010 [-0.116, 0.096]; Pr = 0.576 |
| Behaviourally eligible sample (n = 39) | Expression SDT | Estimate: 1.7 ms; Fearful | meta-d-prime | -0.001 [-0.019, 0.015]; Pr = 0.576 |
| Behaviourally eligible sample (n = 39) | Expression SDT | Estimate: 1.7 ms; Fearful | Type-1 criterion | 0.197 [0.093, 0.297]; Pr = 1.000 |
| Behaviourally eligible sample (n = 39) | Expression SDT | Estimate: 4.4 ms; Fearful | log(M) | -0.116 [-0.527, 0.342]; Pr = 0.703 |
| Behaviourally eligible sample (n = 39) | Expression SDT | Estimate: 4.4 ms; Fearful | M-ratio | 0.890 [0.590, 1.408]; Pr = 0.703 |
| Behaviourally eligible sample (n = 39) | Expression SDT | Estimate: 4.4 ms; Fearful | d-prime | 0.155 [0.051, 0.258]; Pr = 0.998 |
| Behaviourally eligible sample (n = 39) | Expression SDT | Estimate: 4.4 ms; Fearful | meta-d-prime | 0.137 [0.045, 0.268]; Pr = 0.998 |
| Behaviourally eligible sample (n = 39) | Expression SDT | Estimate: 4.4 ms; Fearful | Type-1 criterion | 0.061 [-0.042, 0.161]; Pr = 0.881 |

| | | | | |
|---|---|---|---|---|
| Behaviourally eligible sample (n = 39) | Expression SDT | Estimate: 6.2 ms; Fearful | log(M) | -0.092 [-0.471, 0.346]; Pr = 0.676 |
| Behaviourally eligible sample (n = 39) | Expression SDT | Estimate: 6.2 ms; Fearful | M-ratio | 0.912 [0.624, 1.413]; Pr = 0.676 |
| Behaviourally eligible sample (n = 39) | Expression SDT | Estimate: 6.2 ms; Fearful | d-prime | 0.543 [0.434, 0.648]; Pr = 1.000 |
| Behaviourally eligible sample (n = 39) | Expression SDT | Estimate: 6.2 ms; Fearful | meta-d-prime | 0.492 [0.325, 0.780]; Pr = 1.000 |
| Behaviourally eligible sample (n = 39) | Expression SDT | Estimate: 6.2 ms; Fearful | Type-1 criterion | -0.025 [-0.129, 0.073]; Pr = 0.696 |
| Behaviourally eligible sample (n = 39) | Expression SDT | Estimate: 1.7 ms; Happy | log(M) | -2.064 [-2.885, -1.369]; Pr = 1.000 |
| Behaviourally eligible sample (n = 39) | Expression SDT | Estimate: 1.7 ms; Happy | M-ratio | 0.127 [0.056, 0.254]; Pr = 1.000 |
| Behaviourally eligible sample (n = 39) | Expression SDT | Estimate: 1.7 ms; Happy | d-prime | -0.014 [-0.121, 0.090]; Pr = 0.608 |
| Behaviourally eligible sample (n = 39) | Expression SDT | Estimate: 1.7 ms; Happy | meta-d-prime | -0.002 [-0.019, 0.013]; Pr = 0.608 |
| Behaviourally eligible sample (n = 39) | Expression SDT | Estimate: 1.7 ms; Happy | Type-1 criterion | 0.158 [0.056, 0.257]; Pr = 0.999 |
| Behaviourally eligible sample (n = 39) | Expression SDT | Estimate: 4.4 ms; Happy | log(M) | 0.013 [-0.364, 0.456]; Pr = 0.526 |
| Behaviourally eligible sample (n = 39) | Expression SDT | Estimate: 4.4 ms; Happy | M-ratio | 1.013 [0.695, 1.577]; Pr = 0.526 |
| Behaviourally eligible sample (n = 39) | Expression SDT | Estimate: 4.4 ms; Happy | d-prime | 0.399 [0.295, 0.503]; Pr = 1.000 |
| Behaviourally eligible sample (n = 39) | Expression SDT | Estimate: 4.4 ms; Happy | meta-d-prime | 0.402 [0.254, 0.655]; Pr = 1.000 |
| Behaviourally eligible sample (n = 39) | Expression SDT | Estimate: 4.4 ms; Happy | Type-1 criterion | -0.125 [-0.227, -0.024]; Pr = 0.991 |
| Behaviourally eligible sample (n = 39) | Expression SDT | Estimate: 6.2 ms; Happy | log(M) | -0.126 [-0.497, 0.308]; Pr = 0.732 |
| Behaviourally eligible sample (n = 39) | Expression SDT | Estimate: 6.2 ms; Happy | M-ratio | 0.882 [0.609, 1.361]; Pr = 0.732 |
| Behaviourally eligible sample (n = 39) | Expression SDT | Estimate: 6.2 ms; Happy | d-prime | 1.151 [1.043, 1.258]; Pr = 1.000 |
| Behaviourally eligible sample (n = 39) | Expression SDT | Estimate: 6.2 ms; Happy | meta-d-prime | 1.014 [0.697, 1.573]; Pr = 1.000 |

| | | | | |
|---|---|---|---|---|
| Behaviourally eligible sample (n = 39) | Expression SDT | Estimate: 6.2 ms; Happy | Type-1 criterion | -0.272 [-0.375, -0.171]; Pr = 1.000 |
| Behaviourally eligible sample (n = 39) | Expression SDT | Contrast: Fearful; 4.4 ms - 1.7 ms | log(M) | 1.870 [1.251, 2.697]; Pr = 1.000 |
| Behaviourally eligible sample (n = 39) | Expression SDT | Contrast: Fearful; 4.4 ms - 1.7 ms | M-ratio | 0.748 [0.472, 1.226]; Pr = 1.000 |
| Behaviourally eligible sample (n = 39) | Expression SDT | Contrast: Fearful; 4.4 ms - 1.7 ms | d-prime | 0.164 [0.100, 0.230]; Pr = 1.000 |
| Behaviourally eligible sample (n = 39) | Expression SDT | Contrast: Fearful; 4.4 ms - 1.7 ms | meta-d-prime | 0.138 [0.056, 0.260]; Pr = 0.999 |
| Behaviourally eligible sample (n = 39) | Expression SDT | Contrast: Fearful; 4.4 ms - 1.7 ms | Type-1 criterion | -0.136 [-0.170, -0.102]; Pr = 1.000 |
| Behaviourally eligible sample (n = 39) | Expression SDT | Contrast: Fearful; 6.2 ms - 1.7 ms | log(M) | 1.893 [1.285, 2.704]; Pr = 1.000 |
| Behaviourally eligible sample (n = 39) | Expression SDT | Contrast: Fearful; 6.2 ms - 1.7 ms | M-ratio | 0.768 [0.501, 1.245]; Pr = 1.000 |
| Behaviourally eligible sample (n = 39) | Expression SDT | Contrast: Fearful; 6.2 ms - 1.7 ms | d-prime | 0.552 [0.484, 0.622]; Pr = 1.000 |
| Behaviourally eligible sample (n = 39) | Expression SDT | Contrast: Fearful; 6.2 ms - 1.7 ms | meta-d-prime | 0.494 [0.332, 0.779]; Pr = 1.000 |
| Behaviourally eligible sample (n = 39) | Expression SDT | Contrast: Fearful; 6.2 ms - 1.7 ms | Type-1 criterion | -0.223 [-0.257, -0.188]; Pr = 1.000 |
| Behaviourally eligible sample (n = 39) | Expression SDT | Contrast: Fearful; 6.2 ms - 4.4 ms | log(M) | 0.023 [-0.174, 0.233]; Pr = 0.583 |
| Behaviourally eligible sample (n = 39) | Expression SDT | Contrast: Fearful; 6.2 ms - 4.4 ms | M-ratio | 0.019 [-0.172, 0.216]; Pr = 0.583 |
| Behaviourally eligible sample (n = 39) | Expression SDT | Contrast: Fearful; 6.2 ms - 4.4 ms | d-prime | 0.388 [0.323, 0.454]; Pr = 1.000 |
| Behaviourally eligible sample (n = 39) | Expression SDT | Contrast: Fearful; 6.2 ms - 4.4 ms | meta-d-prime | 0.356 [0.241, 0.557]; Pr = 1.000 |
| Behaviourally eligible sample (n = 39) | Expression SDT | Contrast: Fearful; 6.2 ms - 4.4 ms | Type-1 criterion | -0.087 [-0.121, -0.053]; Pr = 1.000 |
| Behaviourally eligible sample (n = 39) | Expression SDT | Contrast: Happy; 4.4 ms - 1.7 ms | log(M) | 2.074 [1.454, 2.872]; Pr = 1.000 |
| Behaviourally eligible sample (n = 39) | Expression SDT | Contrast: Happy; 4.4 ms - 1.7 ms | M-ratio | 0.880 [0.579, 1.410]; Pr = 1.000 |
| Behaviourally eligible sample (n = 39) | Expression SDT | Contrast: Happy; 4.4 ms - 1.7 ms | d-prime | 0.414 [0.350, 0.480]; Pr = 1.000 |

| | | | | |
|---|---|---|---|---|
| Behaviourally eligible sample (n = 39) | Expression SDT | Contrast: Happy; 4.4 ms - 1.7 ms | meta-d-prime | 0.404 [0.261, 0.652]; Pr = 1.000 |
| Behaviourally eligible sample (n = 39) | Expression SDT | Contrast: Happy; 4.4 ms - 1.7 ms | Type-1 criterion | -0.282 [-0.317, -0.248]; Pr = 1.000 |
| Behaviourally eligible sample (n = 39) | Expression SDT | Contrast: Happy; 6.2 ms - 1.7 ms | log(M) | 1.935 [1.326, 2.741]; Pr = 1.000 |
| Behaviourally eligible sample (n = 39) | Expression SDT | Contrast: Happy; 6.2 ms - 1.7 ms | M-ratio | 0.750 [0.496, 1.194]; Pr = 1.000 |
| Behaviourally eligible sample (n = 39) | Expression SDT | Contrast: Happy; 6.2 ms - 1.7 ms | d-prime | 1.166 [1.095, 1.237]; Pr = 1.000 |
| Behaviourally eligible sample (n = 39) | Expression SDT | Contrast: Happy; 6.2 ms - 1.7 ms | meta-d-prime | 1.016 [0.700, 1.573]; Pr = 1.000 |
| Behaviourally eligible sample (n = 39) | Expression SDT | Contrast: Happy; 6.2 ms - 1.7 ms | Type-1 criterion | -0.430 [-0.467, -0.394]; Pr = 1.000 |
| Behaviourally eligible sample (n = 39) | Expression SDT | Contrast: Happy; 6.2 ms - 4.4 ms | log(M) | -0.139 [-0.265, -0.013]; Pr = 0.984 |
| Behaviourally eligible sample (n = 39) | Expression SDT | Contrast: Happy; 6.2 ms - 4.4 ms | M-ratio | -0.130 [-0.299, -0.012]; Pr = 0.984 |
| Behaviourally eligible sample (n = 39) | Expression SDT | Contrast: Happy; 6.2 ms - 4.4 ms | d-prime | 0.752 [0.683, 0.819]; Pr = 1.000 |
| Behaviourally eligible sample (n = 39) | Expression SDT | Contrast: Happy; 6.2 ms - 4.4 ms | meta-d-prime | 0.609 [0.415, 0.949]; Pr = 1.000 |
| Behaviourally eligible sample (n = 39) | Expression SDT | Contrast: Happy; 6.2 ms - 4.4 ms | Type-1 criterion | -0.148 [-0.183, -0.113]; Pr = 1.000 |
| Behaviourally eligible sample (n = 39) | Expression SDT | Contrast: Happy - fearful; 1.7 ms: happy - fearful | log(M) | -0.073 [-1.070, 0.931]; Pr = 0.560 |
| Behaviourally eligible sample (n = 39) | Expression SDT | Contrast: Happy - fearful; 1.7 ms: happy - fearful | M-ratio | -0.009 [-0.149, 0.126]; Pr = 0.560 |
| Behaviourally eligible sample (n = 39) | Expression SDT | Contrast: Happy - fearful; 1.7 ms: happy - fearful | d-prime | -0.005 [-0.072, 0.062]; Pr = 0.557 |
| Behaviourally eligible sample (n = 39) | Expression SDT | Contrast: Happy - fearful; 1.7 ms: happy - fearful | meta-d-prime | -0.0006 [-0.013, 0.012]; Pr = 0.541 |
| Behaviourally eligible sample (n = 39) | Expression SDT | Contrast: Happy - fearful; 1.7 ms: happy - fearful | Type-1 criterion | -0.039 [-0.074, -0.005]; Pr = 0.987 |
| Behaviourally eligible sample (n = 39) | Expression SDT | Contrast: Happy - fearful; 4.4 ms: happy - fearful | log(M) | 0.130 [-0.069, 0.341]; Pr = 0.896 |
| Behaviourally eligible sample (n = 39) | Expression SDT | Contrast: Happy - fearful; 4.4 ms: happy - fearful | M-ratio | 0.121 [-0.069, 0.353]; Pr = 0.896 |

| | | | | |
|---|---|---|---|---|
| Behaviourally eligible sample (n = 39) | Expression SDT | Contrast: Happy - fearful; 4.4 ms: happy - fearful | d-prime | 0.245 [0.184, 0.306]; Pr = 1.000 |
| Behaviourally eligible sample (n = 39) | Expression SDT | Contrast: Happy - fearful; 4.4 ms: happy - fearful | meta-d-prime | 0.267 [0.175, 0.423]; Pr = 1.000 |
| Behaviourally eligible sample (n = 39) | Expression SDT | Contrast: Happy - fearful; 4.4 ms: happy - fearful | Type-1 criterion | -0.185 [-0.219, -0.152]; Pr = 1.000 |
| Behaviourally eligible sample (n = 39) | Expression SDT | Contrast: Happy - fearful; 6.2 ms: happy - fearful | log(M) | -0.033 [-0.152, 0.087]; Pr = 0.706 |
| Behaviourally eligible sample (n = 39) | Expression SDT | Contrast: Happy - fearful; 6.2 ms: happy - fearful | M-ratio | -0.029 [-0.149, 0.078]; Pr = 0.706 |
| Behaviourally eligible sample (n = 39) | Expression SDT | Contrast: Happy - fearful; 6.2 ms: happy - fearful | d-prime | 0.609 [0.537, 0.679]; Pr = 1.000 |
| Behaviourally eligible sample (n = 39) | Expression SDT | Contrast: Happy - fearful; 6.2 ms: happy - fearful | meta-d-prime | 0.520 [0.353, 0.816]; Pr = 1.000 |
| Behaviourally eligible sample (n = 39) | Expression SDT | Contrast: Happy - fearful; 6.2 ms: happy - fearful | Type-1 criterion | -0.247 [-0.282, -0.211]; Pr = 1.000 |
| EEG-matched sample (n = 32) | Behavioural report information | Estimate: 1.7 ms | MI(PAS; location accuracy) | 0.0084 [0.0058, 0.0124]; Pr = 1.000 |
| EEG-matched sample (n = 32) | Behavioural report information | Estimate: 1.7 ms | MI(PAS; expression accuracy) | 0.0068 [0.0046, 0.0099]; Pr = 1.000 |
| EEG-matched sample (n = 32) | Behavioural report information | Estimate: 1.7 ms | MI(location accuracy; expression accuracy) | 0.0030 [0.0016, 0.0055]; Pr = 1.000 |
| EEG-matched sample (n = 32) | Behavioural report information | Estimate: 1.7 ms | MI(PAS; joint accuracy) | 0.0212 [0.0170, 0.0265]; Pr = 1.000 |
| EEG-matched sample (n = 32) | Behavioural report information | Estimate: 1.7 ms | CMI(PAS; location accuracy \| expression accuracy) | 0.0143 [0.0110, 0.0188]; Pr = 1.000 |
| EEG-matched sample (n = 32) | Behavioural report information | Estimate: 1.7 ms | CMI(PAS; expression accuracy \| location accuracy) | 0.0127 [0.0098, 0.0166]; Pr = 1.000 |
| EEG-matched sample (n = 32) | Behavioural report information | Estimate: 1.7 ms | Co-information(PAS; location accuracy; expression accuracy) | -0.0058 [-0.0084, -0.0040]; Pr = 1.000 |
| EEG-matched sample (n = 32) | Behavioural report information | Estimate: 1.7 ms | Entropy of location accuracy | 0.9944 [0.9901, 0.9969]; Pr = 1.000 |
| EEG-matched sample (n = 32) | Behavioural report information | Estimate: 1.7 ms | Entropy of expression accuracy | 0.9980 [0.9963, 0.9990]; Pr = 1.000 |

| | | | | |
|---|---|---|---|---|
| EEG-matched sample (n = 32) | Behavioural report information | Estimate: 1.7 ms | Entropy of joint accuracy | 1.9891 [1.9843, 1.9924]; Pr = 1.000 |
| EEG-matched sample (n = 32) | Behavioural report information | Estimate: 1.7 ms | Normalised MI(PAS; location accuracy) | 0.0085 [0.0058, 0.0125]; Pr = 1.000 |
| EEG-matched sample (n = 32) | Behavioural report information | Estimate: 1.7 ms | Normalised MI(PAS; expression accuracy) | 0.0068 [0.0046, 0.0099]; Pr = 1.000 |
| EEG-matched sample (n = 32) | Behavioural report information | Estimate: 1.7 ms | Normalised MI(PAS; joint accuracy) | 0.0107 [0.0086, 0.0133]; Pr = 1.000 |
| EEG-matched sample (n = 32) | Behavioural report information | Estimate: 1.7 ms | CMI(PAS; location report \| stimuli) | 0.0700 [0.0578, 0.0855]; Pr = 1.000 |
| EEG-matched sample (n = 32) | Behavioural report information | Estimate: 1.7 ms | CMI(PAS; expression report \| stimuli) | 0.0669 [0.0568, 0.0799]; Pr = 1.000 |
| EEG-matched sample (n = 32) | Behavioural report information | Estimate: 1.7 ms | CMI(location report; expression report \| stimuli) | 0.0540 [0.0412, 0.0721]; Pr = 1.000 |
| EEG-matched sample (n = 32) | Behavioural report information | Estimate: 1.7 ms | CMI(PAS; joint reports \| stimuli) | 0.1659 [0.1484, 0.1869]; Pr = 1.000 |
| EEG-matched sample (n = 32) | Behavioural report information | Estimate: 1.7 ms | Conditional co-information among reports given stimuli | -0.0288 [-0.0368, -0.0201]; Pr = 1.000 |
| EEG-matched sample (n = 32) | Behavioural report information | Estimate: 4.4 ms | MI(PAS; location accuracy) | 0.0546 [0.0407, 0.0735]; Pr = 1.000 |
| EEG-matched sample (n = 32) | Behavioural report information | Estimate: 4.4 ms | MI(PAS; expression accuracy) | 0.0165 [0.0116, 0.0238]; Pr = 1.000 |
| EEG-matched sample (n = 32) | Behavioural report information | Estimate: 4.4 ms | MI(location accuracy; expression accuracy) | 0.0037 [0.0019, 0.0065]; Pr = 1.000 |
| EEG-matched sample (n = 32) | Behavioural report information | Estimate: 4.4 ms | MI(PAS; joint accuracy) | 0.0762 [0.0612, 0.0954]; Pr = 1.000 |
| EEG-matched sample (n = 32) | Behavioural report information | Estimate: 4.4 ms | CMI(PAS; location accuracy \| expression accuracy) | 0.0594 [0.0455, 0.0781]; Pr = 1.000 |
| EEG-matched sample (n = 32) | Behavioural report information | Estimate: 4.4 ms | CMI(PAS; expression accuracy \| location accuracy) | 0.0213 [0.0161, 0.0286]; Pr = 1.000 |
| EEG-matched sample (n = 32) | Behavioural report information | Estimate: 4.4 ms | Co-information(PAS; location accuracy; expression accuracy) | -0.0047 [-0.0074, -0.0025]; Pr = 1.000 |

| | | | | |
|---|---|---|---|---|
| EEG-matched sample (n = 32) | Behavioural report information | Estimate: 4.4 ms | Entropy of location accuracy | 0.4917 [0.4234, 0.5667]; Pr = 1.000 |
| EEG-matched sample (n = 32) | Behavioural report information | Estimate: 4.4 ms | Entropy of expression accuracy | 0.9798 [0.9618, 0.9893]; Pr = 1.000 |
| EEG-matched sample (n = 32) | Behavioural report information | Estimate: 4.4 ms | Entropy of joint accuracy | 1.4667 [1.3912, 1.5465]; Pr = 1.000 |
| EEG-matched sample (n = 32) | Behavioural report information | Estimate: 4.4 ms | Normalised MI(PAS; location accuracy) | 0.1199 [0.0917, 0.1544]; Pr = 1.000 |
| EEG-matched sample (n = 32) | Behavioural report information | Estimate: 4.4 ms | Normalised MI(PAS; expression accuracy) | 0.0171 [0.0119, 0.0246]; Pr = 1.000 |
| EEG-matched sample (n = 32) | Behavioural report information | Estimate: 4.4 ms | Normalised MI(PAS; joint accuracy) | 0.0529 [0.0425, 0.0655]; Pr = 1.000 |
| EEG-matched sample (n = 32) | Behavioural report information | Estimate: 4.4 ms | CMI(PAS; location report \| stimuli) | 0.0917 [0.0778, 0.1106]; Pr = 1.000 |
| EEG-matched sample (n = 32) | Behavioural report information | Estimate: 4.4 ms | CMI(PAS; expression report \| stimuli) | 0.0775 [0.0677, 0.0893]; Pr = 1.000 |
| EEG-matched sample (n = 32) | Behavioural report information | Estimate: 4.4 ms | CMI(location report; expression report \| stimuli) | 0.0413 [0.0308, 0.0597]; Pr = 1.000 |
| EEG-matched sample (n = 32) | Behavioural report information | Estimate: 4.4 ms | CMI(PAS; joint reports \| stimuli) | 0.1968 [0.1771, 0.2234]; Pr = 1.000 |
| EEG-matched sample (n = 32) | Behavioural report information | Estimate: 4.4 ms | Conditional co-information among reports given stimuli | -0.0276 [-0.0354, -0.0196]; Pr = 1.000 |
| EEG-matched sample (n = 32) | Behavioural report information | Estimate: 6.2 ms | MI(PAS; location accuracy) | 0.0302 [0.0217, 0.0415]; Pr = 1.000 |
| EEG-matched sample (n = 32) | Behavioural report information | Estimate: 6.2 ms | MI(PAS; expression accuracy) | 0.0344 [0.0266, 0.0460]; Pr = 1.000 |
| EEG-matched sample (n = 32) | Behavioural report information | Estimate: 6.2 ms | MI(location accuracy; expression accuracy) | 0.0037 [0.0020, 0.0065]; Pr = 1.000 |
| EEG-matched sample (n = 32) | Behavioural report information | Estimate: 6.2 ms | MI(PAS; joint accuracy) | 0.0685 [0.0572, 0.0833]; Pr = 1.000 |
| EEG-matched sample (n = 32) | Behavioural report information | Estimate: 6.2 ms | CMI(PAS; location accuracy \| expression accuracy) | 0.0338 [0.0252, 0.0453]; Pr = 1.000 |
| EEG-matched sample (n = 32) | Behavioural report information | Estimate: 6.2 ms | CMI(PAS; expression accuracy \| location accuracy) | 0.0380 [0.0298, 0.0498]; Pr = 1.000 |

| | | | | |
|---|---|---|---|---|
| EEG-matched sample (n = 32) | Behavioural report information | Estimate: 6.2 ms | Co-information(PAS; location accuracy; expression accuracy) | -0.0035 [-0.0058, -0.0016]; Pr = 0.999 |
| EEG-matched sample (n = 32) | Behavioural report information | Estimate: 6.2 ms | Entropy of location accuracy | 0.2188 [0.1708, 0.2948]; Pr = 1.000 |
| EEG-matched sample (n = 32) | Behavioural report information | Estimate: 6.2 ms | Entropy of expression accuracy | 0.9021 [0.8687, 0.9286]; Pr = 1.000 |
| EEG-matched sample (n = 32) | Behavioural report information | Estimate: 6.2 ms | Entropy of joint accuracy | 1.1170 [1.0525, 1.2079]; Pr = 1.000 |
| EEG-matched sample (n = 32) | Behavioural report information | Estimate: 6.2 ms | Normalised MI(PAS; location accuracy) | 0.1573 [0.1165, 0.2137]; Pr = 1.000 |
| EEG-matched sample (n = 32) | Behavioural report information | Estimate: 6.2 ms | Normalised MI(PAS; expression accuracy) | 0.0389 [0.0298, 0.0515]; Pr = 1.000 |
| EEG-matched sample (n = 32) | Behavioural report information | Estimate: 6.2 ms | Normalised MI(PAS; joint accuracy) | 0.0620 [0.0518, 0.0747]; Pr = 1.000 |
| EEG-matched sample (n = 32) | Behavioural report information | Estimate: 6.2 ms | CMI(PAS; location report \| stimuli) | 0.0769 [0.0661, 0.0890]; Pr = 1.000 |
| EEG-matched sample (n = 32) | Behavioural report information | Estimate: 6.2 ms | CMI(PAS; expression report \| stimuli) | 0.0875 [0.0756, 0.1023]; Pr = 1.000 |
| EEG-matched sample (n = 32) | Behavioural report information | Estimate: 6.2 ms | CMI(location report; expression report \| stimuli) | 0.0309 [0.0238, 0.0412]; Pr = 1.000 |
| EEG-matched sample (n = 32) | Behavioural report information | Estimate: 6.2 ms | CMI(PAS; joint reports \| stimuli) | 0.1830 [0.1657, 0.2040]; Pr = 1.000 |
| EEG-matched sample (n = 32) | Behavioural report information | Estimate: 6.2 ms | Conditional co-information among reports given stimuli | -0.0185 [-0.0246, -0.0128]; Pr = 1.000 |
| Behaviourally eligible sample (n = 39) | Behavioural report information | Estimate: 1.7 ms | MI(PAS; location accuracy) | 0.0087 [0.0061, 0.0125]; Pr = 1.000 |
| Behaviourally eligible sample (n = 39) | Behavioural report information | Estimate: 1.7 ms | MI(PAS; expression accuracy) | 0.0069 [0.0049, 0.0096]; Pr = 1.000 |
| Behaviourally eligible sample (n = 39) | Behavioural report information | Estimate: 1.7 ms | MI(location accuracy; expression accuracy) | 0.0029 [0.0016, 0.0051]; Pr = 1.000 |
| Behaviourally eligible sample (n = 39) | Behavioural report information | Estimate: 1.7 ms | MI(PAS; joint accuracy) | 0.0215 [0.0173, 0.0265]; Pr = 1.000 |
| Behaviourally eligible sample (n = 39) | Behavioural report information | Estimate: 1.7 ms | CMI(PAS; location accuracy \| | 0.0144 [0.0113, 0.0188]; Pr = 1.000 |

| | | | | |
|---|---|---|---|---|
| | | | expression accuracy) | |
| Behaviourally eligible sample (n = 39) | Behavioural report information | Estimate: 1.7 ms | CMI(PAS; expression accuracy \| location accuracy) | 0.0126 [0.0099, 0.0162]; Pr = 1.000 |
| Behaviourally eligible sample (n = 39) | Behavioural report information | Estimate: 1.7 ms | Co-information(PAS; location accuracy; expression accuracy) | -0.0056 [-0.0080, -0.0040]; Pr = 1.000 |
| Behaviourally eligible sample (n = 39) | Behavioural report information | Estimate: 1.7 ms | Entropy of location accuracy | 0.9916 [0.9837, 0.9954]; Pr = 1.000 |
| Behaviourally eligible sample (n = 39) | Behavioural report information | Estimate: 1.7 ms | Entropy of expression accuracy | 0.9978 [0.9961, 0.9988]; Pr = 1.000 |
| Behaviourally eligible sample (n = 39) | Behavioural report information | Estimate: 1.7 ms | Entropy of joint accuracy | 1.9862 [1.9781, 1.9906]; Pr = 1.000 |
| Behaviourally eligible sample (n = 39) | Behavioural report information | Estimate: 1.7 ms | Normalised MI(PAS; location accuracy) | 0.0089 [0.0062, 0.0127]; Pr = 1.000 |
| Behaviourally eligible sample (n = 39) | Behavioural report information | Estimate: 1.7 ms | Normalised MI(PAS; expression accuracy) | 0.0069 [0.0049, 0.0096]; Pr = 1.000 |
| Behaviourally eligible sample (n = 39) | Behavioural report information | Estimate: 1.7 ms | Normalised MI(PAS; joint accuracy) | 0.0108 [0.0087, 0.0134]; Pr = 1.000 |
| Behaviourally eligible sample (n = 39) | Behavioural report information | Estimate: 1.7 ms | CMI(PAS; location report \| stimuli) | 0.0744 [0.0615, 0.0940]; Pr = 1.000 |
| Behaviourally eligible sample (n = 39) | Behavioural report information | Estimate: 1.7 ms | CMI(PAS; expression report \| stimuli) | 0.0709 [0.0590, 0.0879]; Pr = 1.000 |
| Behaviourally eligible sample (n = 39) | Behavioural report information | Estimate: 1.7 ms | CMI(location report; expression report \| stimuli) | 0.0543 [0.0421, 0.0713]; Pr = 1.000 |
| Behaviourally eligible sample (n = 39) | Behavioural report information | Estimate: 1.7 ms | CMI(PAS; joint reports \| stimuli) | 0.1759 [0.1552, 0.2097]; Pr = 1.000 |
| Behaviourally eligible sample (n = 39) | Behavioural report information | Estimate: 1.7 ms | Conditional co-information among reports given stimuli | -0.0308 [-0.0383, -0.0229]; Pr = 1.000 |
| Behaviourally eligible sample (n = 39) | Behavioural report information | Estimate: 4.4 ms | MI(PAS; location accuracy) | 0.0539 [0.0413, 0.0711]; Pr = 1.000 |
| Behaviourally eligible sample (n = 39) | Behavioural report information | Estimate: 4.4 ms | MI(PAS; expression accuracy) | 0.0175 [0.0124, 0.0245]; Pr = 1.000 |
| Behaviourally eligible sample (n = 39) | Behavioural report information | Estimate: 4.4 ms | MI(location accuracy; expression accuracy) | 0.0035 [0.0020, 0.0061]; Pr = 1.000 |

| | | | | |
|---|---|---|---|---|
| Behaviourally eligible sample (n = 39) | Behavioural report information | Estimate: 4.4 ms | MI(PAS; joint accuracy) | 0.0765 [0.0630, 0.0946]; Pr = 1.000 |
| Behaviourally eligible sample (n = 39) | Behavioural report information | Estimate: 4.4 ms | CMI(PAS; location accuracy \| expression accuracy) | 0.0587 [0.0466, 0.0757]; Pr = 1.000 |
| Behaviourally eligible sample (n = 39) | Behavioural report information | Estimate: 4.4 ms | CMI(PAS; expression accuracy \| location accuracy) | 0.0224 [0.0172, 0.0295]; Pr = 1.000 |
| Behaviourally eligible sample (n = 39) | Behavioural report information | Estimate: 4.4 ms | Co-information(PAS; location accuracy; expression accuracy) | -0.0048 [-0.0072, -0.0028]; Pr = 1.000 |
| Behaviourally eligible sample (n = 39) | Behavioural report information | Estimate: 4.4 ms | Entropy of location accuracy | 0.5071 [0.4353, 0.5757]; Pr = 1.000 |
| Behaviourally eligible sample (n = 39) | Behavioural report information | Estimate: 4.4 ms | Entropy of expression accuracy | 0.9795 [0.9651, 0.9882]; Pr = 1.000 |
| Behaviourally eligible sample (n = 39) | Behavioural report information | Estimate: 4.4 ms | Entropy of joint accuracy | 1.4824 [1.4028, 1.5558]; Pr = 1.000 |
| Behaviourally eligible sample (n = 39) | Behavioural report information | Estimate: 4.4 ms | Normalised MI(PAS; location accuracy) | 0.1170 [0.0915, 0.1484]; Pr = 1.000 |
| Behaviourally eligible sample (n = 39) | Behavioural report information | Estimate: 4.4 ms | Normalised MI(PAS; expression accuracy) | 0.0181 [0.0128, 0.0256]; Pr = 1.000 |
| Behaviourally eligible sample (n = 39) | Behavioural report information | Estimate: 4.4 ms | Normalised MI(PAS; joint accuracy) | 0.0530 [0.0433, 0.0650]; Pr = 1.000 |
| Behaviourally eligible sample (n = 39) | Behavioural report information | Estimate: 4.4 ms | CMI(PAS; location report \| stimuli) | 0.0960 [0.0812, 0.1162]; Pr = 1.000 |
| Behaviourally eligible sample (n = 39) | Behavioural report information | Estimate: 4.4 ms | CMI(PAS; expression report \| stimuli) | 0.0807 [0.0701, 0.0932]; Pr = 1.000 |
| Behaviourally eligible sample (n = 39) | Behavioural report information | Estimate: 4.4 ms | CMI(location report; expression report \| stimuli) | 0.0417 [0.0320, 0.0565]; Pr = 1.000 |
| Behaviourally eligible sample (n = 39) | Behavioural report information | Estimate: 4.4 ms | CMI(PAS; joint reports \| stimuli) | 0.2066 [0.1844, 0.2384]; Pr = 1.000 |
| Behaviourally eligible sample (n = 39) | Behavioural report information | Estimate: 4.4 ms | Conditional co-information among reports given stimuli | -0.0301 [-0.0380, -0.0230]; Pr = 1.000 |
| Behaviourally eligible sample (n = 39) | Behavioural report information | Estimate: 6.2 ms | MI(PAS; location accuracy) | 0.0325 [0.0238, 0.0439]; Pr = 1.000 |

| | | | | |
|---|---|---|---|---|
| Behaviourally eligible sample (n = 39) | Behavioural report information | Estimate: 6.2 ms | MI(PAS; expression accuracy) | 0.0342 [0.0263, 0.0452]; Pr = 1.000 |
| Behaviourally eligible sample (n = 39) | Behavioural report information | Estimate: 6.2 ms | MI(location accuracy; expression accuracy) | 0.0039 [0.0022, 0.0064]; Pr = 1.000 |
| Behaviourally eligible sample (n = 39) | Behavioural report information | Estimate: 6.2 ms | MI(PAS; joint accuracy) | 0.0708 [0.0592, 0.0838]; Pr = 1.000 |
| Behaviourally eligible sample (n = 39) | Behavioural report information | Estimate: 6.2 ms | CMI(PAS; location accuracy \| expression accuracy) | 0.0361 [0.0273, 0.0475]; Pr = 1.000 |
| Behaviourally eligible sample (n = 39) | Behavioural report information | Estimate: 6.2 ms | CMI(PAS; expression accuracy \| location accuracy) | 0.0376 [0.0297, 0.0489]; Pr = 1.000 |
| Behaviourally eligible sample (n = 39) | Behavioural report information | Estimate: 6.2 ms | Co-information(PAS; location accuracy; expression accuracy) | -0.0034 [-0.0056, -0.0016]; Pr = 0.999 |
| Behaviourally eligible sample (n = 39) | Behavioural report information | Estimate: 6.2 ms | Entropy of location accuracy | 0.2314 [0.1844, 0.2980]; Pr = 1.000 |
| Behaviourally eligible sample (n = 39) | Behavioural report information | Estimate: 6.2 ms | Entropy of expression accuracy | 0.9031 [0.8713, 0.9279]; Pr = 1.000 |
| Behaviourally eligible sample (n = 39) | Behavioural report information | Estimate: 6.2 ms | Entropy of joint accuracy | 1.1300 [1.0645, 1.2107]; Pr = 1.000 |
| Behaviourally eligible sample (n = 39) | Behavioural report information | Estimate: 6.2 ms | Normalised MI(PAS; location accuracy) | 0.1557 [0.1189, 0.2044]; Pr = 1.000 |
| Behaviourally eligible sample (n = 39) | Behavioural report information | Estimate: 6.2 ms | Normalised MI(PAS; expression accuracy) | 0.0388 [0.0297, 0.0516]; Pr = 1.000 |
| Behaviourally eligible sample (n = 39) | Behavioural report information | Estimate: 6.2 ms | Normalised MI(PAS; joint accuracy) | 0.0634 [0.0531, 0.0751]; Pr = 1.000 |
| Behaviourally eligible sample (n = 39) | Behavioural report information | Estimate: 6.2 ms | CMI(PAS; location report \| stimuli) | 0.0801 [0.0696, 0.0925]; Pr = 1.000 |
| Behaviourally eligible sample (n = 39) | Behavioural report information | Estimate: 6.2 ms | CMI(PAS; expression report \| stimuli) | 0.0890 [0.0777, 0.1030]; Pr = 1.000 |
| Behaviourally eligible sample (n = 39) | Behavioural report information | Estimate: 6.2 ms | CMI(location report; expression report \| stimuli) | 0.0336 [0.0262, 0.0435]; Pr = 1.000 |
| Behaviourally eligible sample (n = 39) | Behavioural report information | Estimate: 6.2 ms | CMI(PAS; joint reports \| stimuli) | 0.1881 [0.1708, 0.2080]; Pr = 1.000 |

| | | | | |
|---|---|---|---|---|
| Behaviourally eligible sample (n = 39) | Behavioural report information | Estimate: 6.2 ms | Conditional co-information among reports given stimuli | -0.0186 [-0.0247, -0.0127]; Pr = 1.000 |
| EEG-matched sample (n = 32) | Behavioural report information | Contrast: 4.4 - 1.7 ms | MI(PAS; location accuracy) | 0.0461 [0.0320, 0.0651]; Pr = 1.000 |
| EEG-matched sample (n = 32) | Behavioural report information | Contrast: 4.4 - 1.7 ms | MI(PAS; expression accuracy) | 0.0097 [0.0038, 0.0176]; Pr = 0.999 |
| EEG-matched sample (n = 32) | Behavioural report information | Contrast: 4.4 - 1.7 ms | MI(location accuracy; expression accuracy) | 0.0006 [-0.0024, 0.0039]; Pr = 0.667 |
| EEG-matched sample (n = 32) | Behavioural report information | Contrast: 4.4 - 1.7 ms | MI(PAS; joint accuracy) | 0.0549 [0.0392, 0.0748]; Pr = 1.000 |
| EEG-matched sample (n = 32) | Behavioural report information | Contrast: 4.4 - 1.7 ms | CMI(PAS; location accuracy \| expression accuracy) | 0.0450 [0.0307, 0.0637]; Pr = 1.000 |
| EEG-matched sample (n = 32) | Behavioural report information | Contrast: 4.4 - 1.7 ms | CMI(PAS; expression accuracy \| location accuracy) | 0.0086 [0.0020, 0.0166]; Pr = 0.993 |
| EEG-matched sample (n = 32) | Behavioural report information | Contrast: 4.4 - 1.7 ms | Co-information(PAS; location accuracy; expression accuracy) | 0.0011 [-0.0023, 0.0044]; Pr = 0.756 |
| EEG-matched sample (n = 32) | Behavioural report information | Contrast: 4.4 - 1.7 ms | Entropy of location accuracy | -0.5025 [-0.5702, -0.4276]; Pr = 1.000 |
| EEG-matched sample (n = 32) | Behavioural report information | Contrast: 4.4 - 1.7 ms | Entropy of expression accuracy | -0.0180 [-0.0363, -0.0084]; Pr = 1.000 |
| EEG-matched sample (n = 32) | Behavioural report information | Contrast: 4.4 - 1.7 ms | Entropy of joint accuracy | -0.5220 [-0.5969, -0.4423]; Pr = 1.000 |
| EEG-matched sample (n = 32) | Behavioural report information | Contrast: 4.4 - 1.7 ms | Normalised MI(PAS; location accuracy) | 0.1112 [0.0835, 0.1462]; Pr = 1.000 |
| EEG-matched sample (n = 32) | Behavioural report information | Contrast: 4.4 - 1.7 ms | Normalised MI(PAS; expression accuracy) | 0.0103 [0.0041, 0.0183]; Pr = 1.000 |
| EEG-matched sample (n = 32) | Behavioural report information | Contrast: 4.4 - 1.7 ms | Normalised MI(PAS; joint accuracy) | 0.0421 [0.0316, 0.0548]; Pr = 1.000 |
| EEG-matched sample (n = 32) | Behavioural report information | Contrast: 4.4 - 1.7 ms | CMI(PAS; location report \| stimuli) | 0.0221 [0.0073, 0.0369]; Pr = 0.997 |
| EEG-matched sample (n = 32) | Behavioural report information | Contrast: 4.4 - 1.7 ms | CMI(PAS; expression report \| stimuli) | 0.0105 [-0.0024, 0.0229]; Pr = 0.946 |

| | | | | |
|---|---|---|---|---|
| EEG-matched sample (n = 32) | Behavioural report information | Contrast: 4.4 - 1.7 ms | CMI(location report; expression report \| stimuli) | -0.0127 [-0.0315, 0.0094]; Pr = 0.895 |
| EEG-matched sample (n = 32) | Behavioural report information | Contrast: 4.4 - 1.7 ms | CMI(PAS; joint reports \| stimuli) | 0.0314 [0.0098, 0.0527]; Pr = 0.997 |
| EEG-matched sample (n = 32) | Behavioural report information | Contrast: 4.4 - 1.7 ms | Conditional co-information among reports given stimuli | 0.0014 [-0.0099, 0.0110]; Pr = 0.608 |
| EEG-matched sample (n = 32) | Behavioural report information | Contrast: 6.2 - 1.7 ms | MI(PAS; location accuracy) | 0.0217 [0.0121, 0.0336]; Pr = 1.000 |
| EEG-matched sample (n = 32) | Behavioural report information | Contrast: 6.2 - 1.7 ms | MI(PAS; expression accuracy) | 0.0276 [0.0191, 0.0395]; Pr = 1.000 |
| EEG-matched sample (n = 32) | Behavioural report information | Contrast: 6.2 - 1.7 ms | MI(location accuracy; expression accuracy) | 0.0007 [-0.0024, 0.0039]; Pr = 0.673 |
| EEG-matched sample (n = 32) | Behavioural report information | Contrast: 6.2 - 1.7 ms | MI(PAS; joint accuracy) | 0.0473 [0.0348, 0.0628]; Pr = 1.000 |
| EEG-matched sample (n = 32) | Behavioural report information | Contrast: 6.2 - 1.7 ms | CMI(PAS; location accuracy \| expression accuracy) | 0.0193 [0.0095, 0.0315]; Pr = 1.000 |
| EEG-matched sample (n = 32) | Behavioural report information | Contrast: 6.2 - 1.7 ms | CMI(PAS; expression accuracy \| location accuracy) | 0.0252 [0.0163, 0.0374]; Pr = 1.000 |
| EEG-matched sample (n = 32) | Behavioural report information | Contrast: 6.2 - 1.7 ms | Co-information(PAS; location accuracy; expression accuracy) | 0.0023 [-0.0007, 0.0057]; Pr = 0.942 |
| EEG-matched sample (n = 32) | Behavioural report information | Contrast: 6.2 - 1.7 ms | Entropy of location accuracy | -0.7755 [-0.8226, -0.7003]; Pr = 1.000 |
| EEG-matched sample (n = 32) | Behavioural report information | Contrast: 6.2 - 1.7 ms | Entropy of expression accuracy | -0.0959 [-0.1295, -0.0693]; Pr = 1.000 |
| EEG-matched sample (n = 32) | Behavioural report information | Contrast: 6.2 - 1.7 ms | Entropy of joint accuracy | -0.8719 [-0.9364, -0.7811]; Pr = 1.000 |
| EEG-matched sample (n = 32) | Behavioural report information | Contrast: 6.2 - 1.7 ms | Normalised MI(PAS; location accuracy) | 0.1486 [0.1076, 0.2050]; Pr = 1.000 |
| EEG-matched sample (n = 32) | Behavioural report information | Contrast: 6.2 - 1.7 ms | Normalised MI(PAS; expression accuracy) | 0.0321 [0.0224, 0.0449]; Pr = 1.000 |
| EEG-matched sample (n = 32) | Behavioural report information | Contrast: 6.2 - 1.7 ms | Normalised MI(PAS; joint accuracy) | 0.0512 [0.0408, 0.0642]; Pr = 1.000 |

| | | | | |
|---|---|---|---|---|
| EEG-matched sample (n = 32) | Behavioural report information | Contrast: 6.2 - 1.7 ms | CMI(PAS; location report \| stimuli) | 0.0066 [-0.0103, 0.0233]; Pr = 0.789 |
| EEG-matched sample (n = 32) | Behavioural report information | Contrast: 6.2 - 1.7 ms | CMI(PAS; expression report \| stimuli) | 0.0207 [0.0034, 0.0381]; Pr = 0.991 |
| EEG-matched sample (n = 32) | Behavioural report information | Contrast: 6.2 - 1.7 ms | CMI(location report; expression report \| stimuli) | -0.0231 [-0.0415, -0.0063]; Pr = 0.997 |
| EEG-matched sample (n = 32) | Behavioural report information | Contrast: 6.2 - 1.7 ms | CMI(PAS; joint reports \| stimuli) | 0.0170 [-0.0085, 0.0440]; Pr = 0.909 |
| EEG-matched sample (n = 32) | Behavioural report information | Contrast: 6.2 - 1.7 ms | Conditional co-information among reports given stimuli | 0.0103 [0.0004, 0.0198]; Pr = 0.979 |
| EEG-matched sample (n = 32) | Behavioural report information | Contrast: 6.2 - 4.4 ms | MI(PAS; location accuracy) | -0.0242 [-0.0428, -0.0080]; Pr = 0.999 |
| EEG-matched sample (n = 32) | Behavioural report information | Contrast: 6.2 - 4.4 ms | MI(PAS; expression accuracy) | 0.0178 [0.0095, 0.0281]; Pr = 1.000 |
| EEG-matched sample (n = 32) | Behavioural report information | Contrast: 6.2 - 4.4 ms | MI(location accuracy; expression accuracy) | 0.0000 [-0.0034, 0.0032]; Pr = 0.513 |
| EEG-matched sample (n = 32) | Behavioural report information | Contrast: 6.2 - 4.4 ms | MI(PAS; joint accuracy) | -0.0075 [-0.0278, 0.0114]; Pr = 0.784 |
| EEG-matched sample (n = 32) | Behavioural report information | Contrast: 6.2 - 4.4 ms | CMI(PAS; location accuracy \| expression accuracy) | -0.0254 [-0.0444, -0.0092]; Pr = 0.999 |
| EEG-matched sample (n = 32) | Behavioural report information | Contrast: 6.2 - 4.4 ms | CMI(PAS; expression accuracy \| location accuracy) | 0.0166 [0.0080, 0.0274]; Pr = 1.000 |
| EEG-matched sample (n = 32) | Behavioural report information | Contrast: 6.2 - 4.4 ms | Co-information(PAS; location accuracy; expression accuracy) | 0.0012 [-0.0020, 0.0045]; Pr = 0.780 |
| EEG-matched sample (n = 32) | Behavioural report information | Contrast: 6.2 - 4.4 ms | Entropy of location accuracy | -0.2703 [-0.3256, -0.2151]; Pr = 1.000 |
| EEG-matched sample (n = 32) | Behavioural report information | Contrast: 6.2 - 4.4 ms | Entropy of expression accuracy | -0.0772 [-0.1019, -0.0574]; Pr = 1.000 |
| EEG-matched sample (n = 32) | Behavioural report information | Contrast: 6.2 - 4.4 ms | Entropy of joint accuracy | -0.3481 [-0.4041, -0.2918]; Pr = 1.000 |
| EEG-matched sample (n = 32) | Behavioural report information | Contrast: 6.2 - 4.4 ms | Normalised MI(PAS; location accuracy) | 0.0375 [-0.0088, 0.0903]; Pr = 0.950 |

| | | | | |
|---|---|---|---|---|
| EEG-matched sample (n = 32) | Behavioural report information | Contrast: 6.2 - 4.4 ms | Normalised MI(PAS; expression accuracy) | 0.0216 [0.0127, 0.0331]; Pr = 1.000 |
| EEG-matched sample (n = 32) | Behavioural report information | Contrast: 6.2 - 4.4 ms | Normalised MI(PAS; joint accuracy) | 0.0093 [-0.0048, 0.0233]; Pr = 0.900 |
| EEG-matched sample (n = 32) | Behavioural report information | Contrast: 6.2 - 4.4 ms | CMI(PAS; location report \| stimuli) | -0.0151 [-0.0340, 0.0016]; Pr = 0.962 |
| EEG-matched sample (n = 32) | Behavioural report information | Contrast: 6.2 - 4.4 ms | CMI(PAS; expression report \| stimuli) | 0.0102 [-0.0047, 0.0259]; Pr = 0.905 |
| EEG-matched sample (n = 32) | Behavioural report information | Contrast: 6.2 - 4.4 ms | CMI(location report; expression report \| stimuli) | -0.0105 [-0.0244, 0.0005]; Pr = 0.969 |
| EEG-matched sample (n = 32) | Behavioural report information | Contrast: 6.2 - 4.4 ms | CMI(PAS; joint reports \| stimuli) | -0.0140 [-0.0404, 0.0112]; Pr = 0.869 |
| EEG-matched sample (n = 32) | Behavioural report information | Contrast: 6.2 - 4.4 ms | Conditional co-information among reports given stimuli | 0.0089 [0.0005, 0.0177]; Pr = 0.981 |
| Behaviourally eligible sample (n = 39) | Behavioural report information | Contrast: 4.4 - 1.7 ms | MI(PAS; location accuracy) | 0.0450 [0.0322, 0.0626]; Pr = 1.000 |
| Behaviourally eligible sample (n = 39) | Behavioural report information | Contrast: 4.4 - 1.7 ms | MI(PAS; expression accuracy) | 0.0106 [0.0046, 0.0181]; Pr = 1.000 |
| Behaviourally eligible sample (n = 39) | Behavioural report information | Contrast: 4.4 - 1.7 ms | MI(location accuracy; expression accuracy) | 0.0006 [-0.0020, 0.0035]; Pr = 0.671 |
| Behaviourally eligible sample (n = 39) | Behavioural report information | Contrast: 4.4 - 1.7 ms | MI(PAS; joint accuracy) | 0.0551 [0.0402, 0.0735]; Pr = 1.000 |
| Behaviourally eligible sample (n = 39) | Behavioural report information | Contrast: 4.4 - 1.7 ms | CMI(PAS; location accuracy \| expression accuracy) | 0.0443 [0.0316, 0.0611]; Pr = 1.000 |
| Behaviourally eligible sample (n = 39) | Behavioural report information | Contrast: 4.4 - 1.7 ms | CMI(PAS; expression accuracy \| location accuracy) | 0.0097 [0.0031, 0.0177]; Pr = 0.999 |
| Behaviourally eligible sample (n = 39) | Behavioural report information | Contrast: 4.4 - 1.7 ms | Co-information(PAS; location accuracy; expression accuracy) | 0.0008 [-0.0021, 0.0039]; Pr = 0.714 |
| Behaviourally eligible sample (n = 39) | Behavioural report information | Contrast: 4.4 - 1.7 ms | Entropy of location accuracy | -0.4838 [-0.5536, -0.4165]; Pr = 1.000 |
| Behaviourally eligible sample (n = 39) | Behavioural report information | Contrast: 4.4 - 1.7 ms | Entropy of expression accuracy | -0.0182 [-0.0328, -0.0093]; Pr = 1.000 |

| | | | | |
|---|---|---|---|---|
| Behaviourally eligible sample (n = 39) | Behavioural report information | Contrast: 4.4 - 1.7 ms | Entropy of joint accuracy | -0.5033 [-0.5815, -0.4314]; Pr = 1.000 |
| Behaviourally eligible sample (n = 39) | Behavioural report information | Contrast: 4.4 - 1.7 ms | Normalised MI(PAS; location accuracy) | 0.1080 [0.0828, 0.1391]; Pr = 1.000 |
| Behaviourally eligible sample (n = 39) | Behavioural report information | Contrast: 4.4 - 1.7 ms | Normalised MI(PAS; expression accuracy) | 0.0111 [0.0050, 0.0189]; Pr = 1.000 |
| Behaviourally eligible sample (n = 39) | Behavioural report information | Contrast: 4.4 - 1.7 ms | Normalised MI(PAS; joint accuracy) | 0.0420 [0.0321, 0.0541]; Pr = 1.000 |
| Behaviourally eligible sample (n = 39) | Behavioural report information | Contrast: 4.4 - 1.7 ms | CMI(PAS; location report \| stimuli) | 0.0213 [0.0075, 0.0348]; Pr = 0.998 |
| Behaviourally eligible sample (n = 39) | Behavioural report information | Contrast: 4.4 - 1.7 ms | CMI(PAS; expression report \| stimuli) | 0.0096 [-0.0032, 0.0215]; Pr = 0.934 |
| Behaviourally eligible sample (n = 39) | Behavioural report information | Contrast: 4.4 - 1.7 ms | CMI(location report; expression report \| stimuli) | -0.0125 [-0.0296, 0.0052]; Pr = 0.923 |
| Behaviourally eligible sample (n = 39) | Behavioural report information | Contrast: 4.4 - 1.7 ms | CMI(PAS; joint reports \| stimuli) | 0.0304 [0.0106, 0.0502]; Pr = 0.998 |
| Behaviourally eligible sample (n = 39) | Behavioural report information | Contrast: 4.4 - 1.7 ms | Conditional co-information among reports given stimuli | 0.0008 [-0.0097, 0.0097]; Pr = 0.553 |
| Behaviourally eligible sample (n = 39) | Behavioural report information | Contrast: 6.2 - 1.7 ms | MI(PAS; location accuracy) | 0.0238 [0.0141, 0.0357]; Pr = 1.000 |
| Behaviourally eligible sample (n = 39) | Behavioural report information | Contrast: 6.2 - 1.7 ms | MI(PAS; expression accuracy) | 0.0272 [0.0188, 0.0384]; Pr = 1.000 |
| Behaviourally eligible sample (n = 39) | Behavioural report information | Contrast: 6.2 - 1.7 ms | MI(location accuracy; expression accuracy) | 0.0009 [-0.0019, 0.0039]; Pr = 0.748 |
| Behaviourally eligible sample (n = 39) | Behavioural report information | Contrast: 6.2 - 1.7 ms | MI(PAS; joint accuracy) | 0.0492 [0.0362, 0.0629]; Pr = 1.000 |
| Behaviourally eligible sample (n = 39) | Behavioural report information | Contrast: 6.2 - 1.7 ms | CMI(PAS; location accuracy \| expression accuracy) | 0.0216 [0.0116, 0.0337]; Pr = 1.000 |
| Behaviourally eligible sample (n = 39) | Behavioural report information | Contrast: 6.2 - 1.7 ms | CMI(PAS; expression accuracy \| location accuracy) | 0.0250 [0.0162, 0.0366]; Pr = 1.000 |
| Behaviourally eligible sample (n = 39) | Behavioural report information | Contrast: 6.2 - 1.7 ms | Co-information(PAS; location accuracy; expression accuracy) | 0.0022 [-0.0006, 0.0053]; Pr = 0.944 |

| | | | | |
|---|---|---|---|---|
| Behaviourally eligible sample (n = 39) | Behavioural report information | Contrast: 6.2 - 1.7 ms | Entropy of location accuracy | -0.7597 [-0.8063, -0.6932]; Pr = 1.000 |
| Behaviourally eligible sample (n = 39) | Behavioural report information | Contrast: 6.2 - 1.7 ms | Entropy of expression accuracy | -0.0947 [-0.1259, -0.0695]; Pr = 1.000 |
| Behaviourally eligible sample (n = 39) | Behavioural report information | Contrast: 6.2 - 1.7 ms | Entropy of joint accuracy | -0.8561 [-0.9209, -0.7751]; Pr = 1.000 |
| Behaviourally eligible sample (n = 39) | Behavioural report information | Contrast: 6.2 - 1.7 ms | Normalised MI(PAS; location accuracy) | 0.1468 [0.1101, 0.1963]; Pr = 1.000 |
| Behaviourally eligible sample (n = 39) | Behavioural report information | Contrast: 6.2 - 1.7 ms | Normalised MI(PAS; expression accuracy) | 0.0318 [0.0223, 0.0449]; Pr = 1.000 |
| Behaviourally eligible sample (n = 39) | Behavioural report information | Contrast: 6.2 - 1.7 ms | Normalised MI(PAS; joint accuracy) | 0.0525 [0.0417, 0.0644]; Pr = 1.000 |
| Behaviourally eligible sample (n = 39) | Behavioural report information | Contrast: 6.2 - 1.7 ms | CMI(PAS; location report \| stimuli) | 0.0053 [-0.0124, 0.0215]; Pr = 0.736 |
| Behaviourally eligible sample (n = 39) | Behavioural report information | Contrast: 6.2 - 1.7 ms | CMI(PAS; expression report \| stimuli) | 0.0180 [-0.0004, 0.0355]; Pr = 0.973 |
| Behaviourally eligible sample (n = 39) | Behavioural report information | Contrast: 6.2 - 1.7 ms | CMI(location report; expression report \| stimuli) | -0.0205 [-0.0374, -0.0065]; Pr = 0.997 |
| Behaviourally eligible sample (n = 39) | Behavioural report information | Contrast: 6.2 - 1.7 ms | CMI(PAS; joint reports \| stimuli) | 0.0112 [-0.0191, 0.0397]; Pr = 0.785 |
| Behaviourally eligible sample (n = 39) | Behavioural report information | Contrast: 6.2 - 1.7 ms | Conditional co-information among reports given stimuli | 0.0123 [0.0031, 0.0207]; Pr = 0.995 |
| Behaviourally eligible sample (n = 39) | Behavioural report information | Contrast: 6.2 - 4.4 ms | MI(PAS; location accuracy) | -0.0212 [-0.0383, -0.0060]; Pr = 0.997 |
| Behaviourally eligible sample (n = 39) | Behavioural report information | Contrast: 6.2 - 4.4 ms | MI(PAS; expression accuracy) | 0.0165 [0.0090, 0.0261]; Pr = 1.000 |
| Behaviourally eligible sample (n = 39) | Behavioural report information | Contrast: 6.2 - 4.4 ms | MI(location accuracy; expression accuracy) | 0.0003 [-0.0027, 0.0032]; Pr = 0.593 |
| Behaviourally eligible sample (n = 39) | Behavioural report information | Contrast: 6.2 - 4.4 ms | MI(PAS; joint accuracy) | -0.0059 [-0.0257, 0.0114]; Pr = 0.746 |
| Behaviourally eligible sample (n = 39) | Behavioural report information | Contrast: 6.2 - 4.4 ms | CMI(PAS; location accuracy \| expression accuracy) | -0.0226 [-0.0399, -0.0076]; Pr = 0.998 |
| Behaviourally eligible sample (n = 39) | Behavioural report information | Contrast: 6.2 - 4.4 ms | CMI(PAS; expression accuracy \| location accuracy) | 0.0151 [0.0073, 0.0248]; Pr = 1.000 |

| | | | | |
|---|---|---|---|---|
| Behaviourally eligible sample (n = 39) | Behavioural report information | Contrast: 6.2 - 4.4 ms | Co-information(PAS; location accuracy; expression accuracy) | 0.0014 [-0.0015, 0.0043]; Pr = 0.836 |
| Behaviourally eligible sample (n = 39) | Behavioural report information | Contrast: 6.2 - 4.4 ms | Entropy of location accuracy | -0.2737 [-0.3234, -0.2222]; Pr = 1.000 |
| Behaviourally eligible sample (n = 39) | Behavioural report information | Contrast: 6.2 - 4.4 ms | Entropy of expression accuracy | -0.0761 [-0.0989, -0.0568]; Pr = 1.000 |
| Behaviourally eligible sample (n = 39) | Behavioural report information | Contrast: 6.2 - 4.4 ms | Entropy of joint accuracy | -0.3503 [-0.4001, -0.3000]; Pr = 1.000 |
| Behaviourally eligible sample (n = 39) | Behavioural report information | Contrast: 6.2 - 4.4 ms | Normalised MI(PAS; location accuracy) | 0.0395 [-0.0023, 0.0847]; Pr = 0.965 |
| Behaviourally eligible sample (n = 39) | Behavioural report information | Contrast: 6.2 - 4.4 ms | Normalised MI(PAS; expression accuracy) | 0.0205 [0.0121, 0.0308]; Pr = 1.000 |
| Behaviourally eligible sample (n = 39) | Behavioural report information | Contrast: 6.2 - 4.4 ms | Normalised MI(PAS; joint accuracy) | 0.0102 [-0.0032, 0.0230]; Pr = 0.939 |
| Behaviourally eligible sample (n = 39) | Behavioural report information | Contrast: 6.2 - 4.4 ms | CMI(PAS; location report \| stimuli) | -0.0160 [-0.0342, 0.0006]; Pr = 0.970 |
| Behaviourally eligible sample (n = 39) | Behavioural report information | Contrast: 6.2 - 4.4 ms | CMI(PAS; expression report \| stimuli) | 0.0084 [-0.0061, 0.0228]; Pr = 0.874 |
| Behaviourally eligible sample (n = 39) | Behavioural report information | Contrast: 6.2 - 4.4 ms | CMI(location report; expression report \| stimuli) | -0.0082 [-0.0200, 0.0019]; Pr = 0.947 |
| Behaviourally eligible sample (n = 39) | Behavioural report information | Contrast: 6.2 - 4.4 ms | CMI(PAS; joint reports \| stimuli) | -0.0189 [-0.0461, 0.0057]; Pr = 0.937 |
| Behaviourally eligible sample (n = 39) | Behavioural report information | Contrast: 6.2 - 4.4 ms | Conditional co-information among reports given stimuli | 0.0116 [0.0037, 0.0201]; Pr = 0.997 |
| EEG-matched sample (n = 32) | Behavioural report information | Permutation test: 1.7 ms | MI(PAS; location accuracy) | Observed = 0.0049 bits; Excess over null = 0.0019 bits; Holm p = 0.0015 |
| EEG-matched sample (n = 32) | Behavioural report information | Permutation test: 1.7 ms | MI(PAS; expression accuracy) | Observed = 0.0033 bits; Excess over null = 0.0003 bits; Holm p = 0.2344 |
| EEG-matched sample (n = 32) | Behavioural report information | Permutation test: 1.7 ms | MI(location accuracy; expression accuracy) | Observed = 0.0019 bits; Excess over null = 0.0006 bits; Holm p = 0.0340 |
| EEG-matched sample (n = 32) | Behavioural report information | Permutation test: 1.7 ms | MI(PAS; joint accuracy) | Observed = 0.0110 bits; Excess over null = 0.0020 bits; Holm p = 0.0065 |
| EEG-matched sample (n = 32) | Behavioural report information | Permutation test: 1.7 ms | CMI(PAS; location accuracy \| | Observed = 0.0077 bits; Excess over null = 0.0017 bits; Holm p = 0.0035 |

| | | | | |
|---|---|---|---|---|
| | | | expression accuracy) | |
| EEG-matched sample (n = 32) | Behavioural report information | Permutation test: 1.7 ms | CMI(PAS; expression accuracy \| location accuracy) | Observed = 0.0061 bits; Excess over null = 0.0001 bits; Holm p = 0.4538 |
| EEG-matched sample (n = 32) | Behavioural report information | Permutation test: 1.7 ms | Co-information(PAS; location accuracy; expression accuracy) | Observed = -0.0028 bits; Excess over null = 0.0002 bits; Holm p = 0.6247 |
| EEG-matched sample (n = 32) | Behavioural report information | Permutation test: 1.7 ms | Entropy of location accuracy | Observed = 0.9954 bits; Excess over null = 0.0000 bits |
| EEG-matched sample (n = 32) | Behavioural report information | Permutation test: 1.7 ms | Entropy of expression accuracy | Observed = 0.9991 bits; Excess over null = -0.0000 bits |
| EEG-matched sample (n = 32) | Behavioural report information | Permutation test: 1.7 ms | Entropy of joint accuracy | Observed = 1.9927 bits; Excess over null = 0.0000 bits |
| EEG-matched sample (n = 32) | Behavioural report information | Permutation test: 1.7 ms | Normalised MI(PAS; location accuracy) | Observed = 0.0050 bits; Excess over null = 0.0020 bits; Holm p = 0.0015 |
| EEG-matched sample (n = 32) | Behavioural report information | Permutation test: 1.7 ms | Normalised MI(PAS; expression accuracy) | Observed = 0.0033 bits; Excess over null = 0.0003 bits; Holm p = 0.2344 |
| EEG-matched sample (n = 32) | Behavioural report information | Permutation test: 1.7 ms | Normalised MI(PAS; joint accuracy) | Observed = 0.0055 bits; Excess over null = 0.0010 bits; Holm p = 0.0065 |
| EEG-matched sample (n = 32) | Behavioural report information | Permutation test: 1.7 ms | CMI(PAS; location report \| stimuli) | Observed = 0.0465 bits; Excess over null = 0.0236 bits; Holm p = 0.0015 |
| EEG-matched sample (n = 32) | Behavioural report information | Permutation test: 1.7 ms | CMI(PAS; expression report \| stimuli) | Observed = 0.0435 bits; Excess over null = 0.0176 bits; Holm p = 0.0015 |
| EEG-matched sample (n = 32) | Behavioural report information | Permutation test: 1.7 ms | CMI(location report; expression report \| stimuli) | Observed = 0.0456 bits; Excess over null = 0.0348 bits; Holm p = 0.0015 |
| EEG-matched sample (n = 32) | Behavioural report information | Permutation test: 1.7 ms | CMI(PAS; joint reports \| stimuli) | Observed = 0.1031 bits; Excess over null = 0.0449 bits; Holm p = 0.0015 |
| EEG-matched sample (n = 32) | Behavioural report information | Permutation test: 1.7 ms | Conditional co-information among reports given stimuli | Observed = -0.0131 bits; Excess over null = -0.0037 bits; Holm p = 0.0015 |
| EEG-matched sample (n = 32) | Behavioural report information | Permutation test: 4.4 ms | MI(PAS; location accuracy) | Observed = 0.0517 bits; Excess over null = 0.0479 bits; Holm p = 0.0015 |
| EEG-matched sample (n = 32) | Behavioural report information | Permutation test: 4.4 ms | MI(PAS; expression accuracy) | Observed = 0.0130 bits; Excess over null = 0.0095 bits; Holm p = 0.0015 |
| EEG-matched sample (n = 32) | Behavioural report information | Permutation test: 4.4 ms | MI(location accuracy; expression accuracy) | Observed = 0.0026 bits; Excess over null = 0.0014 bits; Holm p = 0.0015 |

| | | | | |
|---|---|---|---|---|
| EEG-matched sample (n = 32) | Behavioural report information | Permutation test: 4.4 ms | MI(PAS; joint accuracy) | Observed = 0.0663 bits; Excess over null = 0.0560 bits; Holm p = 0.0015 |
| EEG-matched sample (n = 32) | Behavioural report information | Permutation test: 4.4 ms | CMI(PAS; location accuracy \| expression accuracy) | Observed = 0.0533 bits; Excess over null = 0.0465 bits; Holm p = 0.0015 |
| EEG-matched sample (n = 32) | Behavioural report information | Permutation test: 4.4 ms | CMI(PAS; expression accuracy \| location accuracy) | Observed = 0.0147 bits; Excess over null = 0.0085 bits; Holm p = 0.0015 |
| EEG-matched sample (n = 32) | Behavioural report information | Permutation test: 4.4 ms | Co-information(PAS; location accuracy; expression accuracy) | Observed = -0.0017 bits; Excess over null = 0.0014 bits; Holm p = 0.0110 |
| EEG-matched sample (n = 32) | Behavioural report information | Permutation test: 4.4 ms | Entropy of location accuracy | Observed = 0.4940 bits; Excess over null = -0.0000 bits |
| EEG-matched sample (n = 32) | Behavioural report information | Permutation test: 4.4 ms | Entropy of expression accuracy | Observed = 0.9802 bits; Excess over null = 0.0000 bits |
| EEG-matched sample (n = 32) | Behavioural report information | Permutation test: 4.4 ms | Entropy of joint accuracy | Observed = 1.4716 bits; Excess over null = 0.0000 bits |
| EEG-matched sample (n = 32) | Behavioural report information | Permutation test: 4.4 ms | Normalised MI(PAS; location accuracy) | Observed = 0.1120 bits; Excess over null = 0.1028 bits; Holm p = 0.0015 |
| EEG-matched sample (n = 32) | Behavioural report information | Permutation test: 4.4 ms | Normalised MI(PAS; expression accuracy) | Observed = 0.0135 bits; Excess over null = 0.0099 bits; Holm p = 0.0015 |
| EEG-matched sample (n = 32) | Behavioural report information | Permutation test: 4.4 ms | Normalised MI(PAS; joint accuracy) | Observed = 0.0458 bits; Excess over null = 0.0387 bits; Holm p = 0.0015 |
| EEG-matched sample (n = 32) | Behavioural report information | Permutation test: 4.4 ms | CMI(PAS; location report \| stimuli) | Observed = 0.0684 bits; Excess over null = 0.0430 bits; Holm p = 0.0015 |
| EEG-matched sample (n = 32) | Behavioural report information | Permutation test: 4.4 ms | CMI(PAS; expression report \| stimuli) | Observed = 0.0524 bits; Excess over null = 0.0300 bits; Holm p = 0.0015 |
| EEG-matched sample (n = 32) | Behavioural report information | Permutation test: 4.4 ms | CMI(location report; expression report \| stimuli) | Observed = 0.0336 bits; Excess over null = 0.0243 bits; Holm p = 0.0015 |
| EEG-matched sample (n = 32) | Behavioural report information | Permutation test: 4.4 ms | CMI(PAS; joint reports \| stimuli) | Observed = 0.1327 bits; Excess over null = 0.0742 bits; Holm p = 0.0015 |
| EEG-matched sample (n = 32) | Behavioural report information | Permutation test: 4.4 ms | Conditional co-information among reports given stimuli | Observed = -0.0118 bits; Excess over null = -0.0013 bits; Holm p = 0.1409 |
| EEG-matched sample (n = 32) | Behavioural report information | Permutation test: 6.2 ms | MI(PAS; location accuracy) | Observed = 0.0271 bits; Excess over null = 0.0221 bits; Holm p = 0.0015 |

| | | | | |
|---|---|---|---|---|
| EEG-matched sample (n = 32) | Behavioural report information | Permutation test: 6.2 ms | MI(PAS; expression accuracy) | Observed = 0.0311 bits; Excess over null = 0.0275 bits; Holm p = 0.0015 |
| EEG-matched sample (n = 32) | Behavioural report information | Permutation test: 6.2 ms | MI(location accuracy; expression accuracy) | Observed = 0.0026 bits; Excess over null = 0.0015 bits; Holm p = 0.0015 |
| EEG-matched sample (n = 32) | Behavioural report information | Permutation test: 6.2 ms | MI(PAS; joint accuracy) | Observed = 0.0591 bits; Excess over null = 0.0486 bits; Holm p = 0.0015 |
| EEG-matched sample (n = 32) | Behavioural report information | Permutation test: 6.2 ms | CMI(PAS; location accuracy \| expression accuracy) | Observed = 0.0280 bits; Excess over null = 0.0209 bits; Holm p = 0.0015 |
| EEG-matched sample (n = 32) | Behavioural report information | Permutation test: 6.2 ms | CMI(PAS; expression accuracy \| location accuracy) | Observed = 0.0320 bits; Excess over null = 0.0264 bits; Holm p = 0.0015 |
| EEG-matched sample (n = 32) | Behavioural report information | Permutation test: 6.2 ms | Co-information(PAS; location accuracy; expression accuracy) | Observed = -0.0009 bits; Excess over null = 0.0011 bits; Holm p = 0.0075 |
| EEG-matched sample (n = 32) | Behavioural report information | Permutation test: 6.2 ms | Entropy of location accuracy | Observed = 0.2240 bits; Excess over null = 0.0000 bits |
| EEG-matched sample (n = 32) | Behavioural report information | Permutation test: 6.2 ms | Entropy of expression accuracy | Observed = 0.9025 bits; Excess over null = 0.0000 bits |
| EEG-matched sample (n = 32) | Behavioural report information | Permutation test: 6.2 ms | Entropy of joint accuracy | Observed = 1.1239 bits; Excess over null = -0.0000 bits |
| EEG-matched sample (n = 32) | Behavioural report information | Permutation test: 6.2 ms | Normalised MI(PAS; location accuracy) | Observed = 0.1368 bits; Excess over null = 0.0998 bits; Holm p = 0.0015 |
| EEG-matched sample (n = 32) | Behavioural report information | Permutation test: 6.2 ms | Normalised MI(PAS; expression accuracy) | Observed = 0.0350 bits; Excess over null = 0.0310 bits; Holm p = 0.0015 |
| EEG-matched sample (n = 32) | Behavioural report information | Permutation test: 6.2 ms | Normalised MI(PAS; joint accuracy) | Observed = 0.0532 bits; Excess over null = 0.0436 bits; Holm p = 0.0015 |
| EEG-matched sample (n = 32) | Behavioural report information | Permutation test: 6.2 ms | CMI(PAS; location report \| stimuli) | Observed = 0.0542 bits; Excess over null = 0.0193 bits; Holm p = 0.0015 |
| EEG-matched sample (n = 32) | Behavioural report information | Permutation test: 6.2 ms | CMI(PAS; expression report \| stimuli) | Observed = 0.0631 bits; Excess over null = 0.0401 bits; Holm p = 0.0015 |
| EEG-matched sample (n = 32) | Behavioural report information | Permutation test: 6.2 ms | CMI(location report; expression report \| stimuli) | Observed = 0.0230 bits; Excess over null = 0.0110 bits; Holm p = 0.0015 |
| EEG-matched sample (n = 32) | Behavioural report information | Permutation test: 6.2 ms | CMI(PAS; joint reports \| stimuli) | Observed = 0.1219 bits; Excess over null = 0.0602 bits; Holm p = 0.0015 |

| | | | | |
|---|---|---|---|---|
| EEG-matched sample (n = 32) | Behavioural report information | Permutation test: 6.2 ms | Conditional co-information among reports given stimuli | Observed = -0.0046 bits; Excess over null = -0.0009 bits; Holm p = 0.1409 |
| Behaviourally eligible sample (n = 39) | Behavioural report information | Permutation test: 1.7 ms | MI(PAS; location accuracy) | Observed = 0.0053 bits; Excess over null = 0.0022 bits; Holm p = 0.0015 |
| Behaviourally eligible sample (n = 39) | Behavioural report information | Permutation test: 1.7 ms | MI(PAS; expression accuracy) | Observed = 0.0034 bits; Excess over null = 0.0003 bits; Holm p = 0.2234 |
| Behaviourally eligible sample (n = 39) | Behavioural report information | Permutation test: 1.7 ms | MI(location accuracy; expression accuracy) | Observed = 0.0017 bits; Excess over null = 0.0005 bits; Holm p = 0.0665 |
| Behaviourally eligible sample (n = 39) | Behavioural report information | Permutation test: 1.7 ms | MI(PAS; joint accuracy) | Observed = 0.0112 bits; Excess over null = 0.0021 bits; Holm p = 0.0030 |
| Behaviourally eligible sample (n = 39) | Behavioural report information | Permutation test: 1.7 ms | CMI(PAS; location accuracy \| expression accuracy) | Observed = 0.0079 bits; Excess over null = 0.0017 bits; Holm p = 0.0045 |
| Behaviourally eligible sample (n = 39) | Behavioural report information | Permutation test: 1.7 ms | CMI(PAS; expression accuracy \| location accuracy) | Observed = 0.0060 bits; Excess over null = -0.0001 bits; Holm p = 0.5647 |
| Behaviourally eligible sample (n = 39) | Behavioural report information | Permutation test: 1.7 ms | Co-information(PAS; location accuracy; expression accuracy) | Observed = -0.0026 bits; Excess over null = 0.0004 bits; Holm p = 0.3039 |
| Behaviourally eligible sample (n = 39) | Behavioural report information | Permutation test: 1.7 ms | Entropy of location accuracy | Observed = 0.9923 bits; Excess over null = 0.0000 bits |
| Behaviourally eligible sample (n = 39) | Behavioural report information | Permutation test: 1.7 ms | Entropy of expression accuracy | Observed = 0.9990 bits; Excess over null = 0.0000 bits |
| Behaviourally eligible sample (n = 39) | Behavioural report information | Permutation test: 1.7 ms | Entropy of joint accuracy | Observed = 1.9895 bits; Excess over null = 0.0000 bits |
| Behaviourally eligible sample (n = 39) | Behavioural report information | Permutation test: 1.7 ms | Normalised MI(PAS; location accuracy) | Observed = 0.0054 bits; Excess over null = 0.0023 bits; Holm p = 0.0015 |
| Behaviourally eligible sample (n = 39) | Behavioural report information | Permutation test: 1.7 ms | Normalised MI(PAS; expression accuracy) | Observed = 0.0034 bits; Excess over null = 0.0003 bits; Holm p = 0.2234 |
| Behaviourally eligible sample (n = 39) | Behavioural report information | Permutation test: 1.7 ms | Normalised MI(PAS; joint accuracy) | Observed = 0.0057 bits; Excess over null = 0.0011 bits; Holm p = 0.0030 |
| Behaviourally eligible sample (n = 39) | Behavioural report information | Permutation test: 1.7 ms | CMI(PAS; location report \| stimuli) | Observed = 0.0514 bits; Excess over null = 0.0291 bits; Holm p = 0.0015 |
| Behaviourally eligible sample (n = 39) | Behavioural report information | Permutation test: 1.7 ms | CMI(PAS; expression report \| stimuli) | Observed = 0.0476 bits; Excess over null = 0.0229 bits; Holm p = 0.0015 |

| | | | | |
|---|---|---|---|---|
| Behaviourally eligible sample (n = 39) | Behavioural report information | Permutation test: 1.7 ms | CMI(location report; expression report \| stimuli) | Observed = 0.0461 bits; Excess over null = 0.0356 bits; Holm p = 0.0015 |
| Behaviourally eligible sample (n = 39) | Behavioural report information | Permutation test: 1.7 ms | CMI(PAS; joint reports \| stimuli) | Observed = 0.1140 bits; Excess over null = 0.0569 bits; Holm p = 0.0015 |
| Behaviourally eligible sample (n = 39) | Behavioural report information | Permutation test: 1.7 ms | Conditional co-information among reports given stimuli | Observed = -0.0150 bits; Excess over null = -0.0050 bits; Holm p = 0.0015 |
| Behaviourally eligible sample (n = 39) | Behavioural report information | Permutation test: 4.4 ms | MI(PAS; location accuracy) | Observed = 0.0508 bits; Excess over null = 0.0470 bits; Holm p = 0.0015 |
| Behaviourally eligible sample (n = 39) | Behavioural report information | Permutation test: 4.4 ms | MI(PAS; expression accuracy) | Observed = 0.0141 bits; Excess over null = 0.0105 bits; Holm p = 0.0015 |
| Behaviourally eligible sample (n = 39) | Behavioural report information | Permutation test: 4.4 ms | MI(location accuracy; expression accuracy) | Observed = 0.0024 bits; Excess over null = 0.0012 bits; Holm p = 0.0015 |
| Behaviourally eligible sample (n = 39) | Behavioural report information | Permutation test: 4.4 ms | MI(PAS; joint accuracy) | Observed = 0.0666 bits; Excess over null = 0.0562 bits; Holm p = 0.0015 |
| Behaviourally eligible sample (n = 39) | Behavioural report information | Permutation test: 4.4 ms | CMI(PAS; location accuracy \| expression accuracy) | Observed = 0.0525 bits; Excess over null = 0.0457 bits; Holm p = 0.0015 |
| Behaviourally eligible sample (n = 39) | Behavioural report information | Permutation test: 4.4 ms | CMI(PAS; expression accuracy \| location accuracy) | Observed = 0.0159 bits; Excess over null = 0.0096 bits; Holm p = 0.0015 |
| Behaviourally eligible sample (n = 39) | Behavioural report information | Permutation test: 4.4 ms | Co-information(PAS; location accuracy; expression accuracy) | Observed = -0.0018 bits; Excess over null = 0.0013 bits; Holm p = 0.0090 |
| Behaviourally eligible sample (n = 39) | Behavioural report information | Permutation test: 4.4 ms | Entropy of location accuracy | Observed = 0.5082 bits; Excess over null = 0.0000 bits |
| Behaviourally eligible sample (n = 39) | Behavioural report information | Permutation test: 4.4 ms | Entropy of expression accuracy | Observed = 0.9800 bits; Excess over null = 0.0000 bits |
| Behaviourally eligible sample (n = 39) | Behavioural report information | Permutation test: 4.4 ms | Entropy of joint accuracy | Observed = 1.4858 bits; Excess over null = -0.0000 bits |
| Behaviourally eligible sample (n = 39) | Behavioural report information | Permutation test: 4.4 ms | Normalised MI(PAS; location accuracy) | Observed = 0.1084 bits; Excess over null = 0.0991 bits; Holm p = 0.0015 |
| Behaviourally eligible sample (n = 39) | Behavioural report information | Permutation test: 4.4 ms | Normalised MI(PAS; expression accuracy) | Observed = 0.0146 bits; Excess over null = 0.0110 bits; Holm p = 0.0015 |
| Behaviourally eligible sample (n = 39) | Behavioural report information | Permutation test: 4.4 ms | Normalised MI(PAS; joint accuracy) | Observed = 0.0459 bits; Excess over null = 0.0388 bits; Holm p = 0.0015 |

| | | | | |
|---|---|---|---|---|
| Behaviourally eligible sample (n = 39) | Behavioural report information | Permutation test: 4.4 ms | CMI(PAS; location report \| stimuli) | Observed = 0.0727 bits; Excess over null = 0.0475 bits; Holm p = 0.0015 |
| Behaviourally eligible sample (n = 39) | Behavioural report information | Permutation test: 4.4 ms | CMI(PAS; expression report \| stimuli) | Observed = 0.0559 bits; Excess over null = 0.0335 bits; Holm p = 0.0015 |
| Behaviourally eligible sample (n = 39) | Behavioural report information | Permutation test: 4.4 ms | CMI(location report; expression report \| stimuli) | Observed = 0.0337 bits; Excess over null = 0.0248 bits; Holm p = 0.0015 |
| Behaviourally eligible sample (n = 39) | Behavioural report information | Permutation test: 4.4 ms | CMI(PAS; joint reports \| stimuli) | Observed = 0.1432 bits; Excess over null = 0.0847 bits; Holm p = 0.0015 |
| Behaviourally eligible sample (n = 39) | Behavioural report information | Permutation test: 4.4 ms | Conditional co-information among reports given stimuli | Observed = -0.0147 bits; Excess over null = -0.0037 bits; Holm p = 0.0015 |
| Behaviourally eligible sample (n = 39) | Behavioural report information | Permutation test: 6.2 ms | MI(PAS; location accuracy) | Observed = 0.0296 bits; Excess over null = 0.0248 bits; Holm p = 0.0015 |
| Behaviourally eligible sample (n = 39) | Behavioural report information | Permutation test: 6.2 ms | MI(PAS; expression accuracy) | Observed = 0.0308 bits; Excess over null = 0.0272 bits; Holm p = 0.0015 |
| Behaviourally eligible sample (n = 39) | Behavioural report information | Permutation test: 6.2 ms | MI(location accuracy; expression accuracy) | Observed = 0.0027 bits; Excess over null = 0.0016 bits; Holm p = 0.0015 |
| Behaviourally eligible sample (n = 39) | Behavioural report information | Permutation test: 6.2 ms | MI(PAS; joint accuracy) | Observed = 0.0612 bits; Excess over null = 0.0508 bits; Holm p = 0.0015 |
| Behaviourally eligible sample (n = 39) | Behavioural report information | Permutation test: 6.2 ms | CMI(PAS; location accuracy \| expression accuracy) | Observed = 0.0305 bits; Excess over null = 0.0235 bits; Holm p = 0.0015 |
| Behaviourally eligible sample (n = 39) | Behavioural report information | Permutation test: 6.2 ms | CMI(PAS; expression accuracy \| location accuracy) | Observed = 0.0316 bits; Excess over null = 0.0260 bits; Holm p = 0.0015 |
| Behaviourally eligible sample (n = 39) | Behavioural report information | Permutation test: 6.2 ms | Co-information(PAS; location accuracy; expression accuracy) | Observed = -0.0008 bits; Excess over null = 0.0012 bits; Holm p = 0.0015 |
| Behaviourally eligible sample (n = 39) | Behavioural report information | Permutation test: 6.2 ms | Entropy of location accuracy | Observed = 0.2351 bits; Excess over null = 0.0000 bits |
| Behaviourally eligible sample (n = 39) | Behavioural report information | Permutation test: 6.2 ms | Entropy of expression accuracy | Observed = 0.9032 bits; Excess over null = -0.0000 bits |
| Behaviourally eligible sample (n = 39) | Behavioural report information | Permutation test: 6.2 ms | Entropy of joint accuracy | Observed = 1.1356 bits; Excess over null = 0.0000 bits |
| Behaviourally eligible sample (n = 39) | Behavioural report information | Permutation test: 6.2 ms | Normalised MI(PAS; location accuracy) | Observed = 0.1360 bits; Excess over null = 0.1015 bits; Holm p = 0.0015 |

| | | | | |
|---|---|---|---|---|
| Behaviourally eligible sample (n = 39) | Behavioural report information | Permutation test: 6.2 ms | Normalised MI(PAS; expression accuracy) | Observed = 0.0350 bits; Excess over null = 0.0310 bits; Holm p = 0.0015 |
| Behaviourally eligible sample (n = 39) | Behavioural report information | Permutation test: 6.2 ms | Normalised MI(PAS; joint accuracy) | Observed = 0.0546 bits; Excess over null = 0.0452 bits; Holm p = 0.0015 |
| Behaviourally eligible sample (n = 39) | Behavioural report information | Permutation test: 6.2 ms | CMI(PAS; location report \| stimuli) | Observed = 0.0574 bits; Excess over null = 0.0239 bits; Holm p = 0.0015 |
| Behaviourally eligible sample (n = 39) | Behavioural report information | Permutation test: 6.2 ms | CMI(PAS; expression report \| stimuli) | Observed = 0.0648 bits; Excess over null = 0.0417 bits; Holm p = 0.0015 |
| Behaviourally eligible sample (n = 39) | Behavioural report information | Permutation test: 6.2 ms | CMI(location report; expression report \| stimuli) | Observed = 0.0255 bits; Excess over null = 0.0138 bits; Holm p = 0.0015 |
| Behaviourally eligible sample (n = 39) | Behavioural report information | Permutation test: 6.2 ms | CMI(PAS; joint reports \| stimuli) | Observed = 0.1268 bits; Excess over null = 0.0659 bits; Holm p = 0.0015 |
| Behaviourally eligible sample (n = 39) | Behavioural report information | Permutation test: 6.2 ms | Conditional co-information among reports given stimuli | Observed = -0.0046 bits; Excess over null = -0.0004 bits; Holm p = 0.4128 |

**Supplementary Table S7 | P1-LPP transfer-entropy robustness analyses.** The table reports planned and sensitivity-map results, participant-influence summaries, and leave-one-participant-out estimates.

**A. Map-level robustness tests**

| Robustness analysis | Test | Minimum cluster p | Holm-adjusted p | Significant after correction | Cluster direction | Early interval (ms) | Late interval (ms) |
|---|---|---|---|---|---|---|---|
| Planned direction test | P1 → LPP: 6.2 ms – 4.4 ms | 0.8228 | 1.0000 | No | | | |
| Planned direction test | LPP → P1: 6.2 ms – 4.4 ms | 0.0036 | 0.0108 | Yes | Positive | 88–112 | 440–500 |
| Planned direction test | Direction interaction: 6.2 ms – 4.4 ms | 0.6051 | 1.0000 | No | | | |
| Extended late-window boundary | LPP → P1, extended late window: 6.2 ms – 4.4 ms | 0.0026 | 0.0026 | Yes | Positive | 84–112 | 440–504 |
| Signal-representation sensitivity | CSD, published sensor nodes: 6.2 ms – 4.4 ms | 0.2825 | 0.5595 | No | | | |
| Signal-representation sensitivity | Voltage, ROI sensor nodes: 6.2 ms – 4.4 ms | 0.2797 | 0.5595 | No | | | |

| | | | | | | | |
|---|---|---|---|---|---|---|---|
| Signal-representation sensitivity | Voltage, published sensor nodes: 6.2 ms – 4.4 ms | 0.1340 | 0.4019 | No | | | |
| Expression-specific sensitivity | Fearful trials: 6.2 ms – 4.4 ms | 0.0220 | 0.0660 | No | | | |
| Expression-specific sensitivity | Happy trials: 6.2 ms – 4.4 ms | 0.4931 | 0.4931 | No | | | |
| Expression-specific sensitivity | Fearful minus happy: 6.2 ms – 4.4 ms | 0.1086 | 0.2172 | No | | | |
| Raw-estimator reproduction | LPP → P1, raw estimator: 6.2 ms – 4.4 ms | 0.0004 | 0.0004 | Yes | Positive | 88–136 | 436–500 |

**B. Participant-influence summary**

| Estimator | Mean effect (bits) | SEM (bits) | Cohen's $d_z$ | Participants with positive effect (%) | Leave-one-out minimum (bits) | Leave-one-out maximum (bits) |
|---|---|---|---|---|---|---|
| Raw transfer entropy | 0.006 | 0.001 | 0.942 | 84 | 0.0054082 | 0.0062856 |
| Surrogate-centred transfer entropy | 0.006 | 0.001 | 0.887 | 78 | 0.0053759 | 0.0062655 |

**Supplementary Table S8 | EEG cluster and map results.** The table collates ERP, multivariate decoding, representational-similarity, and sensor-network tests. Probability values retain the correction field produced by the corresponding planned analysis family. Additional analyses are described in the Supplementary Results.

| Analysis | Test or contrast | Direction | Time interval(s) (ms) | Cluster mass | Cluster p | Adjusted p | Channels, n |
|---|---|---|---|---|---|---|---|
| ERP: exposure duration | Exposure duration: 1.7 vs 4.4 vs 6.2 ms | Positive | 48-596 | 262104.923 | 0.0001 | | 64 |
| ERP: exposure duration | 1.7 - 4.4 ms | Positive | 272-596 | 8205.824 | 0.0002 | | 36 |
| ERP: exposure duration | 1.7 - 4.4 ms | Positive | 48-264 | 6013.923 | 0.0004 | | 63 |
| ERP: exposure duration | 1.7 - 4.4 ms | Negative | 152-596 | -10745.989 | 0.0002 | | 58 |
| ERP: exposure duration | 1.7 - 4.4 ms | Negative | 52-148 | -2131.577 | 0.0254 | | 27 |
| ERP: exposure duration | 4.4 - 6.2 ms | Positive | 244-596 | 7357.506 | 0.0002 | | 36 |
| ERP: exposure duration | 4.4 - 6.2 ms | Positive | 64-200 | 2972.837 | 0.0062 | | 51 |

| | | | | | | | |
|---|---|---|---|---|---|---|---|
| ERP: exposure duration | 4.4 - 6.2 ms | Negative | 64-596 | -12569.052 | 0.0002 | | 47 |
| ERP: exposure duration | 1.7 - 6.2 ms | Positive | 44-596 | 15264.913 | 0.0002 | | 60 |
| ERP: exposure duration | 1.7 - 6.2 ms | Negative | 48-596 | -14279.697 | 0.0002 | | 59 |
| ERP: emotional versus neutral faces | 6.2 ms \| Emotional - neutral | Positive | 284-480 | 1562.413 | 0.0010 | 0.0030 | 31 |
| ERP: emotional versus neutral faces | 6.2 ms \| Emotional - neutral | Positive | 492-596 | 927.541 | 0.0094 | 0.0030 | 23 |
| ERP: emotional versus neutral faces | 6.2 ms \| Emotional - neutral | Negative | 284-436 | -989.862 | 0.0076 | 0.0030 | 19 |
| ERP: emotional versus neutral faces | 6.2 ms \| Emotional - neutral | Negative | 520-596 | -640.987 | 0.0254 | 0.0030 | 20 |
| MVPA: expression and location decoding | Expression decoding generalised across face identities; 6.2 ms | Positive | 348-596 | 245.51 | 0.0002 | 0.0044 | |
| MVPA: expression and location decoding | Location decoding generalised across face identities; 4.4 ms | Positive | 0-596 | 1023.045 | 0.0002 | 0.0008 | |
| MVPA: expression and location decoding | Location decoding generalised across face identities; 6.2 ms | Positive | 0-596 | 1275.936 | 0.0002 | 0.0008 | |
| MVPA: PAS modulation of neural evidence | PAS slope for location evidence; 4.4 ms | Positive | 0-208 | 150.809 | 0.0008 | 0.0048 | |
| MVPA: PAS modulation of neural evidence | PAS slope for location evidence; 4.4 ms | Positive | 240-596 | 413.477 | 0.0002 | 0.0016 | |
| MVPA: PAS modulation of neural evidence | PAS slope for location evidence; 6.2 ms | Positive | 312-452 | 110.552 | 0.0006 | 0.0042 | |
| MVPA: follow-up analyses | Expression minus location evidence; 4.4 ms | Negative | 248-596 | 334.976 | 0.0002 | 0.0018 | |
| Information theory: EPN-LPP network | Co-information: 1.7 ms | Positive | | 360.51 | 0.0072 | 0.0144 | |
| Information theory: EPN-LPP network | Co-information: 4.4 ms | Positive | | 287.196 | 0.0164 | 0.0492 | |
| Information theory: EPN-LPP network | Co-information: 4.4 ms | Positive | | 463.24 | 0.0008 | 0.0024 | |
| Information theory: PAS and network dynamics | PAS mutual information: posterior 4.4 ms | Positive | 180-320 | 17.559 | 0.0006 | 0.0036 | |

| | | | | | | |
|---|---|---|---|---|---|---|
| Information theory: PAS and network dynamics | PAS mutual information: posterior 4.4 ms | Positive | 440-520 | 10.744 | 0.0066 | 0.0396 |
| Information theory: PAS and network dynamics | PAS mutual information: posterior 6.2 ms | Positive | 260-380 | 16.736 | 0.0018 | 0.0080 |
| Information theory: PAS and network dynamics | PAS mutual information: posterior 6.2 ms | Positive | 420-480 | 9.624 | 0.0098 | 0.0392 |
| Information theory: PAS and network dynamics | PAS mutual information: posterior 6.2 ms - 1.7 ms | Positive | 420-480 | 10.888 | 0.0022 | 0.0132 |
| Information theory: PAS and network dynamics | PAS mutual information: centroparietal 1.7 ms | Positive | 420-480 | 7.959 | 0.0148 | 0.0296 |
| Information theory: PAS and network dynamics | PAS mutual information: centroparietal 4.4 ms | Positive | 500-596 | 14.511 | 0.0022 | 0.0080 |
| Information theory: PAS and network dynamics | PAS mutual information: centroparietal 6.2 ms | Positive | 420-540 | 17.133 | 0.0016 | 0.0080 |
| Information theory: P1-EPN network | Mutual information: EPN 6.2 ms | Positive | | 19.122 | 0.0006 | 0.0036 |
| Information theory: P1-EPN network | Mutual information: EPN 6.2 ms - 1.7 ms | Positive | | 15.469 | 0.0050 | 0.0300 |
| Information theory: P1-LPP network | Transfer entropy: LPP to P1 6.2 ms - 4.4 ms | Positive | | 243.302 | 0.0010 | 0.0090 |
| MVPA: expression and location decoding | Identity-generalising expression decoding: 6.2–4.4 ms | Positive | 288–324 | 26.720 | .0236 | |
| MVPA: expression and location decoding | Identity-generalising expression decoding: 6.2–4.4 ms | Positive | 440–472 | 22.909 | .0402 | |

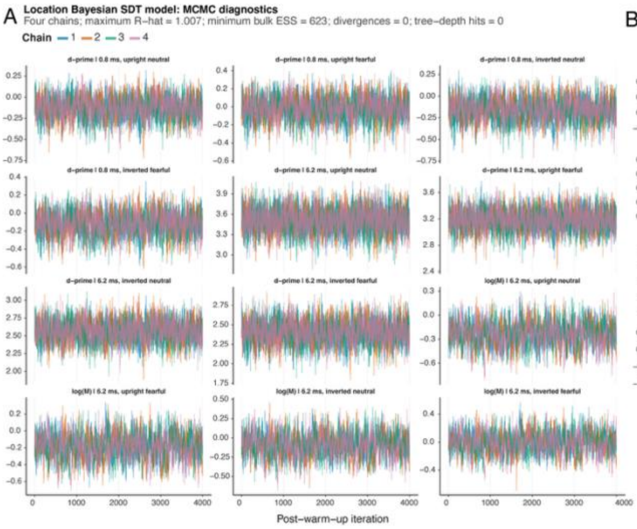


**Supplementary Figure S1. MCMC diagnostics for the Bayesian hierarchical signal-detection models in Experiment 1.** Trace plots show selected group-level parameters from the four post-warm-up Markov chains, with 4,000 retained iterations per chain, for the 31 participants included in the behavioural models. **A**, Location model. Traces are shown for type-1 sensitivity, d', at the shortest and longest exposure durations for upright-neutral, upright-fearful, inverted-neutral, and inverted-fearful faces, together with log metacognitive efficiency, log(M), at 6.2 ms for the same conditions. **B**, Expression model. Traces are shown for d' and log(M) at 0.8 and 6.2 ms for upright and inverted faces. Chain colours identify the four independently initialised chains. The diagnostic summaries above each panel report the worst value across all monitored parameters, not only those displayed. For the location model, maximum $\hat{R}$ was 1.007 and minimum bulk effective sample size was 623; for the expression model, maximum $\hat{R}$ was 1.003 and minimum bulk effective sample size was 1,126. Neither model produced divergent transitions or maximum-tree-depth hits.

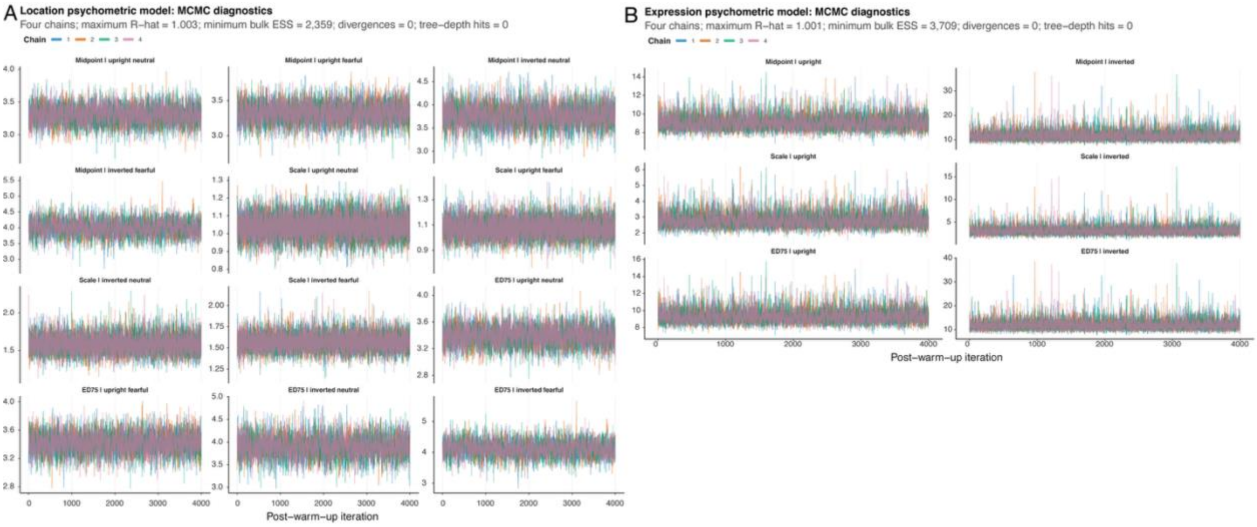


**Supplementary Figure S2. MCMC diagnostics for the hierarchical psychometric-function models in Experiment 1.** Trace plots show condition-level parameters from the four post-warm-up Markov chains, with 4,000 retained iterations per chain, for the 31 participants included in the behavioural models. **A**, Location model. Traces are shown for the midpoint, scale, and $ED_{75}$ of the cumulative-Gaussian psychometric function for upright-neutral, upright-fearful, inverted-neutral, and inverted-fearful faces. **B**, Expression model. The corresponding parameters are shown for upright and inverted faces. $ED_{75}$ denotes the exposure duration at which expected accuracy reached .75 after accounting for the lapse parameter. Chain colours identify the four independently initialised chains. The summaries above each panel report the worst diagnostic value across all monitored parameters. For the location model, maximum $\hat{R}$ was 1.003 and minimum bulk effective sample size was 2,359; for the expression model, maximum R̂ was 1.001 and minimum bulk effective sample size was 3,709. Neither model produced divergent transitions or maximum-tree-depth hits.

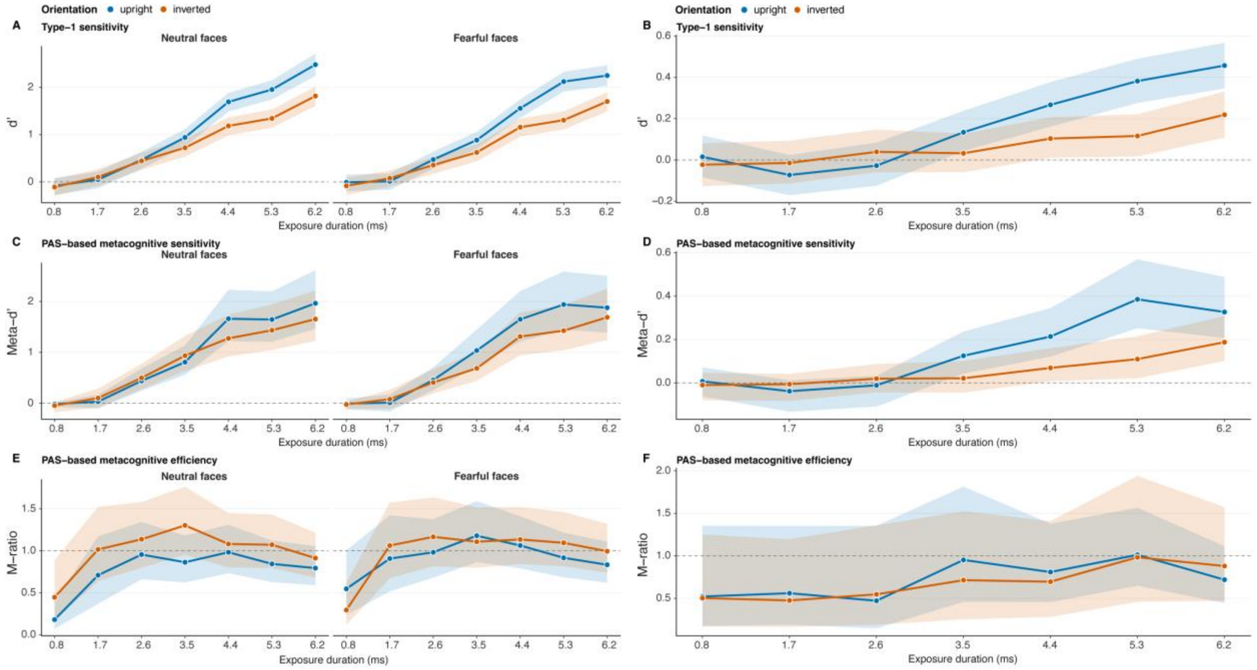


**Supplementary Figure S3. Bayesian signal-detection estimates for Experiment 1.** Posterior estimates from the hierarchical signal-detection models are shown across the seven exposure durations for upright faces (blue) and inverted faces (orange). **A, B**, Type-1 sensitivity, d', for location and expression decisions, respectively. **C, D**, PAS-based metacognitive sensitivity, meta-d', for location and expression decisions. **E, F**, PAS-based metacognitive efficiency, expressed as the M-ratio, for location and expression decisions. Location estimates are shown separately for neutral and fearful faces; expression estimates combine the two presented expressions within each orientation condition. Points and lines denote posterior medians, and shaded regions denote 95% credible intervals across the 31 included participants. The dashed horizontal lines indicate zero sensitivity in panels A–D and an M-ratio of 1 in panels E and F. Location d' and meta-d' were divided by $\sqrt{2}$ to express sensitivity under the two-alternative forced-choice convention used in the original study.

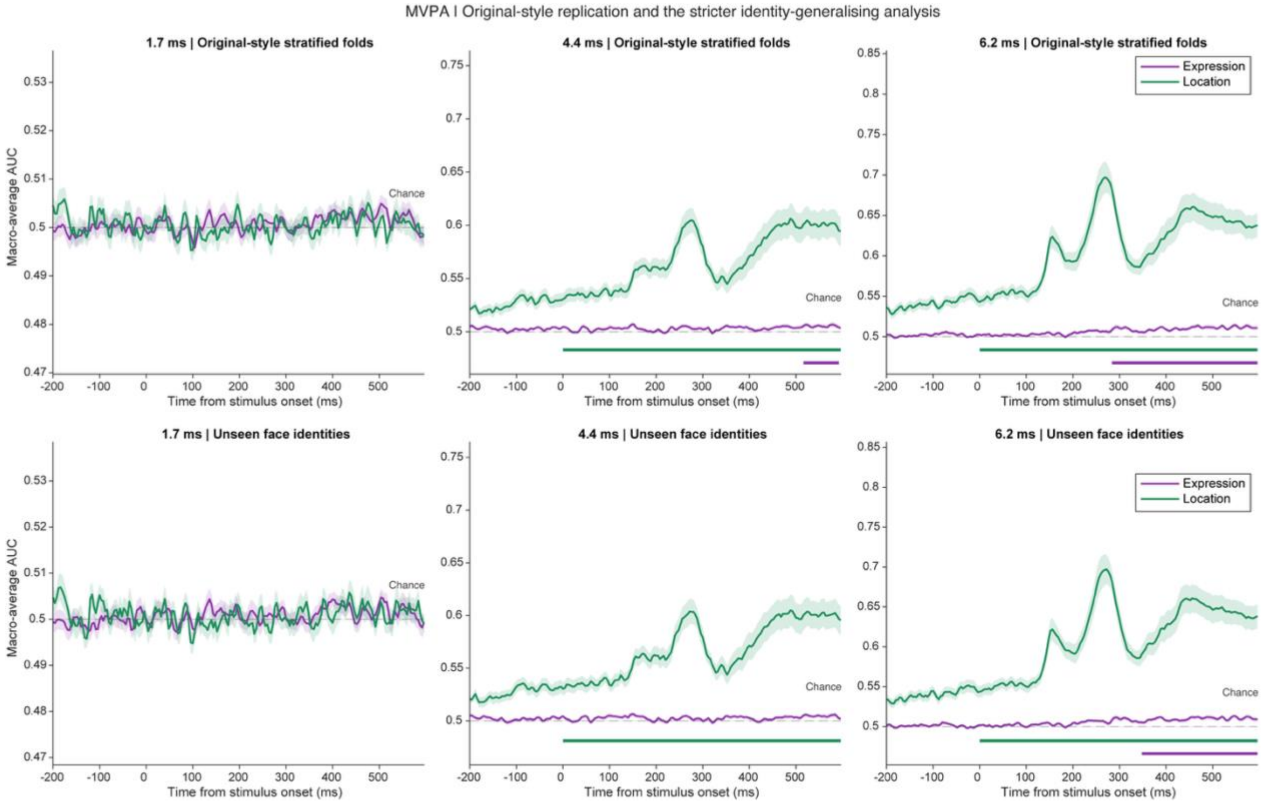


**Supplementary Figure S4. Original-style and identity-generalising EEG decoding.** Time-resolved macro-averaged area under the receiver-operating-characteristic curve (AUC) for facial-expression classification (purple) and intact-face location classification (green) at exposure durations of 1.7, 4.4, and 6.2 ms. The upper row shows repeated trial-stratified cross-validation, approximating the original analysis, in which trials depicting the same face could occur in both the training and test sets. The lower row shows the stricter identity-grouped analysis, in which all trials depicting a given face were assigned to the same fold, requiring classification to generalise to previously unseen faces. Curves show the participant mean and shaded regions indicate ±1 SEM across the 32 participants included in the EEG analyses. The dashed horizontal line denotes chance performance, AUC = .50. Purple and green horizontal bars indicate expression and location decoding, respectively, in temporal clusters surviving the cluster-permutation procedure at the familywise threshold of α = .05. The vertical scale differs across exposure-duration columns.

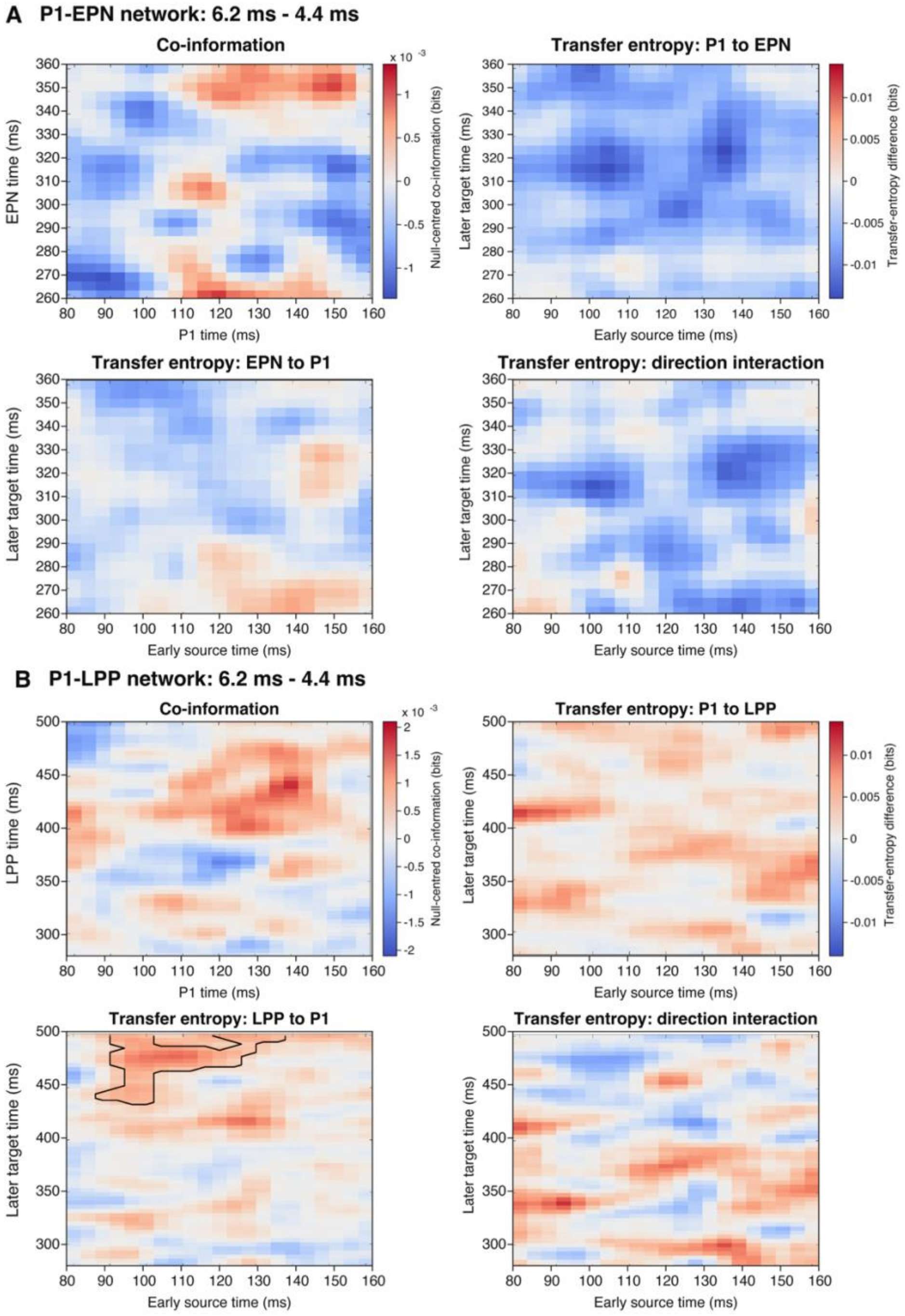


**Supplementary Figure S5. Exposure-dependent P1–EPN and P1–LPP information dynamics.** Information-theoretic contrasts between the 6.2-ms and 4.4-ms exposure conditions for the P1–EPN network (**A**) and P1–LPP network (**B**) in the 32 participants included in the EEG analyses. Analyses used scalp-current-density signals and the planned broader component windows: 80–160 ms for P1, 260–360 ms for EPN, and 280–500 ms for LPP. Within each panel, the upper-left map shows the 6.2-ms minus 4.4-ms change in signed, permutation-null-centred co-information about emotional versus neutral status. Positive co-information contrasts indicate a shift towards a more redundancy-dominant balance at 6.2 ms, whereas negative contrasts indicate a shift towards a more synergy-dominant balance. The upper-right and lower-left maps show the corresponding exposure-duration contrast in the emotional-minus-neutral transfer-entropy effect for the P1-to-target and target-to-P1 directions, respectively. The lower-right map shows the direction interaction, defined as the P1-to-target emotional-minus-neutral effect minus the target-to-P1 emotional-minus-neutral effect, contrasted between 6.2 and 4.4 ms. Warm colours denote positive 6.2-ms minus 4.4-ms differences, and cool colours denote negative differences. Black contours identify clusters surviving maximum-cluster-mass correction within the map and Holm correction across the planned maps in the corresponding inferential family. Colour variation outside the outlined clusters is descriptive.
